# The human language system, including its inferior frontal component in ‘Broca’s area’, does not support music perception

**DOI:** 10.1101/2021.06.01.446439

**Authors:** Xuanyi Chen, Josef Affourtit, Rachel Ryskin, Tamar I. Regev, Samuel Norman-Haignere, Olessia Jouravlev, Saima Malik-Moraleda, Hope Kean, Rosemary Varley, Evelina Fedorenko

**Author notes:** Corresponding Authors and; 43 Vassar Street, Room 46-3037, Cambridge, MA, 02139. Co-senior authors (Rosemary Varley, Evelina Fedorenko).

## Abstract

Language and music are two human-unique capacities whose relationship remains debated. Some have argued for overlap in processing mechanisms, especially for structure processing. Such claims often concern the inferior frontal component of the language system located within ‘Broca’s area’. However, others have failed to find overlap. Using a robust individual-subject fMRI approach, we examined the responses of language brain regions to music stimuli, and probed the musical abilities of individuals with severe aphasia. Across four experiments, we obtained a clear answer: music perception does not engage the language system, and judgments about music structure are possible even in the presence of severe damage to the language network. In particular, the language regions’ responses to music are generally low, often below the fixation baseline, and never exceed responses elicited by non-music auditory conditions, like animal sounds. Further, the language regions are not sensitive to music structure: they show low responses to intact and structure-scrambled music, and to melodies with vs. without structural violations. Finally, in line with past patient investigations, individuals with aphasia who cannot judge sentence grammaticality perform well on melody well-formedness judgments. Thus the mechanisms that process structure in language do not appear to process music, including music syntax.

## Introduction

To interpret language or appreciate music, we must understand how different elements—words in language, notes and chords in music—relate to each other. Parallels between the structural properties of language and music have been drawn for over a century (e.g., Riemann 1877, as cited in Swain 1995; Lindblom and Sundberg 1969; Fay 1971; Boiles 1973; Cooper 1973; Bernstein 1976; Sundberg and Lindblom 1976; Lerdahl and Jackendoff 1977, 1983; Roads and Wieneke 1979; Krumhansl and Keil 1982; Baroni et al. 1983; Swain 1995; cf. Jackendoff 2009; Temperley 2022). However, the question of whether music processing relies on the same mechanisms as those that support language processing continues to spark debate.

The empirical landscape is complex. A large number of studies have argued for overlap in structural processing based on behavioral (e.g., Fedorenko et al. 2009; Slevc et al. 2009; Hoch et al. 2011; Van de Cavey and Hartsuiker 2016; Kunert et al. 2016), ERP (e.g., Janata 1995; Patel et al. 1998; Koelsch et al. 2000), MEG (e.g., Maess et al. 2001), fMRI (e.g., Koelsch et al. 2002; Levitin and Menon 2003; Tillmann et al. 2003; Koelsch 2006; Kunert et al. 2015; Musso et al. 2015), and ECoG (e.g., Sammler et al. 2009, 2013; Rietmolen et al. 2022) evidence (see Tillman 2012; Kunert and Slevc 2015; LaCroix et al. 2016, for reviews). However, we would argue that no prior study has compellingly established reliance on shared syntactic processing mechanisms in language and music.

*First*, evidence from behavioral, ERP, and, to a large extent, MEG studies is indirect because they do not allow to unambiguously determine where neural responses originate (in ERP and MEG, this is due to the ‘inverse problem’; Tarantola 2004; Baillet et al. 2014).

*Second*, the bulk of the evidence comes from structure-violation paradigms. In such paradigms, responses to the critical condition—which contains an element that violates the rules of tonal music—are contrasted with responses to the control condition, where stimuli obey the rules of tonal music. (For language, syntactic violations, like violations of number agreement, are often used.) Because structural violations (across domains) constitute unexpected events, a brain region that responds more strongly to the structure-violation condition than the control (no violation) condition *may* support structure processing in music, but it may also reflect domain-general processes, like attention or error detection/correction (e.g., Bigand et al. 2001; Poulin-Charronat et al. 2005; Tillmann et al. 2006; Hoch et al. 2011; Perruchet and Poulin-Charronnat 2013) or low-level sensory effects (e.g., Bigand et al. 2014; Collins et al. 2014; cf. Koelsch et al. 2007). In order to argue that a brain region that shows a *structure-violation > no violation* effect supports structure processing in music, one would need to establish that this brain region i) is selective for structural violations and does not respond to unexpected non-structural (but similarly salient) events in music or other domains, and ii) responds to music stimuli even when no violation is present. This latter point is (surprisingly) not often discussed but is deeply important: if a brain region supports the processing of music structure, it should be engaged whenever music is processed (similar to how language areas respond robustly to well-formed sentences, in addition to showing sensitivity to violated linguistic expectations; e.g., Fedorenko et al. 2020). After all, in order to detect a structural violation, a brain region needs to process the structure of the preceding context, which implies that it should be working whenever a music stimulus is present. No previous study has established both of the properties above—selectivity for structural relative to non-structural violations and robust responses to music stimuli with no violations—for the brain regions that have been argued to support structure processing in music (and to overlap with regions that support structure processing in language). In fact, some studies that have compared unexpected structural and non-structural events in music (e.g., a timbre change) have reported similar neural responses in fMRI (e.g., Koelsch et al. 2002; cf. some differences in EEG effects – e.g., Koelsch et al. 2001). Relatedly, and in support of the idea that effects of music structure violations largely reflect domain-general attentional effects, meta-analyses of neural responses to unexpected events across domains (e.g., Corbetta and Shulman 2002; Fouragnan et al. 2018; Corlett et al. 2021) have identified regions that grossly resemble those reported in studies of music structure violations (see Fedorenko and Varley 2016 for discussion).

*Third*, most prior fMRI (and MEG) investigations have relied on comparisons of group-level activation maps. Such analyses suffer from low functional resolution (e.g., Nieto-Castañón and Fedorenko 2012; Fedorenko 2021), especially in cases where the precise locations of functional regions vary across individuals, as in the association cortex (Fischl et al. 2008; Frost and Goebel 2012; Tahmasebi et al. 2012; Vazquez-Rodriguez et al. 2019). Thus, observing activation overlap at the group level does not unequivocally support shared mechanisms. Indeed, studies that have used individual-subject-level analyses have reported a low or no response to music in the language-responsive regions (Fedorenko et al. 2011; Rogalsky et al. 2011; Deen et al. 2015).

*Fourth*, the interpretation of some of the observed effects has relied on the so-called ‘reverse inference’ (Poldrack 2006, 2011; Fedorenko 2021), where function is inferred from a coarse anatomical location: for example, some music-structure-related effects observed in or around ‘Broca’s area’ have been interpreted as reflecting the engagement of linguistic-structure-processing mechanisms (e.g., Maess et al. 2001; Koelsch et al. 2002) given the long-standing association between ‘Broca’s area’ and language, including syntactic processing specifically (e.g., Caramazza and Zurif 1976; Friederici et al. 2006). However, this reasoning is not valid: Broca’s area is a heterogeneous region, which houses components of at least two functionally distinct brain networks (Fedorenko et al. 2012; Fedorenko and Blank 2020): the language-selective network, which responds during language processing, visual or auditory, but does not respond to diverse non-linguistic stimuli (Fedorenko et al. 2011; Monti et al. 2009, 2012; see Fedorenko and Varley 2016 for a review) and the domain-general executive control or ‘multiple demand (MD)’ network, which responds to any demanding cognitive task and is robustly modulated by task difficulty (Duncan 2010, 2013; Fedorenko et al. 2013; Assem et al. 2020). As a result, here and more generally, functional interpretation based on coarse anatomical localization is not justified.

*Fifth*, many prior fMRI investigations have not reported the magnitudes of response to the relevant conditions and only examined statistical significance maps for the contrast of interest (e.g., a whole brain map showing voxels that respond reliably more strongly to melodies with vs. without a structural violation, and to sentences with vs. without a structural violation). Response magnitudes of experimental conditions relative to a low-level baseline and to each other are critical for interpreting a functional profile of a brain region (see e.g., Chen et al. 2017, for discussion). For example, a reliable *violation > no violation* effect in music (similar arguments apply to language) could be observed when both conditions elicit above-baseline responses, and the violation condition elicits a stronger response (**Figure 1A** left bar graph)—a reasonable profile for a brain region that supports music processing and is sensitive to the target structural manipulation. However, a reliable *violation > no violation* effect could also be observed when both conditions elicit below-baseline responses, and the violation condition elicits a less negative response (**Figure 1A** middle bar graph), or when both conditions elicit low responses—in the presence of a strong response to stimuli in other domains—and the between-condition difference is small (**Figure 1A** right bar graph; note that with sufficient power even very small effects can be highly reliable, but this does not make them theoretically meaningful; e.g., Cumming 2012; Sullivan and Feinn, 2012). The two latter profiles, where a brain region is more active during silence than when listening to music, or when the response is overall low and the effect of interest is minuscule, would be harder to reconcile with a role of this brain region in music processing (see also the second point above).

**Figure 1:**
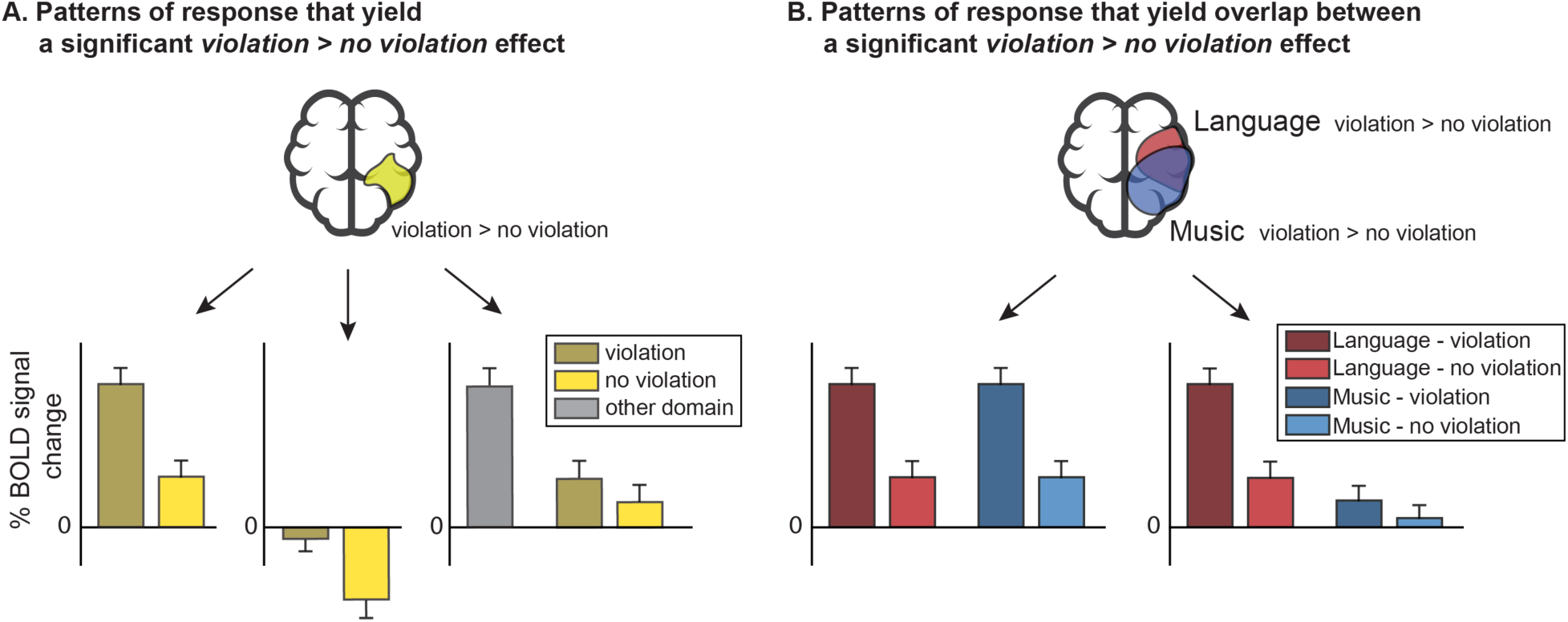
Illustration of the importance of examining the magnitudes of neural response to the experimental conditions rather than only the statistical significance maps for the contrast(s) of interest. A significant *violation > no violation* effect (A) and overlap between a significant *violation > no violation* effect in language vs. in music (B) are each compatible with multiple distinct functional profiles, only one of which (on the left in each case) supports the typically proposed interpretation (a region that processes structure in some domain of interest in A, and a region that processes structure across domains, in both language and music, in B).

Similarly, with respect to the music-language overlap question, a reliable *violation > no violation* effect for both language and music could be observed in a brain region where sentences and melodies with violations elicit similarly strong responses, and those without violations elicit lower responses (**Figure 1B** left bar graph); but it could also arise in a brain region where sentences with violations elicit a strong response, sentences without violations elicit a lower response, but melodies elicit an overall low response, with the violation condition eliciting a higher response than the no-violation condition (**Figure 1B** right bar graph). Whereas in the first case, it may be reasonable to argue that the brain region in question supports some computation that is necessary to process structure violations in both domains, such interpretation would not be straightforward in the second case. In particular, given the large main effect of language > music, any account of possible computations supported by such a brain region would need to explain this difference instead of simply focusing on the presence of a reliable effect of violation in both domains. In summary, without examining the magnitudes of response, it is not possible to distinguish among many, potentially very different, functional profiles, without which formulating hypotheses about a brain region’s computations is precarious.

Aside from the limitations above, to the best of our knowledge, all prior brain imaging studies have used a single manipulation in one set of materials and one set of participants. To compellingly argue that a brain region supports (some aspects of) structural processing in both language and music, it is important to establish both the *robustness* of the key effect by replicating it with a new set of experimental materials and/or in a new group of participants, and its *generalizability* to other contrasts between conditions that engage the hypothesized computation and ones that do not. For example, to argue that a brain region houses a core syntactic mechanism needed to process hierarchical relations and/or recursion in both language and music (e.g., Patel 2003; Fadiga et al. 2009; Roberts 2012; Koelsch et al. 2013; Fitch and Martins 2014), one would need to demonstrate that this region i) responds robustly to diverse structured linguistic and musical stimuli (which all invoke the hypothesized shared computation), ii) shows replicable responses across materials and participants, and iii) is sensitive to more than a single manipulation targeting the hypothesized computations specifically, as needed to rule out paradigm-/task-specific accounts (e.g., structured vs. unstructured stimuli, stimuli with vs. without structural violations, stimuli that are more vs. less structurally complex—e.g., with long-distance vs. local dependencies, adaptation to structure vs. some other aspect of the stimulus, etc.).

Finally, the neuropsychological patient evidence is at odds with the idea of shared mechanisms for processing language and music. If language and music relied on the same syntactic processing mechanism, individuals impaired in their processing of linguistic syntax should also exhibit impairments in musical syntax. Although some prior studies report subtle musical deficits in patients with aphasia (Patel et al. 2008a; Sammler et al. 2011), the evidence is equivocal, and many aphasic patients appear to have little or no difficulties with music, including the processing of music structure (Luria et al. 1965; Brust 1980; Marin 1982; Basso and Capitani 1985; Polk and Kertesz 1993; Slevc et al. 2016; Faroqi-Shah et al. 2020; Chiapetta et al. 2022; cf. Omigie and Samson 2014 and Sihvonen et al. 2017 for discussions of evidence that musical training may lead to better outcomes following brain damage/resection). Similarly, children with Specific Language Impairment (now called Developmental Language Disorder)—a developmental disorder that affects several aspects of linguistic and cognitive processing, including syntactic processing (e.g., Bortolini et al. 1998; Bishop and Norbury 2002)—show no impairments in musical processing (Fancourt 2013; cf. Jentschke et al. 2008). In an attempt to reconcile the evidence from acquired and developmental disorders with claims about structure-processing overlap based on behavioral and neural evidence from neurotypical participants, Patel (2003, 2008, 2012; see Slevc and Okada 2015, Patel and Morgan 2017, and Asano et al. 2021 for related proposals) put forward a hypothesis whereby the representations that mediate language and music are stored in distinct brain areas, but the mechanisms that perform online computations on those representations are partially overlapping. We return to this idea in the Discussion.

To bring clarity to this ongoing debate, we conducted three fMRI experiments with neurotypical adults, and a behavioral study with individuals with severe aphasia. For the fMRI experiments, we took an approach where we focused on the ‘language network’—a well-characterized set of left frontal and temporal brain areas that selectively support linguistic processing (e.g., Fedorenko et al. 2011) and asked whether any parts of this network show responses to music and sensitivity to music structure. In each experiment, we used an extensively validated language ‘localizer’ task based on the reading of sentences and nonword sequences (Fedorenko et al. 2010; see Scott et al. 2017 and Malik-Moraleda, Ayyash et al. 2022 for evidence that this localizer is modality-independent) to identify language-responsive areas in each participant individually. Importantly, these areas have been shown, across dozens of brain imaging studies, to be robustly sensitive to linguistic syntactic processing demands in diverse manipulations (e.g., Keller et al. 2001; Röder et al. 2002; Friederici 2011; Pallier et al. 2011; Bautista and Wilson 2016, among many others)—including when defined with the same localizer as the one used here (e.g., Fedorenko et al. 2010, 2012a, 2020; Blank et al. 2016; Mollica et al. 2020; Shain, Blank et al. 2020; Shain et al. 2022)—and their damage leads to linguistic, including syntactic, deficits (e.g., Caplan et al. 1996; Dick et al. 2001; Wilson and Saygin 2004; Tyler et al. 2011; Wilson et al. 2012; Mesulam et al. 2014; Ding et al. 2020; Matchin and Hickok 2020, among many others). To address the critical research question, we examined the responses of these language areas to music, and their necessity for processing music structure. In Experiment 1, we included several types of music stimuli including orchestral music, single-instrument music, synthetic drum music, and synthetic melodies, a minimal comparison between songs and spoken lyrics, and a set of non-music auditory control conditions. We additionally examined sensitivity to structure in music across two structure-scrambling manipulations. In Experiment 2, we further probed sensitivity to structure in music using the most common manipulation, contrasting responses to well-formed melodies vs. melodies containing a note that does not obey the constraints of Western tonal music. And in Experiment 3, we examined the ability to discriminate between well-formed melodies and melodies containing a structural violation in three profoundly aphasic individuals across two tasks. Finally, in Experiment 4, we examined the responses of the language regions to yet another set of music stimuli in a new set of participants. Further, the participants were all native speakers of Mandarin, a tonal language, which allowed us to evaluate the hypothesis that language regions may play a greater role in music processing in individuals with higher sensitivity to linguistic pitch (e.g., Deutsch et al. 2006, 2009; Bidelman et al. 2011; Creel et al. 2018; Ngo et al. 2016; Liu et al. 2021).

## Materials and methods

### Participants

#### Experiments 1, 2, and 4 (fMRI)

48 individuals (age 18-51, mean 24.3; 28 female, 20 male) from the Cambridge/Boston, MA community participated for payment across three fMRI experiments (n=18 in Experiment 1; n=20 in Experiment 2; n=18 in Experiment 4; 8 participants overlapped between Experiments 1 and 2). 33 participants were right-handed and four left-handed, as determined by the Edinburgh handedness inventory (Oldfield 1971), or self-report (see Willems et al. 2014, for arguments for including left-handers in cognitive neuroscience research); the handedness data for the remaining 11 participants (one in Experiment 2 and 10 in Experiment 4) were not collected. All but one participant (with no handedness information) in Experiment 4 showed typical left-lateralized language activations in the language localizer task described below (as assessed by numbers of voxels falling within the language parcels in the left vs. right hemisphere (LH vs. RH), using the following formula: (LH-RH)/(LH+RH); e.g., Jouravlev et al. 2020; individuals with values of 0.25 or greater were considered to have a left-lateralized language system). For the participant with right-lateralized language activations (with a lateralization value at or below -0.25), we used right-hemisphere language regions for the analyses (see **SI-3** for analyses where the LH language regions were used for this participant and when this participant is excluded; the critical results were not affected). Participants in Experiments 1 and 2 were native English speakers; participants in Experiment 4 were native Mandarin speakers and proficient speakers of English (none had any knowledge of Russian, which was used in the unfamiliar foreign-language condition in Experiment 4). Detailed information on the participants’ music background was, unfortunately, not collected, except for ensuring that the participants were not professional musicians. All participants gave informed written consent in accordance with the requirements of MIT’s Committee on the Use of Humans as Experimental Subjects (COUHES).

#### Experiment 3 (behavioral)

##### Individuals with aphasia

Three participants with severe and chronic aphasia were recruited to the study (SA, PR, and PP). All participants gave informed consent in accordance with the requirements of UCL’s Institutional Review Board. Background information on each participant is presented in **Table 1**. Anatomical scans are shown in **Figure 2A** and extensive perisylvian damage in the left hemisphere, encompassing areas where language activity is observed in neurotypical individuals, is illustrated in **Figure 2B**.

**Table 1.**
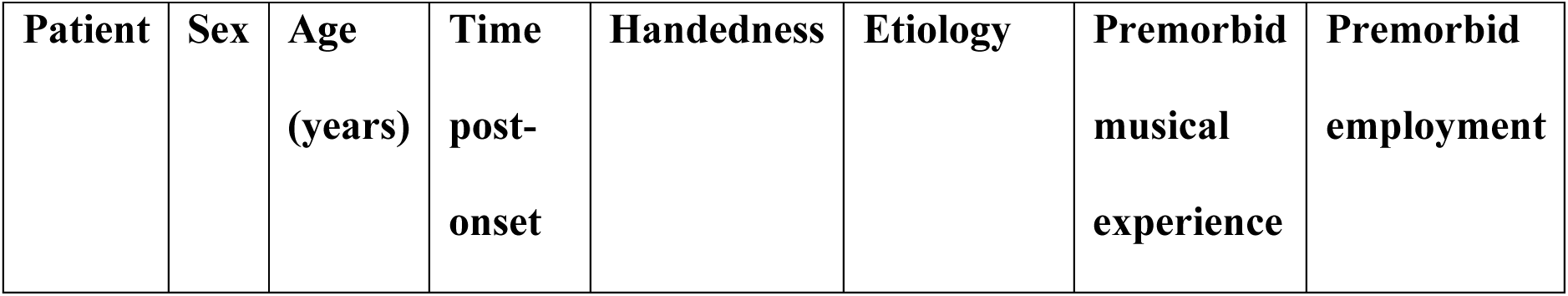
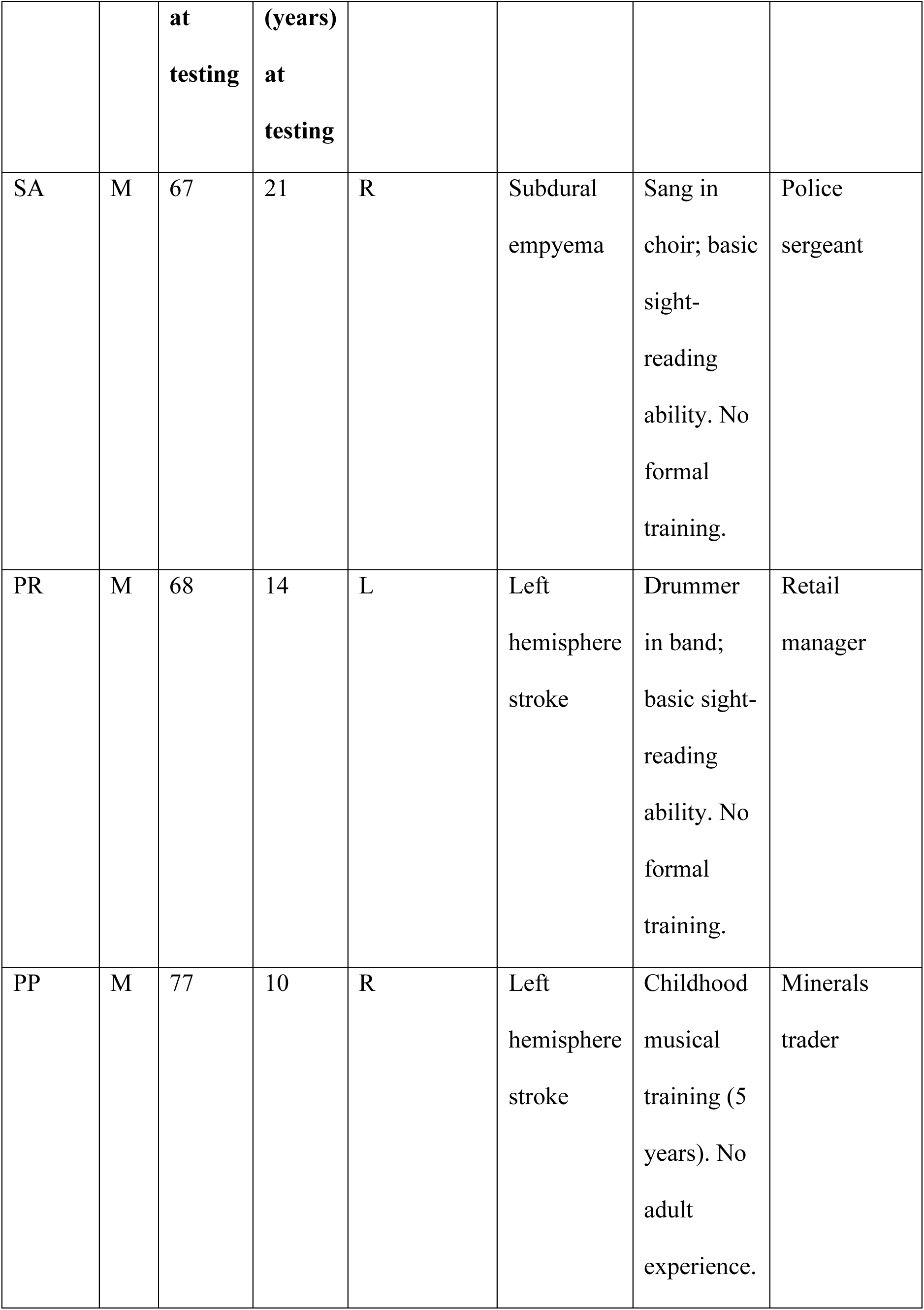
Background information on the participants with aphasia.

**Figure 2:**
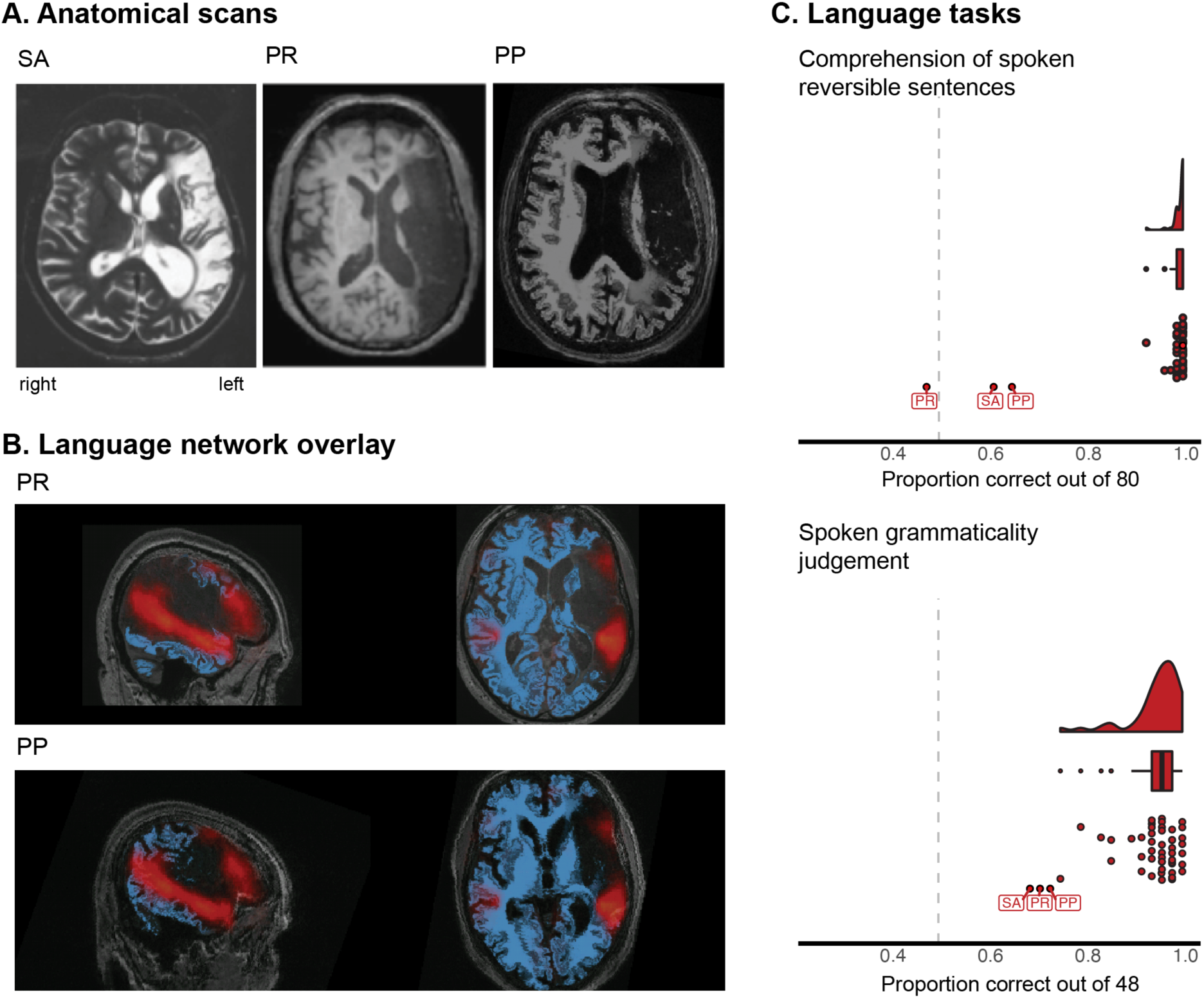
**A.** Anatomical scans (T2-weighted for SA, T1-weighted for PR and PP) of the aphasic participants (all scans were performed during the chronic phase, as can be seen from the ventricular enlargement). Note that the right side of the image represents the left side of the brain. **B.** P.R.’s (top) and P.P.’s (bottom) anatomical scans (blue-tinted) shown with the probabilistic activation overlap map for the fronto-temporal language network overlaid (SA’s raw anatomical data were not available). The map was created by overlaying thresholded individual activation maps (red-tinted) for the *sentences > nonwords* contrast (Fedorenko et al. 2010) in 220 neurotypical participants (none of whom were participants in any experiments in the current study). As the images show, the language network falls largely within the lesioned tissue in the left hemisphere. **C.** Performance of the control participants and participants with aphasia on two measures of linguistic syntax processing (see Design, materials, and procedure – Experiment 3): the comprehension of spoken reversible sentences (top), and the spoken grammaticality judgments (bottom). The densities show the distribution of proportion correct scores in the control participants and the boxplot shows the quartiles of the control population (the whiskers show 1.5x interquartile range and points represent outliers). The dots show individual participants (for the individuals with aphasia, the initials indicate the specific participant). Dashed grey lines indicate chance performance.

##### Control participants

We used Amazon.com’s Mechanical Turk platform to recruit normative samples for the music tasks and a subset of the language tasks that are most critical to linguistic syntactic comprehension. Ample evidence now shows that online experiments yield data that closely mirror the data patterns in experiments conducted in a lab setting (e.g., Crump et al. 2013). Data from participants with IP addresses in the US who self-reported being native English speakers were included in the analyses. 50 participants performed the critical music task, and the Scale task from the MBEA (Peretz et al. 2003), as detailed below. Data from participants who responded incorrectly to the catch trial in the MBEA Scale task (n=5) were excluded from the analyses, for a final sample of 45 control participants for the music tasks. A separate sample of 50 participants performed the *Comprehension of spoken reversible sentences* task. Data from one participant who completed fewer than 75% of the questions and another participant who did not report being a native English speaker were excluded for a final sample of 48 control participants. Finally, a third sample of 50 participants performed the *Spoken grammaticality judgment* task. Data from one participant who did not report being a native English speaker were excluded for a final sample of 49 control participants.

### Design, materials, and procedure

#### Experiments 1, 2, and 4 (fMRI)

Each participant completed a language localizer task (Fedorenko et al. 2010) and one or more of the critical music perception experiments, along with one or more tasks for unrelated studies. The scanning sessions lasted approximately two hours.

##### Language localizer

This task is described in detail in Fedorenko et al. (2010) and subsequent studies from the Fedorenko lab (e.g., Fedorenko et al. 2011; Blank et al. 2014; Blank et al. 2016; Pritchett et al. 2018; Paunov et al. 2019; Fedorenko et al. 2020; Shain, Blank et al. 2020, among others) and is available for download from https://evlab.mit.edu/funcloc/). Briefly, participants read sentences and lists of unconnected, pronounceable nonwords in a blocked design. Stimuli were presented one word/nonword at a time at the rate of 450ms per word/nonword. Participants read the materials passively and performed a simple button-press task at the end of each trial (included in order to help participants remain alert). Each participant completed two ∼6-minute runs. This localizer task has been extensively validated and shown to be robust to changes in the materials, modality of presentation (visual vs. auditory), and task (e.g., Fedorenko et al. 2010; Fedorenko 2014; Scott et al. 2017; Diachek, Blank, Siegelman et al. 2020; Malik-Moraleda, Ayyash et al. 2022; Lipkin et al. 2022; see the results of Experiments 1 and 4 for additional replications of modality robustness). Further, a network that corresponds closely to the localizer contrast (*sentences > nonwords*) emerges robustly from whole-brain task-free data—voxel fluctuations during rest (e.g., Braga et al. 2020; see Braga 2021 for a general discussion of how well-validated localizers tend to show tight correspondence with intrinsic networks recovered in a data-driven way). The fact that different regions of the language network show strong correlations in their activity during naturalistic cognition (see also Blank et al. 2014; Paunov et al. 2019; Malik-Moraleda, Ayyash et al. 2022) provides support for the idea that this network constitutes a ‘natural kind’ in the brain (a subset of the brain that is strongly internally integrated and robustly dissociable from the rest of the brain) and thus a meaningful unit of analysis. However, we also examine individual regions of this network, to paint a more complete picture, given that many past claims about language-music overlap have concerned the inferior frontal component of the language network.

##### Experiment 1

Participants passively listened to diverse stimuli across 18 conditions in a long-event-related design. The materials for this and all other experiments are available at OSF: https://osf.io/68y7c/. All stimuli were 9 s in length. The conditions were selected to probe responses to music, to examine sensitivity to structure scrambling in music, to compare responses to songs vs. spoken lyrics, and to compare responses to music stimuli vs. other auditory stimuli.

The four non-vocal music conditions (all Western tonal music) included orchestral music, single-instrument music, synthetic drum music, and synthetic melodies (see **SI-5** for a summary of the acoustic properties of these and other conditions, as quantified with the MIR toolbox; Lartillot and Toiviainen 2007; Lartillot and Grandjean 2019). The orchestral music condition consisted of 12 stimuli (**SI-Table 4a**) selected from classical orchestras or jazz bands. The single-instrument music condition consisted of 12 stimuli (**SI-Table 4b**) that were played on one of the following instruments: cello (n=1), flute (n=1), guitar (n=4), piano (n=4), sax (n=1), or violin (n=1). The synthetic drum music condition consisted of 12 stimuli synthesized using percussion patches from MIDI files taken from freely available online collections. The stimuli were synthesized using the MIDI toolbox for MATLAB (writemidi). The synthetic melodies condition consisted of 12 stimuli transcribed from folk tunes obtained from freely available online collections. Each melody was defined by a sequence of notes with corresponding pitches and durations. Each note was composed of harmonics 1 through 10 of the fundamental presented in equal amplitude, with no gap in-between notes. Phase discontinuities between notes were avoided by ensuring that the starting phase of the next note was equal to the ending phase of the previous note.

The synthetic drum music and the synthetic melodies conditions had scrambled counterparts to probe sensitivity to music structure. This intact > scrambled contrast has been used in some past studies of structure processing in music (e.g., Levitin and Menon 2003) and is parallel to the sentences > word-list contrast in language, which has been often used to probe sensitivity for combinatorial processing (e.g., Fedorenko et al. 2010). The scrambled drum music condition was created by jittering the inter-note-interval (INI). The amount of jitter was sampled from a uniform distribution (from -0.5 to 0.5 beats). The scrambled INIs were truncated to be no smaller than 5% of the distribution of INIs from the intact drum track. The total distribution of INIs was then scaled up or down to ensure that the total duration remained unchanged. The scrambled melodies condition was created by scrambling both pitch and rhythm information. Pitch information was scrambled by randomly re-ordering the sequence of pitches and then adding jitter to disrupt the key. The amount of jitter for each note was sampled from a uniform distribution centered on the note’s pitch after shuffling (from -3 to +3 semitones). The duration of each note was also jittered (from -0.2 to 0.2 beats). To ensure the total duration was unaffected by jitter, N/2 positive jitter values were sampled, where N is the number of notes, and then a negative jitter was added with the same magnitude for each of the positive samples, such that the sum of all jitters equaled 0. To ensure the duration of each note remained positive, the smallest jitters were added to the notes with the smallest durations. Specifically, the note durations and sampled jitters were sorted by their magnitude, summed, and then the jittered durations were randomly re-ordered.

To allow for a direct comparison between music and linguistic conditions within the same experiment, we included auditory sentences and auditory nonword sequences. The sentence condition consisted of 24 lab-constructed stimuli (half recorded by a male, and half by a female). Each stimulus consisted of a short story (each three sentences long) describing common, everyday events. Any given participant heard 12 of the stimuli (6 male, 6 female). The nonword sequence condition consisted of 12 stimuli (recorded by a male).

We also included two other linguistic conditions: songs and spoken lyrics. These conditions were included to test whether the addition of a melodic contour to speech (in songs) would increase the responses of the language regions. Such a pattern might be expected of a brain region that responds to both linguistic content and music structure. The songs and the lyrics conditions each consisted of 24 stimuli. We selected songs with a tune that was easy to sing without accompaniment. These materials were recorded by four male singers: each recorded between 2 and 11 song-lyrics pairs. The singers were actively performing musicians (e.g., in *a cappella* groups) but were not professionals. Any given participant heard either the song or the lyrics version of an item for 12 stimuli in each condition.

Finally, to assess the specificity of potential responses to music, we included three non-music conditions: animal sounds and two kinds of environmental sounds (pitched and unpitched), which all share some low-level acoustic properties with music (see SI-5). The animal sounds condition and the environmental sounds conditions each consisted of 12 stimuli taken from in-lab collections. If individual recordings were shorter than 9 s, then several recordings of the same type of sound were concatenated together (100 ms gap in between). We included the pitch manipulation in order to test for general responsiveness to pitch—a key component of music—in the language regions.

(The remaining five conditions were not directly relevant to the current study or redundant with other conditions for our research questions and therefore not included in the analyses. These included three distorted speech conditions—lowpass-filtered speech, speech with a flattened pitch contour, and lowpass-filtered speech with a flattened pitch contour—and two additional low-level controls for the synthetic melody stimuli. The speech conditions were included to probe sensitivity to linguistic prosody for an unrelated study. The additional synthetic music control conditions were included to allow for a more rigorous interpretation of the intact > scrambled synthetic melodies effect had we observed such an effect. For completeness, on the OSF page, https://osf.io/68y7c/, we provide a data table that includes responses to these five conditions.)

For each participant, stimuli were randomly divided into six sets (corresponding to runs) with each set containing two stimuli from each condition. The order of the conditions for each run was selected from four predefined palindromic orders, which were constructed so that conditions targeting similar mental processes (e.g., orchestral music and single-instrument music) were separated by other conditions (e.g., speech or animal sounds). Each run contained three 10 s fixation periods: at the beginning, in the middle, and at the end. Otherwise, the stimuli were separated by 3 s fixation periods, for a total run duration of 456 s (7 min 36 s). All but two of the 18 participants completed all six runs (and thus got a total of 12 experimental events per condition); the remaining two completed four runs (and thus got 8 events per condition).

Because, as noted above, we have previously established that the language localizer is robust to presentation modality, we used the visual localizer to define the language regions. However, in SI-2 we show that the critical results are similar when auditory contrasts (*sentences > nonwords* in Experiment 1, or *Mandarin sentences* > *foreign* in Experiment 4) are instead used to define the language regions.

##### Experiment 2

Participants listened to well-formed melodies (adapted and expanded from Fedorenko et al. 2009) and melodies with a structural violation in a long-event-related design and judged the well-formedness of the melodies. As discussed in the Introduction, this type of manipulation is commonly used to probe sensitivity to music structure, including in studies examining language-music overlap (e.g., Patel et al. 1998; Koelsch et al. 2000, 2002; Maess et al. 2001; Tillmann et al. 2003; Fedorenko et al. 2009; Slevc et al. 2009; Kunert et al. 2015; Musso et al. 2015). The melodies were between 11 and 14 notes. The well-formed condition consisted of 90 melodies, which were tonal and ended in a tonic note with an authentic cadence in the implied harmony. All melodies were isochronous, consisting of quarter notes except for the final half note. The first five notes established a strong sense of key. Each melody was then altered to create a version with a “sour” note: the pitch of one note (from among the last four notes in a melody) was altered up or down by one or two semitones, so as to result in a non-diatonic note while keeping the melodic contour (the up-down pattern) the same. The structural position of the note that underwent this change varied among the tonic, the fifth, and the major third. The full set of 180 melodies was distributed across two lists following a Latin Square design. Any given participant heard stimuli from one list.

For each participant, stimuli were randomly divided into two sets (corresponding to runs) with each set containing 45 melodies (22 or 23 per condition). The order of the conditions, and the distribution of inter-trial fixation periods, was determined by the optseq2 algorithm (Dale et al. 1999). The order was selected from among four predefined orders, with no more than four trials of the same condition in a row. In each trial, participants were presented with a melody for three seconds followed by a question, presented visually on the screen, about the well-formedness of the melody (“Is the melody well-formed?”). To respond, participants had to press one of two buttons on a button box within two seconds. When participants answered, the question was replaced by a blank screen for the remainder of the two-second window; if no response was made within the two-second window, the experiment advanced to the next trial. Responses received within one second after the end of the previous trial were still recorded to account for the possible slow responses. The screen was blank during the presentation of the melodies. Each run contained 151 s of fixation interleaved among the trials, for a total run duration of 376 s (6 min 16 s). Fourteen of the 20 participants completed both runs, four participants completed one run, and the two remaining participants completed two runs but we only included their first run because, due to experimenter error, the second run came from a different experimental list and thus included some of the melodies from the first run in the other condition (the data pattern was qualitatively and quantitatively the same if both runs were included for these participants). Finally, due to a script error, participants only heard the first 12 notes of each melody during the three seconds of stimulus presentation. Therefore, we only analyzed the 80 of the 90 pairs (160 of the 180 total melodies) where the contrastive note appeared within the first 12 notes.

##### Experiment 4

Participants passively listened to single-instrument music, environmental sounds, sentences in their native language (Mandarin), and sentences in an unfamiliar foreign language (Russian) in a blocked design. All stimuli were 5-5.95s in length. The conditions were selected to probe responses to music, and to compare responses to music stimuli vs. other auditory stimuli. The critical music condition consisted of 60 stimuli selected from classical pieces by J.S. Bach played on cello, flute, or violin (n=15 each) and jazz music played on saxophone (n=15). The environmental sounds condition consisted of 60 stimuli selected from in-lab collections and included both pitched and unpitched stimuli. The foreign language condition consisted of 60 stimuli selected from Russian audiobooks (short stories by Paustovsky and “Fathers and Sons” by Turgenev). The foreign language condition was included because creating a ‘nonwords’ condition (the baseline condition we typically use for defining the language regions; Fedorenko et al. 2010) is challenging in Mandarin given that most words are monosyllabic, thus most syllables carry some meaning. As a result, sequences of syllables are more akin to lists of words. Therefore, we included the unfamiliar foreign language condition, which also works well as a baseline for language processing (Malik-Moraleda, Ayyash et al. 2022). The Mandarin sentence condition consisted of 120 lab-constructed sentences, each recorded by a male and a female native speaker. (The experiment also included five conditions that were not relevant to the current study and therefore not included in the analyses. These included three conditions probing responses to the participants’ second language (English) and two control conditions for Mandarin sentences. For completeness, on the OSF page, https://osf.io/68y7c/, we provide a data table that includes responses to these five conditions.)

Stimuli were grouped into blocks with each block consisting of three stimuli and lasting 18s (stimuli were padded with silence to make each trial exactly six seconds long). For each participant, blocks were divided into 10 sets (corresponding to runs), with each set containing two blocks from each condition. The order of the conditions for each run was selected from eight predefined palindromic orders. Each run contained three 14 s fixation periods: at the beginning, in the middle, and at the end, for a total run duration of 366 s (6 min 6 s). Five participants completed eight of the 10 runs (and thus got 16 blocks per condition; the remaining thirteen completed six runs (and thus got 12 blocks per condition). (We had created enough materials for 10 runs, but based on observing robust effects for several key contrasts in the first few participants who completed six to eight runs, we administered 6-8 runs to the remaining participants.)

Because we have previously found that an English localizer works well in native speakers of diverse languages, including Mandarin, as long as they are proficient in English (Malik-Moraleda, Ayyash et al. 2022), we used the same localizer in Experiment 4 as the one used in Experiments 1 and 2, for consistency. However, in SI-2 (**SI-Figure 2c, SI-Table 2c**) we show that the critical results are similar when the *Mandarin sentences > foreign* contrast is instead used to define the language regions.

#### Experiment 3 (behavioral)

*Language assessments.* Participants with aphasia were assessed for the integrity of lexical processing using word-to-picture matching tasks in both spoken and written modalities (ADA Spoken and Written Word-Picture Matching; Franklin et al. 1992). Productive vocabulary was assessed through picture naming. In the spoken modality, the Boston Naming Test was employed (Kaplan et al. 2001), and in writing, the PALPA Written Picture Naming subtest (Kay et al. 1992). Sentence processing was evaluated in both spoken and written modalities through comprehension (sentence-to-picture matching) of reversible sentences in active and passive voice. In a reversible sentence, the heads of both noun phrases are plausible agents, and therefore, word order, function words, and functional morphology are the only cues to who is doing what to whom. Participants also completed spoken and written grammaticality judgment tasks, where they made a yes/no decision as to the grammaticality of a word string. The task employed a subset of sentences from Linebarger et al. (1983).

All three participants exhibited severe language impairments that disrupted both comprehension and production (**Table 2**). For lexical-semantic tasks, all three participants displayed residual comprehension ability for high imageability/picturable vocabulary, although more difficulty was evident on the synonym matching test, which included abstract words. They were all severely anomic in speech and writing. Sentence production was severely impaired with output limited to single words, social speech (expressions, like “How are you?”), and other formulaic expressions (e.g., “and so forth”). Critically, all three performed at or close to chance level on spoken and written comprehension of reversible sentences and grammaticality judgments; each patient’s scores were lower than all of the healthy controls (**Table 2** and **Figure 2C**).

**Table 2.** Results of language assessments for participants with aphasia and healthy controls. For each test, we show number of correctly answered questions out of the total number of questions.

| Participant | SA | PR | PP | Controls |
| --- | --- | --- | --- | --- |
| **Lexical-semantic assessments** |  |  |  |  |
| ADA Spoken Word-Picture Matching<br>(chance = 16.5) | 60/66 | 61/66 | 64/66 | N/A |
| ADA Written Word-Picture Matching<br>(chance = 16.5) | 62/66 | 66/66 | 58/66 | N/A |
| ADA spoken synonym matching (chance = 80) | 123/160 | 121/160 | 135/160 | N/A |
| ADA written synonym matching (chance = 80) | 121/160 | 145/160 | 143/160 | N/A |
| Boston Naming Test<br>(NB: accepting both spoken and written responses) | 4/60 | 4/60 | 11/60 | N/A |
| PALPA 54 Written Picture Naming | 24/60 | 2/60 | 1/60 | N/A |
| **Syntactic assessments** |  |  |  |  |
| Comprehension of spoken reversible sentences (chance = 40) | 49/80 | 38/80 | 52/80 | Mean =<br>79.5/80<br>SD = 1.03<br>Min = 74/80<br>Max = 80/80<br>N=48 |
| Comprehension of written reversible sentences (chance = 40) | 42/80 | 49/80 | 51/80 | N/A |
| Spoken grammaticality judgments (chance = 24) | 33/48 | 34/48 | 35/48 | Mean =<br>45.5/48<br>SD = 2.52<br>Min = 36/48<br>Max = 48/48<br>N=49 |
| Written grammaticality judgments (chance = 24) | 29/48 | 24/48 | 29/48 | N/A |

##### Critical music task

Participants judged the well-formedness of the melodies from Experiment 2. Judgments were intended to reflect the detection of the key violation in the sour versions of the melodies. The full set of 180 melodies was distributed across two lists following a Latin Square design. All participants heard all 180 melodies. The control participants heard the melodies from one list, followed by the melodies from the other list, with the order of lists counter-balanced across participants. For the participants with aphasia, each list was further divided in half, and each participant was tested across four sessions, with 45 melodies per session, to minimize fatigue.

##### Montreal Battery for the Evaluation of Amusia

To obtain another measure of music competence/sensitivity to music structure, we administered the Montreal Battery for the Evaluation of Amusia (MBEA) (Peretz et al. 2003). The battery consists of six tasks that assess musical processing components described by Peretz and Coltheart (2003): three target melodic processing, two target rhythmic processing, and one assesses memory for melodies. Each task consists of 30 experimental trials (and uses the same set of 30 base melodies) and is preceded by practice examples. Some of the tasks additionally include a catch trial, as described below. For the purposes of the current investigation, the critical task is the “Scale” task. Participants are presented with pairs of melodies that they have to judge as identical or not. On half of the trials, one of the melodies is altered by modifying the pitch of one of the tones to be out of scale. Like our critical music task, this task aims to test participants’ ability to represent and use tonal structure in Western music, except that instead of making judgments on each individual melody, participants compare two melodies on each trial. This task thus serves as a conceptual replication (Schmidt 2009). One trial contains stimuli designed to be easy, intended as a catch trial to ensure that participants are paying attention. In this trial, the comparison melody has all its pitches set at random. This trial is excluded when computing the scores.

Control participants performed just the Scale task. Participants with aphasia performed all six tasks, distributed across three testing sessions to minimize fatigue.

#### fMRI data acquisition, preprocessing, and first-level modeling (for Experiments 1, 2, and 4)

##### Data acquisition

Whole-brain structural and functional data were collected on a whole-body 3 Tesla Siemens Trio scanner with a 32-channel head coil at the Athinoula A. Martinos Imaging Center at the McGovern Institute for Brain Research at MIT. T1-weighted structural images were collected in 176 axial slices with 1 mm isotropic voxels (repetition time (TR) = 2,530 ms; echo time (TE) = 3.48 ms). Functional, blood oxygenation level-dependent (BOLD) data were acquired using an EPI sequence with a 90° flip angle and using GRAPPA with an acceleration factor of 2; the following parameters were used: thirty-one 4.4 mm thick near-axial slices acquired in an interleaved order (with 10% distance factor), with an in-plane resolution of 2.1 mm × 2.1 mm, FoV in the phase encoding (A >> P) direction 200 mm and matrix size 96 × 96 voxels, TR = 2000 ms and TE = 30 ms. The first 10 s of each run were excluded to allow for steady state magnetization (see OSF https://osf.io/68y7c/ for the pdf of the scanning protocols). (Note that we opted to use a regular, continuous, scanning sequence in spite of investigating responses to auditory conditions. However, effects of scanner noise are unlikely to be detrimental given that all the stimuli are clearly perceptible, as also confirmed by examining responses in the auditory areas.)

##### Preprocessing

fMRI data were analyzed using SPM12 (release 7487), CONN EvLab module (release 19b), and other custom MATLAB scripts. Each participant’s functional and structural data were converted from DICOM to NIFTI format. All functional scans were coregistered and resampled using B-spline interpolation to the first scan of the first session (Friston et al. 1995). Potential outlier scans were identified from the resulting subject-motion estimates as well as from BOLD signal indicators using default thresholds in CONN preprocessing pipeline (5 standard deviations above the mean in global BOLD signal change, or framewise displacement values above 0.9 mm; Nieto-Castañón 2020). Functional and structural data were independently normalized into a common space (the Montreal Neurological Institute [MNI] template; IXI549Space) using SPM12 unified segmentation and normalization procedure (Ashburner and Friston 2005) with a reference functional image computed as the mean functional data after realignment across all timepoints omitting outlier scans. The output data were resampled to a common bounding box between MNI-space coordinates (-90, -126, -72) and (90, 90, 108), using 2mm isotropic voxels and 4th order spline interpolation for the functional data, and 1mm isotropic voxels and trilinear interpolation for the structural data. Last, the functional data were smoothed spatially using spatial convolution with a 4 mm FWHM Gaussian kernel.

##### First-level modeling

For both the language localizer task and the critical experiments, effects were estimated using a General Linear Model (GLM) in which each experimental condition was modeled with a boxcar function convolved with the canonical hemodynamic response function (HRF) (fixation was modeled implicitly, such that all timepoints that did not correspond to one of the conditions were assumed to correspond to a fixation period). Temporal autocorrelations in the BOLD signal timeseries were accounted for by a combination of high-pass filtering with a 128 seconds cutoff, and whitening using an AR(0.2) model (first-order autoregressive model linearized around the coefficient a=0.2) to approximate the observed covariance of the functional data in the context of Restricted Maximum Likelihood estimation (ReML). In addition to experimental condition effects, the GLM design included first-order temporal derivatives for each condition (included to model variability in the HRF delays), as well as nuisance regressors to control for the effect of slow linear drifts, subject-motion parameters, and potential outlier scans on the BOLD signal.

#### Definition of the language functional regions of interest (fROIs) (for Experiments 1, 2, and 4)

For each critical experiment, we defined a set of language functional regions of interest (fROIs) using group-constrained, subject-specific localization (Fedorenko et al. 2010). In particular, each individual map for the *sentences > nonwords* contrast from the language localizer was intersected with a set of five binary masks. These masks (**Figure 3**; available at OSF: https://osf.io/68y7c/) were derived from a probabilistic activation overlap map for the same contrast in a large independent set of participants (n=220) using watershed parcellation, as described in Fedorenko et al. (2010) for a smaller set of participants. These masks covered the fronto-temporal language network in the left hemisphere. Within each mask, a participant-specific language fROI was defined as the top 10% of voxels with the highest *t*-values for the localizer contrast.

**Figure 3.**
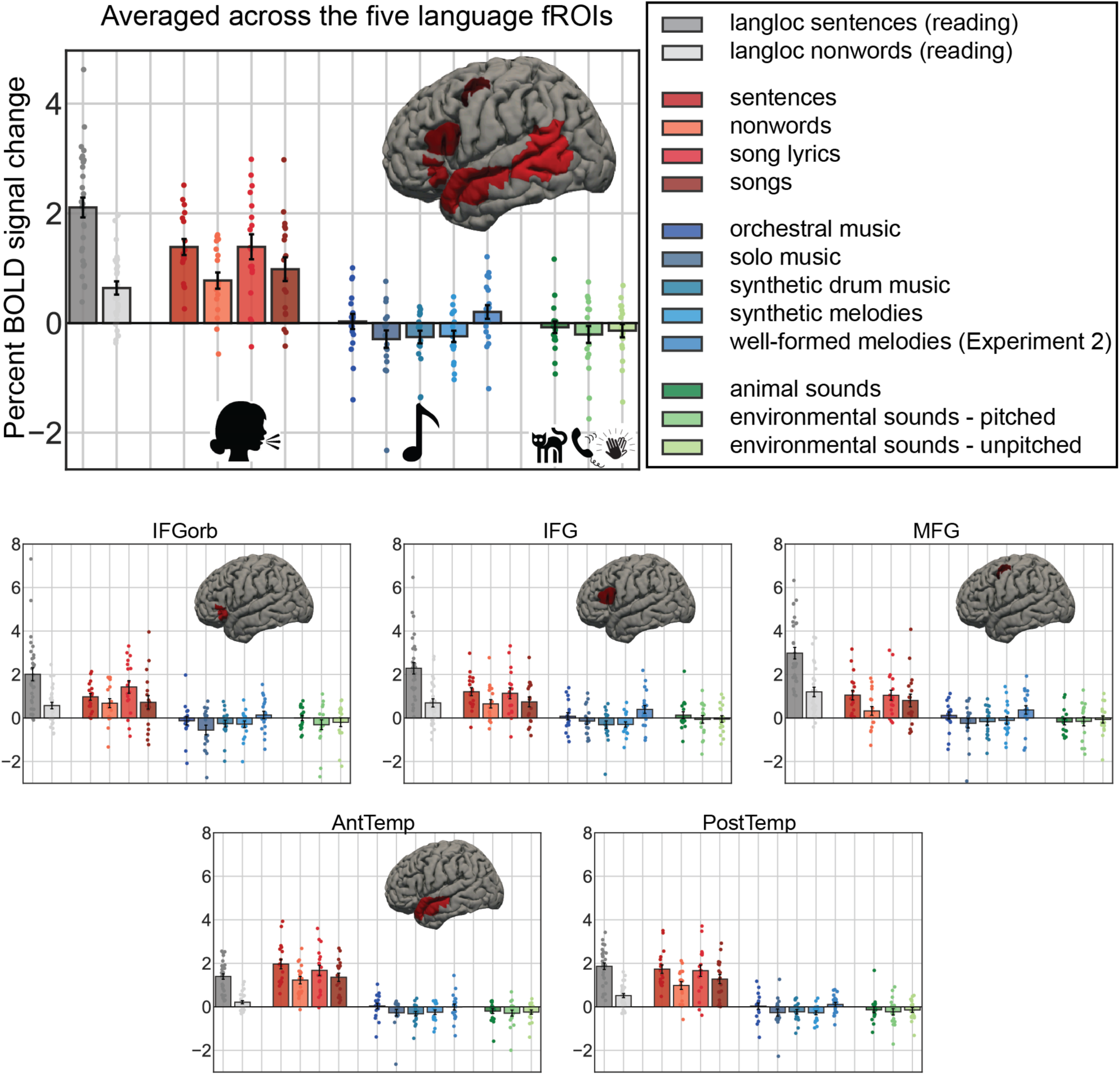
Responses of the language fROIs (pooling across the network – top, and for each fROI individually – bottom) to the language localizer conditions (in grey), to the four auditory conditions containing speech in Experiment 1 (red shades), to the five music conditions in Experiments 1 and 2 (blue shades), and to the three non-linguistic/non-music auditory conditions (green shades) in Experiment 1. Here and elsewhere, the error bars represent standard errors of the mean by participants. For the language localizer results, we include here all participants in Experiments 1 and 2. The responses to the music conditions cluster around the fixation baseline, are much lower than the responses to sentences, and are not higher than the responses to non-music sounds.

#### Validation of the language fROIs

To ensure that the language fROIs behave as expected (i.e., show a reliably greater response to the sentences condition compared to the nonwords condition), we used an across-runs cross-validation procedure (e.g., Nieto-Castañón and Fedorenko 2012). In this analysis, the first run of the localizer was used to define the fROIs, and the second run to estimate the responses (in percent BOLD signal change, PSC) to the localizer conditions, ensuring independence (e.g., Kriegeskorte et al. 2009); then the second run was used to define the fROIs, and the first run to estimate the responses; finally, the extracted magnitudes were averaged across the two runs to derive a single response magnitude for each of the localizer conditions. Statistical analyses were performed on these extracted PSC values. Consistent with much previous work (e.g., Fedorenko et al. 2010; Mahowald and Fedorenko 2016; Diachek, Blank, Siegelman et al. 2020), each of the language fROIs showed a robust *sentences > nonwords* effect (all *p*s < 0.001).

### Statistical analyses for the fMRI experiments

All analyses were performed with linear mixed-effects models using the “lme4” package in R with *p*-value approximation performed by the “lmerTest” package (Bates et al. 2015; Kuznetsova et al. 2017). Effect size (Cohen’s d) was calculated using the method from Westfall et al. (2014) and Brysbaert and Stevens (2018).

#### Sanity check analyses and results

To estimate the responses in the language fROIs to the conditions of the critical experiments here and in the critical analyses, the data from all the runs of the language localizer were used to define the fROIs, and the responses to each condition were then estimated in these regions. Statistical analyses were then performed on these extracted PSC values. (For Experiments 1 and 4, we repeated the analyses using alternative language localizer contrasts to define the language fROIs (auditory *sentences > nonwords* in Experiment 1, and *Mandarin sentences > foreign* in Experiment 4), which yielded quantitatively and qualitatively similar responses (see **SI-2**).)

We conducted two sets of sanity check analyses. First, to ensure that auditory conditions that contain meaningful linguistic content elicit strong responses in the language regions relative to perceptually similar conditions with no discernible linguistic content, we compared the auditory sentences condition with the auditory nonwords condition (Experiment 1) or with the foreign language condition (Experiment 4). Indeed, as expected, the auditory sentence condition elicited a stronger response than the auditory nonwords condition (Experiment 1) or the foreign language condition (Experiment 4). These effects were robust at the network level (*p*s < 0.001; **SI-Table 1a**). Further, the *sentences* > *nonwords* effect was significant in all but one language fROI in Experiment 1, and the *sentences* > *foreign* effect was significant in all language fROIs in Experiment 4 (*p*s < 0.05; **SI-Table 1a**).

And second, to ensure that the music conditions elicit strong responses in auditory cortex, we extracted the responses from a bilateral anatomically defined auditory cortical region (area Te1.2 from the Morosan et al. 2001 cytoarchitectonic probabilistic atlas) to the six critical music conditions: orchestral music, single instrument music, synthetic drum music, and synthetic melodies in Experiment 1; well-formed melodies in Experiment 2; and the music condition in Experiment 4. Statistical analyses, comparing each condition to the fixation baseline, were performed on these extracted PSC values. As expected, all music conditions elicited strong responses in a primary auditory area bilaterally (all *p*s ≅ 0.001; **SI-Table 1b; SI-Figure 1**).

#### Critical analyses

To characterize the responses in the language network to music perception, we asked three questions. First, we asked whether music conditions elicit strong responses in the language regions. Second, we investigated whether the language network is sensitive to structure in music, as would be evidenced by stronger responses to intact than scrambled music, and stronger responses to melodies with structural violations compared to the no-violation control condition. And third, we asked whether music conditions elicit strong responses in the language regions of individuals with high sensitivity to linguistic pitch—native speakers of a tonal language (Mandarin).

For each contrast (the contrasts relevant to the three research questions are detailed below), we used two types of linear mixed-effect regression models:

i. the language network model, which examined the language network as a whole; and
ii. the individual language fROI models, which examined each language fROI separately.

As alluded to in the Introduction, treating the language network as an integrated system is reasonable given that the regions of this network a) show similar functional profiles, both with respect to selectivity for language over non-linguistic processes (e.g., Fedorenko et al. 2011; Pritchett et al. 2018; Jouravlev et al. 2019; Ivanova et al. 2020, 2021) and with respect to their role in lexico-semantic and syntactic processing (e.g., Fedorenko et al. 2012b; Blank et al. 2016; Fedorenko et al. 2020); and b) exhibit strong inter-region correlations in both their activity during naturalistic cognition paradigms (e.g., Blank et al. 2014; Braga et al. 2020; Paunov et al. 2019; Malik-Moraleda, Ayyash et al. 2022) and key functional markers, like the strength or extent of activation in response to language stimuli (e.g., Mahowald and Fedorenko 2016; Mineroff, Blank et al. 2018). However, to allow for the possibility that language regions differ in their response to music and to examine the region on which most claims about language-music overlap have focused (the region that falls within ‘Broca’s area’), we supplement the network-wise analyses with the analyses of the five language fROIs separately.

For each network-wise analysis, we fit a linear mixed-effect regression model predicting the level of BOLD response in the language fROIs in the contrasted conditions. The model included a fixed effect for condition (the relevant contrasts are detailed below for each analysis) and random intercepts for fROIs and participants. Here and elsewhere, the *p*-value was estimated by applying the Satterthwaite’s method-of-moment approximation to obtain the degrees of freedom (Giesbrecht and Burns 1985; Fai and Cornelius 1996; as described in Kuznetsova et al. 2017). For the comparison against the fixation baseline, the random intercept for participants was removed because it is no longer applicable.

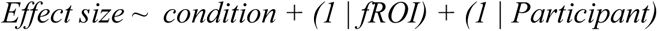

For each fROI-wise analysis, we fit a linear mixed-effect regression model predicting the level of BOLD response in each of the five language fROIs in the contrasted conditions. The model included a fixed effect for condition and random intercepts for participants. For each analysis, the results were FDR-corrected for the five fROIs. For the comparison against the fixation baseline, the random intercept for participants was removed because it is no longer applicable.

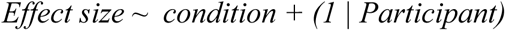

## Results

### Does music elicit a response in the language network?

As discussed in the Introduction, a brain region that supports (some aspect of) music processing, including structure processing, should show a strong response to music stimuli. To test whether language regions respond to music, we used four contrasts using data from Experiments 1 and 2. First, we compared the responses to each of the music conditions (orchestral music, single instrument music, synthetic drum music, and synthetic melodies in Experiment 1; well-formed melodies in Experiment 2) against the fixation baseline—the most liberal baseline. Second, we compared the responses to the music conditions against the response to the nonword strings condition—an unstructured and meaningless linguistic stimulus (in Experiment 1, we used the auditory nonwords condition, and in Experiment 2, we used the visual nonwords condition from the language localizer). Third, in Experiment 1, we additionally compared the responses to the music conditions against the response to non-linguistic, non-music stimuli (animal and environmental sounds). A brain region that supports music processing should elicit a strong positive response relative to the fixation baseline and the nonwords condition (our baseline for the language regions); further, if the response is selective, it should be stronger than the response elicited by non-music auditory stimuli. Finally, in Experiment 1, we also directly compared the responses to songs vs. lyrics. A brain region that responds to music should respond more strongly to songs given that they contain a melodic contour in addition to the linguistic content.

None of the music conditions elicited a strong response in the language network (**Figure 3**; **Table 3**). The responses to music (i) fell at or below the fixation baseline (except for the well-formed melodies condition in Experiment 2, which elicited a small positive response in some regions), (ii) were lower than the response elicited by auditory nonwords (except for the LMFG language fROI, where the responses to music and nonwords were similarly low), and (iii) did not significantly differ from the responses elicited by non-linguistic, non-music conditions. Finally, the response to songs, which contain both linguistic content and a melodic contour, was not significantly higher than the response elicited by the linguistic content alone (lyrics); in fact, at the network level, the response to songs was reliably lower than to lyrics.

**Table 3.** Statistical results (two-sided) for the contrasts between music conditions and three kinds of baselines (fixation, nonwords, and non-linguistic non-music auditory conditions—animal sounds and environmental sounds) in Experiments 1 and 2, and for the contrast between songs and lyrics in Experiment 1. Abbreviations: b=the beta estimate for the effect; se=standard error of the mean by participants; df=degrees of freedom; d=Cohen’s d (Westfall et al. 2014; Brysbaert and Stevens 2018); t=the t statistic; p=the significance value (for the individual fROIs, these values have been FDR-corrected for the number of fROIs (n=5)). In light grey, we highlight the results that are ***not consistent*** with the role of the language regions in music perception: of the 84 tests performed, 1 showed an effect predicted by language-music overlap accounts: a small and statistically weak (only emerging at the network level but not in any individual fROI) positive response, relative to the weakest baseline (fixation), to one of the five music conditions examined; and this response was still ∼4 times lower than the response to an unstructured linguistic condition (nonwords).

| Contrast | Language<br>network | LIFGorb | LIFG | LMFG | LAnt<br>Temp | LPost<br>Temp |
| --- | --- | --- | --- | --- | --- | --- |
| *music > fixation* |  |  |  |  |  |  |
| orchestral music<br>>fixation | $\beta=0.028$<br>se=0.135<br>df=17.383<br>d=0.042<br>t=0.209<br>p=0.837 | $\beta=-0.129$<br>se=0.193<br>df=nan<br>d=nan<br>t=-0.667<br>p=1.000 | $\beta=0.082$<br>se=0.162<br>df=nan<br>d=nan<br>t=0.506<br>p=1.000 | $\beta=0.117$<br>se=0.165<br>df=nan<br>d=nan<br>t=0.711<br>p=1.000 | $\beta=0.040$<br>se=0.130<br>df=nan<br>d=nan<br>t=0.310<br>p=1.000 | $\beta=0.030$<br>se=0.143<br>df=nan<br>d=nan<br>t=0.210<br>p=1.000 |
| single-instrument<br>music<br>>fixation | $\beta=-0.294$<br>se=0.163<br>df=18.616<br>d=-0.378<br>t=-1.809<br>p=0.087 | $\beta=-0.552$<br>se=0.223<br>df=nan<br>d=nan<br>t=-2.471<br>p=0.120 | $\beta=-0.141$<br>se=0.155<br>df=nan<br>d=nan<br>t=-0.906<br>p=1.000 | $\beta=-0.243$<br>se=0.217<br>df=nan<br>d=nan<br>t=-1.122<br>p=1.000 | $\beta=-0.272$<br>se=0.152<br>df=nan<br>d=nan<br>t=-1.794<br>p=0.455 | $\beta=-0.264$<br>se=0.164<br>df=nan<br>d=nan<br>t=-1.611<br>p=0.630 |
| synthetic drum<br>music<br>>fixation | $\beta=-0.255$<br>se=0.112<br>df=18.000<br>d=-0.432<br>t=-2.281<br>p=0.035* | $\beta=-0.258$<br>se=0.155<br>df=nan<br>d=nan<br>t=-1.667<br>p=0.570 | $\beta=-0.306$<br>se=0.172<br>df=nan<br>d=nan<br>t=-1.780<br>p=0.465 | $\beta=-0.168$<br>se=0.162<br>df=nan<br>d=nan<br>t=-1.040<br>p=1.000 | $\beta=-0.319$<br>se=0.106<br>df=nan<br>d=nan<br>t=-3.020<br>p=0.040* | $\beta=-0.226$<br>se=0.103<br>df=nan<br>d=nan<br>t=-2.189<br>p=0.215 |
| synthetic<br>melodies | $\beta=-0.243$<br>se=0.100 | $\beta=-0.286$<br>se=0.154 | $\beta=-0.299$<br>se=0.120 | $\beta=-0.108$<br>se=0.177 | $\beta=-0.247$<br>se=0.103<br>df=nan | $\beta=-0.276$<br>se=0.089<br>df=nan |
| >fixation | df=18.000<br>d=-0.441<br>t=-2.423<br>p=0.026* | df=nan<br>d=nan<br>t=-1.856<br>p=0.405 | df=nan<br>d=nan<br>t=-2.485<br>p=0.120 | df=nan<br>d=nan<br>t=-0.611<br>p=1.000 | d=nan<br>t=-2.395<br>p=0.140 | d=nan<br>t=-3.093<br>p=0.035* |
| well-formed<br>melodies (Expt<br>2)<br>>fixation | $\beta$=0.201<br>se=0.135<br>df=17.483<br>d=0.281<br>t=1.488<br>p=0.155 | $\beta$=0.139<br>se=0.166<br>df=nan<br>d=nan<br>t=0.836<br>p=1.000 | $\beta$=0.396<br>se=0.181<br>df=nan<br>d=nan<br>t=2.182<br>p=0.210 | $\beta$=0.371<br>se=0.197<br>df=nan<br>d=nan<br>t=1.885<br>p=0.375 | $\beta$=-0.008<br>se=0.136<br>df=nan<br>d=nan<br>t=-0.056<br>p=1.000 | $\beta$=0.107<br>se=0.096<br>df=nan<br>d=nan<br>t=1.109<br>p=1.000 |
| ***music > nonwords*** |  |  |  |  |  |  |
| orchestral music<br>>nonwords | $\beta$=-0.746<br>se=0.092<br>df=157.707<br>d=-0.978<br>t=-8.097<br>p<0.001*** | $\beta$=-0.811<br>se=0.276<br>df=36.000<br>d=-0.981<br>t=-2.945<br>p=0.030* | $\beta$=-0.569<br>se=0.142<br>df=18.000<br>d=-0.779<br>t=-4.015<br>p=0.005* | $\beta$=-0.210<br>se=0.221<br>df=18.000<br>d=-0.276<br>t=-0.954<br>p=1.000 | $\beta$=-1.187<br>se=0.147<br>df=18.000<br>d=-1.884<br>t=-8.101<br>p<0.001*** | $\beta$=-0.950<br>se=0.205<br>df=18.000<br>d=-1.427<br>t=-4.646<br>p<0.001*** |
| single-instrument<br>music<br>>nonwords | $\beta$=-1.068<br>se=0.100<br>df=157.689<br>d=-1.314<br>t=-10.714<br>p<0.001*** | $\beta$=-1.234<br>se=0.296<br>df=36.000<br>d=-1.388<br>t=-4.167<br>p<0.001*** | $\beta$=-0.791<br>se=0.222<br>df=18.000<br>d=-1.101<br>t=-3.567<br>p=0.010* | $\beta$=-0.571<br>se=0.235<br>df=18.000<br>d=-0.661<br>t=-2.431<br>p=0.130 | $\beta$=-1.500<br>se=0.196<br>df=18.000<br>d=-2.236<br>t=-7.648<br>p<0.001*** | $\beta$=-1.244<br>se=0.234<br>df=17.998<br>d=-1.765<br>t=-5.315<br>p<0.001*** |
| synthetic drum<br>music<br>>nonwords | $\beta$=-1.029<br>se=0.087<br>df=157.720<br>d=-1.408 | $\beta$=-0.940<br>se=0.212<br>df=18.000<br>d=-1.246 | $\beta$=-0.956<br>se=0.182<br>df=18.000<br>d=-1.275 | $\beta$=-0.496<br>se=0.245<br>df=18.000<br>d=-0.658 | $\beta$=-1.546<br>se=0.187<br>df=18.000<br>d=-2.621 | $\beta$=-1.207<br>se=0.177<br>df=18.000<br>d=-2.012 |
|  | t=-11.839<br>p<0.001*** | t=-4.430<br>p<0.001*** | t=-5.252<br>p<0.001*** | t=-2.026<br>p=0.290 | t=-8.262<br>p<0.001*** | t=-6.817<br>p<0.001*** |
| synthetic<br>melodies<br>-nonwords | $\beta$=-1.017<br>se=0.087<br>df=157.683<br>d=-1.421<br>t=-11.623<br>p<0.001*** | $\beta$=-0.969<br>se=0.209<br>df=18.000<br>d=-1.286<br>t=-4.642<br>p<0.001*** | $\beta$=-0.949<br>se=0.153<br>df=18.000<br>d=-1.441<br>t=-6.223<br>p<0.001*** | $\beta$=-0.435<br>se=0.252<br>df=18.000<br>d=-0.556<br>t=-1.727<br>p=0.505 | $\beta$=-1.474<br>se=0.195<br>df=36.000<br>d=-2.513<br>t=-7.541<br>p<0.001*** | $\beta$=-1.256<br>se=0.176<br>df=18.000<br>d=-2.164<br>t=-7.136<br>p<0.001*** |
| well-formed<br>melodies (Expt<br>2)<br>>nonwords<br>(visual) | $\beta$=-0.449<br>se=0.090<br>df=179.063<br>d=-0.562<br>t=-4.998<br>p<0.001*** | $\beta$=-0.490<br>se=0.226<br>df=20.989<br>d=-0.611<br>t=-2.164<br>p=0.210 | $\beta$=-0.403<br>se=0.208<br>df=20.056<br>d=-0.444<br>t=-1.938<br>p=0.335 | $\beta$=-0.686<br>se=0.250<br>df=20.173<br>d=-0.705<br>t=-2.748<br>p=0.060 | $\beta$=-0.242<br>se=0.134<br>df=20.737<br>d=-0.470<br>t=-1.812<br>p=0.420 | $\beta$=-0.375<br>se=0.123<br>df=20.455<br>d=-0.792<br>t=-3.056<br>p=0.030* |
| ***music > non-linguistic, non-music auditory conditions*** |  |  |  |  |  |  |
| music<br>(combined)<br>>animal sounds | $\beta$=-0.114<br>se=0.060<br>df=427.876<br>d=-0.177<br>t=-1.915<br>p=0.056 | $\beta$=-0.306<br>se=0.148<br>df=72.000<br>d=-0.422<br>t=-2.069<br>p=0.210 | $\beta$=-0.295<br>se=0.146<br>df=72.000<br>d=-0.451<br>t=-2.021<br>p=0.235 | $\beta$=0.080<br>se=0.151<br>df=72.000<br>d=0.111<br>t=0.528<br>p=1.000 | $\beta$=-0.002<br>se=0.090<br>df=72.000<br>d=-0.004<br>t=-0.023<br>p=1.000 | $\beta$=-0.048<br>se=0.094<br>df=72.000<br>d=-0.088<br>t=-0.513<br>p=1.000 |
| music<br>(combined)<br>>environmental<br>(pitched) | $\beta$=0.019<br>se=0.060<br>df=427.902<br>d=0.028<br>t=0.307<br>p=0.759 | $\beta$=0.005<br>se=0.144<br>df=72.000<br>d=0.006<br>t=0.033<br>p=1.000 | $\beta$=-0.104<br>se=0.133<br>df=72.000<br>d=-0.156<br>t=-0.781<br>p=1.000 | $\beta$=0.055<br>se=0.159<br>df=72.000<br>d=0.071<br>t=0.347<br>p=1.000 | $\beta$=0.092<br>se=0.094<br>df=72.000<br>d=0.171<br>t=0.975<br>p=1.000 | $\beta$=0.045<br>se=0.094<br>df=72.000<br>d=0.081<br>t=0.475<br>p=1.000 |
| music<br>(combined)<br>>environmental<br>(unpitched) | $\beta=-0.052$<br>se=0.063<br>df=427.856<br>d=-0.079<br>t=-0.823<br>p=0.411 | $\beta=-0.109$<br>se=0.163<br>df=72.000<br>d=-0.140<br>t=-0.666<br>p=1.000 | $\beta=-0.118$<br>se=0.152<br>df=72.000<br>d=-0.182<br>t=-0.778<br>p=1.000 | $\beta=-0.030$<br>se=0.151<br>df=72.000<br>d=-0.040<br>t=-0.198<br>p=1.000 | $\beta=0.042$<br>se=0.097<br>df=72.000<br>d=0.083<br>t=0.429<br>p=1.000 | $\beta=-0.043$<br>se=0.100<br>df=72.000<br>d=-0.082<br>t=-0.426<br>p=1.000 |
| ***(melodic contour + linguistic content) > linguistic content*** |  |  |  |  |  |  |
| songs<br>>lyrics | $\beta=-0.408$<br>se=0.102<br>df=157.896<br>d=-0.377<br>t=-4.014<br>p<0.001*** | $\beta=-0.705$<br>se=0.287<br>df=18.000<br>d=-0.569<br>t=-2.454<br>p=0.125 | $\beta=-0.394$<br>se=0.195<br>df=18.000<br>d=-0.400<br>t=-2.025<br>p=0.290 | $\beta=-0.243$<br>se=0.219<br>df=18.000<br>d=-0.226<br>t=-1.107<br>p=1.000 | $\beta=-0.313$<br>se=0.163<br>df=18.000<br>d=-0.356<br>t=-1.925<br>p=0.350 | $\beta=-0.384$<br>se=0.171<br>df=18.000<br>d=-0.392<br>t=-2.246<br>p=0.185 |

#### Is the language network sensitive to structure in music?

##### Experiments 1 and 2 (fMRI)

Because most prior claims about the overlap between language and music concern the processing of *structure*—given the parallels that can be drawn between the syntactic structure of language and the tonal and rhythmic structure in music (e.g., Lerdahl and Jackendoff 1977, 1983; cf. Jackendoff 2009)—we used three contrasts to test whether language regions are sensitive to music structure. First and second, in Experiment 1, we compared the responses to synthetic melodies vs. their scrambled counterparts, and to synthetic drum music vs. the scrambled drum music condition. The former targets both tonal and rhythmic structure, and the latter selectively targets rhythmic structure. The reason to examine rhythmic structure is that some patient studies have argued that pitch contour processing relies on the right hemisphere, and rhythm processing draws on the left hemisphere (e.g., Zatorre 1984; Peretz 1990; Alcock et al. 2000; cf. Boebinger 2021 for fMRI evidence of bilateral responses in high-level auditory areas to both tonal and rhythmic structure processing and for lack of spatial segregation between the two), so although most prior work examining the language-music relationship has focused on tonal structure, rhythmic structure may *a priori* be more likely to overlap with linguistic syntactic structure given their alleged co-lateralization based on the patient literature. And third, in Experiment 2, we compared the responses to well-formed melodies vs. melodies with a sour note. A brain region that responds to structure in music should respond more strongly to intact than scrambled music (similar to how language regions respond more strongly to sentences than lists of words; e.g., Fedorenko et al. 2010; Diachek, Blank, Siegelman et al. 2020), and also exhibit sensitivity to structure violations (similar to how language regions respond more strongly to sentences that contain grammatical errors: e.g., Embick et al. 2000; Newman et al. 2001; Kuperberg et al. 2003; Cooke et al. 2006; Friederici et al. 2010; Herrmann et al. 2012; Fedorenko et al. 2020). Note that given the lack of a strong and consistent response to music in the language regions (**Figure 3** and **Table 3**), the answer to this narrower question is somewhat of a foregone conclusion: even if one or more of the language regions showed a reliable effect in these music-structure-probing contrasts, such effects would be difficult to interpret as reflecting music structure processing given that structured music stimuli elicit a response approximately at the level of the fixation baseline in the language areas. Nevertheless, we report the results for these three contrasts for completeness, and because most prior studies have focused on such contrasts.

The language regions did not show consistent sensitivity to structural manipulations in music (**Figure 4**; **Table 4**). In Experiment 1, the responses to synthetic melodies did not significantly differ from (or were weaker than) the responses to the scrambled counterparts, and the responses to synthetic drum music did not significantly differ from the responses to scrambled drum music. In Experiment 2, at the network level, we observed a small and weakly significant (p<0.05) effect of *sour-note* > *well-formed melodies*. This effect was not significant in any of the five individual fROIs (even prior to the FDR correction).

**Figure 4.**
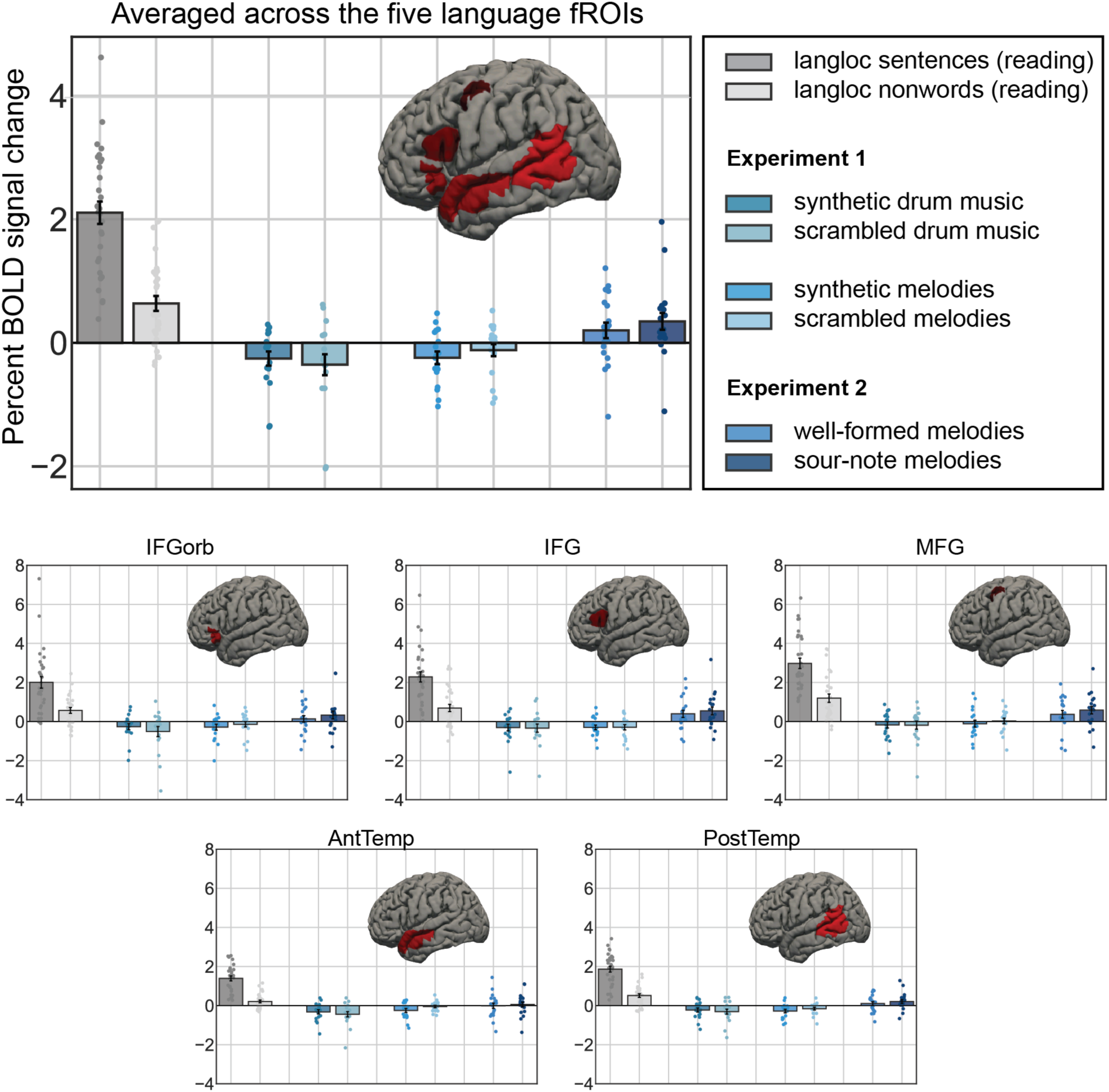
Responses of the language fROIs (pooling across the network – top, and for each fROI individually – bottom) to the language localizer conditions (in grey), and to the three sets of conditions that target structure in music (in blue). The error bars represent standard error of the mean by participants. For the language localizer results, we include here participants in Experiments 1 and 2. The responses to the music conditions cluster around the fixation baseline, and are much lower than the response to sentences. One of the three critical contrasts (*sour-note > well-formed* melodies) elicits a small and weakly reliable effect at the network level, but it is not individually significant in any of the five fROIs.

**Table 4.** Statistical results (two-sided) for the contrasts between the synthetic drum music and scrambled drum music, synthetic melodies and scrambled synthetic melodies, and sour-note and well-formed melodies contrasts in Experiments 1 and 2. Abbreviations: b=the beta estimate for the effect; se=standard error of the mean by participants; df=degrees of freedom; d=Cohen’s d (Westfall et al. 2014; Brysbaert and Stevens 2018); t=the t statistic; p=the significance value (for the individual fROIs, these values have been FDR-corrected for the number of fROIs (n=5)). In light grey, we highlight the results that are **not consistent** with the role of the language regions in the processing of music structure: of the 18 tests performed, 1 showed an effect predicted by language-music overlap accounts: a small and statistically weak response to one of the three structure-targeting contrasts (in the presence of an overall very weak response to music relative to fixation; see Figure 3 and Table 3).

| Contrast | Language<br>network | LIFGorb | LIFG | LMFG | LAnt<br>Temp | LPost<br>Temp |
| --- | --- | --- | --- | --- | --- | --- |
| synthetic drum<br>music<br>>scrambled<br>drum music | $\beta=0.099$<br>se=0.073<br>df=157.823<br>d=0.140<br>t=1.358<br>p=0.176 | $\beta=0.252$<br>se=0.191<br>df=18.000<br>d=0.288<br>t=1.322<br>p=1.000 | $\beta=0.027$<br>se=0.176<br>df=18.000<br>d=0.034<br>t=0.156<br>p=1.000 | $\beta=0.014$<br>se=0.186<br>df=18.000<br>d=0.018<br>t=0.073<br>p=1.000 | $\beta=0.124$<br>se=0.103<br>df=18.000<br>d=0.247<br>t=1.210<br>p=1.000 | $\beta=0.079$<br>se=0.110<br>df=18.000<br>d=0.165<br>t=0.718<br>p=1.000 |
| synthetic<br>melodies<br>>scrambled<br>synthetic<br>melodies | $\beta=-0.124$<br>se=0.061<br>df=157.720<br>d=-0.238<br>t=-2.015<br>p=0.046* | $\beta=-0.147$<br>se=0.130<br>df=18.000<br>d=-0.245<br>t=-1.133<br>p=1.000 | $\beta=-0.009$<br>se=0.153<br>df=18.000<br>d=-0.017<br>t=-0.057<br>p=1.000 | $\beta=-0.143$<br>se=0.202<br>df=18.000<br>d=-0.216<br>t=-0.708<br>p=1.000 | $\beta=-0.199$<br>se=0.101<br>df=18.000<br>d=-0.572<br>t=-1.971<br>p=0.320 | $\beta=-0.121$<br>se=0.106<br>df=18.000<br>d=-0.365<br>t=-1.142<br>p=1.000 |
| sour-note<br>melodies<br>>well-formed<br>melodies | $\beta=0.145$<br>se=0.069<br>df=175.884<br>d=0.196<br>t=2.102<br>p=0.037* | $\beta=0.195$<br>se=0.098<br>df=20.000<br>d=0.245<br>t=1.985<br>p=0.305 | $\beta=0.150$<br>se=0.105<br>df=20.000<br>d=0.180<br>t=1.431<br>p=0.840 | $\beta=0.212$<br>se=0.090<br>df=20.000<br>d=0.252<br>t=2.363<br>p=0.140 | $\beta=0.065$<br>se=0.051<br>df=20.000<br>d=0.114<br>t=1.280<br>p=1.000 | $\beta=0.104$<br>se=0.056<br>df=20.000<br>d=0.248<br>t=1.856<br>p=0.390 |

##### Experiment 3 (behavioral)

In Experiment 3, we further asked whether individuals with severe deficits in processing linguistic syntax also exhibit difficulties in processing music structure. To do so, we assessed participants’ ability to discriminate well-formed (“good”) melodies from melodies with a sour note (“bad”), while controlling for their response bias (how likely they are overall to say that something is well-formed) by computing *d’* for each participant (Green and Swets 1966), in addition to proportion correct. We then compared the *d’* values of each individual with aphasia to the distribution of *d’* values of healthy control participants using a Bayesian test for single case assessment (Crawford and Garthwaite 2007) as implemented in the *psycho* package in R (Makowski 2018). (Note that for the linguistic syntax tasks, it was not necessary to conduct statistical tests comparing the performance of each individual with aphasia to the control distribution because the performance of each individual with aphasia was lower than 100% of the control participants’ performances.) We similarly compared the proportion correct on the MBEA scale task of each individual with aphasia to the distribution of accuracies of healthy controls. If linguistic and music syntax draw on the same resources, then individuals with linguistic syntactic impairments should also exhibit deficits on tasks requiring the processing of music syntax.

In the critical music task, where participants were asked to judge the well-formedness of musical structure, neurotypical control participants responded correctly, on average, on 87.1% of trials, suggesting that the task was sufficiently difficult to preclude ceiling effects. Patients with severe aphasia showed intact sensitivity to music structure. The three patients had accuracies of 89.4% (PR), 94.4% (SA), and 97.8% (PP), falling on the higher end of the controls’ performance range (**Figure 5**; **Table 5**). Crucially, none of the three aphasic participants’ *d’* scores were lower than the average control participants’ *d’* scores (M = 2.75, SD = 0.75). In fact, the patients’ *d’* scores were high: SA’s *d’* was 3.51, higher than 83.91% (95% Credible Interval (CI) [75.20, 92.03]) of the control population, PR’s *d’* was 3.09, higher than 67.26% (95% CI [56.60, 78.03]) of the control population, and PP’s *d’* was 3.99, higher than 94.55% (95% CI [89.40, 98.57]) of the control population. None of the three aphasic participants’ bias/criterion c scores (Green and Swets 1966) differed reliably from the control participants’ c scores (M = -0.40, SD = 0.40). SA’s c was -0.53, lower than 62.34% (95% CI [50.40, 71.67]) of the control population, PR’s c was -0.74, lower than 79.48% (95% CI [69.58, 88.44]) of the control population, and PP’s c was-0.29, higher than 60.88% (95% CI [50.08, 70.04]) of the control population. In the Scale task from the Montreal Battery for the Evaluation of Aphasia, the control participants’ performance showed a similar distribution to that reported in Peretz et al. (2003). All participants with aphasia performed within the normal range, with two participants making no errors. PR and PP’s score was higher than 85.24% (95% CI [76.94, 93.06]) of the control population, providing a conceptual replication of the results from the well-formed/sour-note melody discrimination task. SA’s score was higher than 30.57% (95% CI [20.00, 41.50]) of the control population.

**Figure 5.**
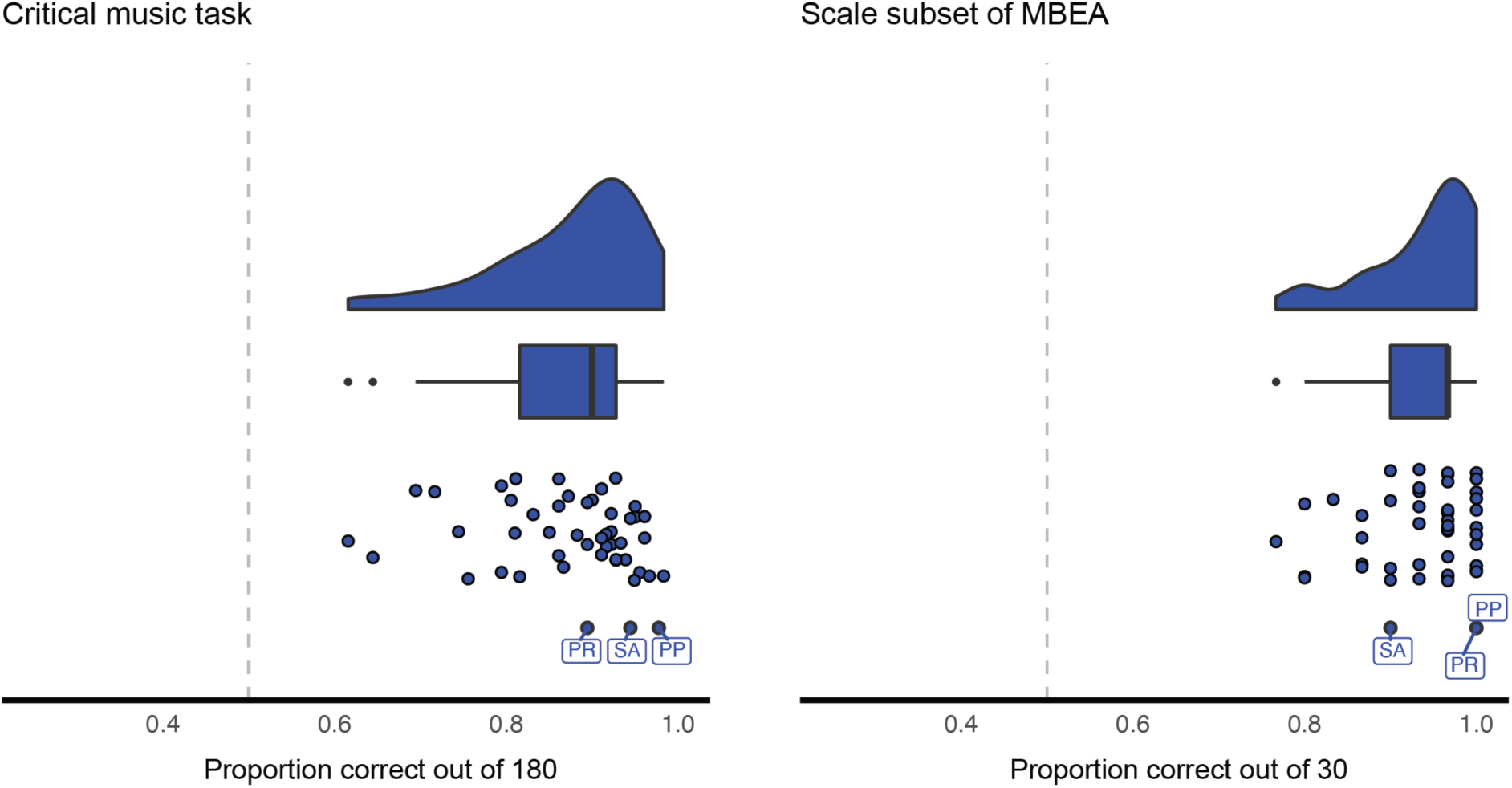
Performance of the control and aphasic participants on two measures of music syntax processing: the critical music task (left), the Scale task of the MBEA (right). The densities show the distribution of proportion correct scores in the control participants and the boxplot shows the quartiles of the control population (the whiskers show 1.5x interquartile range and points represent outliers). The dots show individual participants (for the aphasic individuals, the initials indicate the specific participant). Dashed grey lines indicate chance performance.

**Table 5.**
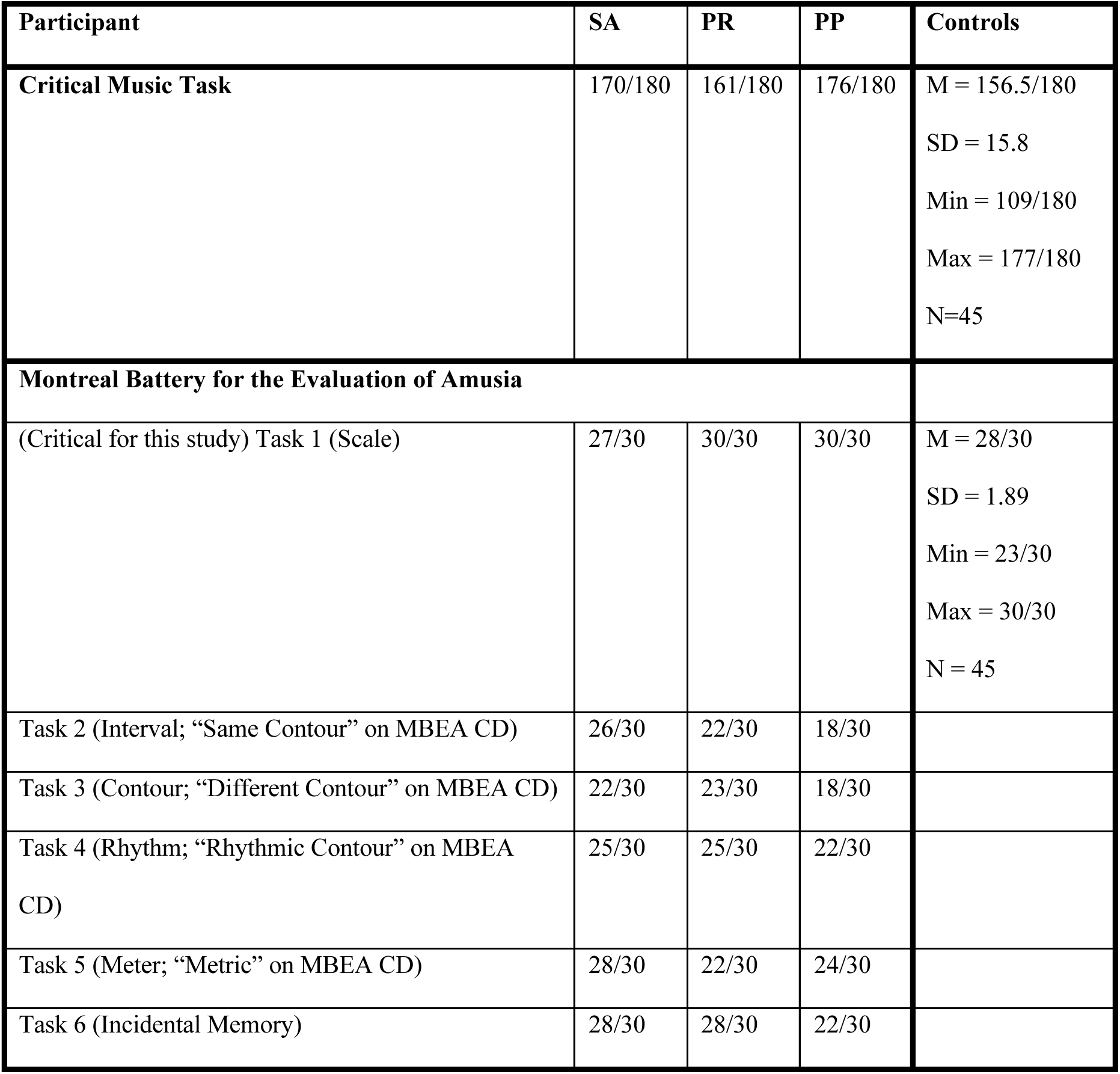
Results for participants with aphasia and control participants on the critical music task and the Scale task of the MBEA (Peretz et al., 2003). For participants with aphasia, we report the results from all six MBEA tasks, for completeness.

#### Does music elicit a response in the language network of native speakers of a tonal language?

The above analyses focus on the language network’s responses to music stimuli and its sensitivity to music structure in English native speakers. However, some have argued that responses to music may differ in speakers of languages that use pitch to make lexical or grammatical distinctions (e.g., Deutsch et al. 2006, 2009; Bidelman et al. 2011; Creel et al. 2018; Ngo et al. 2016, Liu et al. 2021). In Experiment 4, we therefore tested whether language regions of Mandarin native speakers respond to music. Similar to Experiment 1, we compared the response to the music condition against a) the fixation baseline, b) the foreign language condition, and c) a non-linguistic, non-music condition (environmental sounds). A brain region that supports music processing should respond more strongly to music than the fixation baseline and the foreign condition; if the response is further selective, it should be stronger than the response elicited by environmental sounds.

Results from Mandarin native speakers replicated the results from Experiment 1: the music condition did not elicit a strong response in the language network (**Figure 6**; **Table 6**). Although the response to music was above the fixation baseline at the network level and in some fROIs, the response did not differ from (or was lower than) the responses elicited by an unfamiliar foreign language (Russian) and environmental sounds.

**Figure 6.**
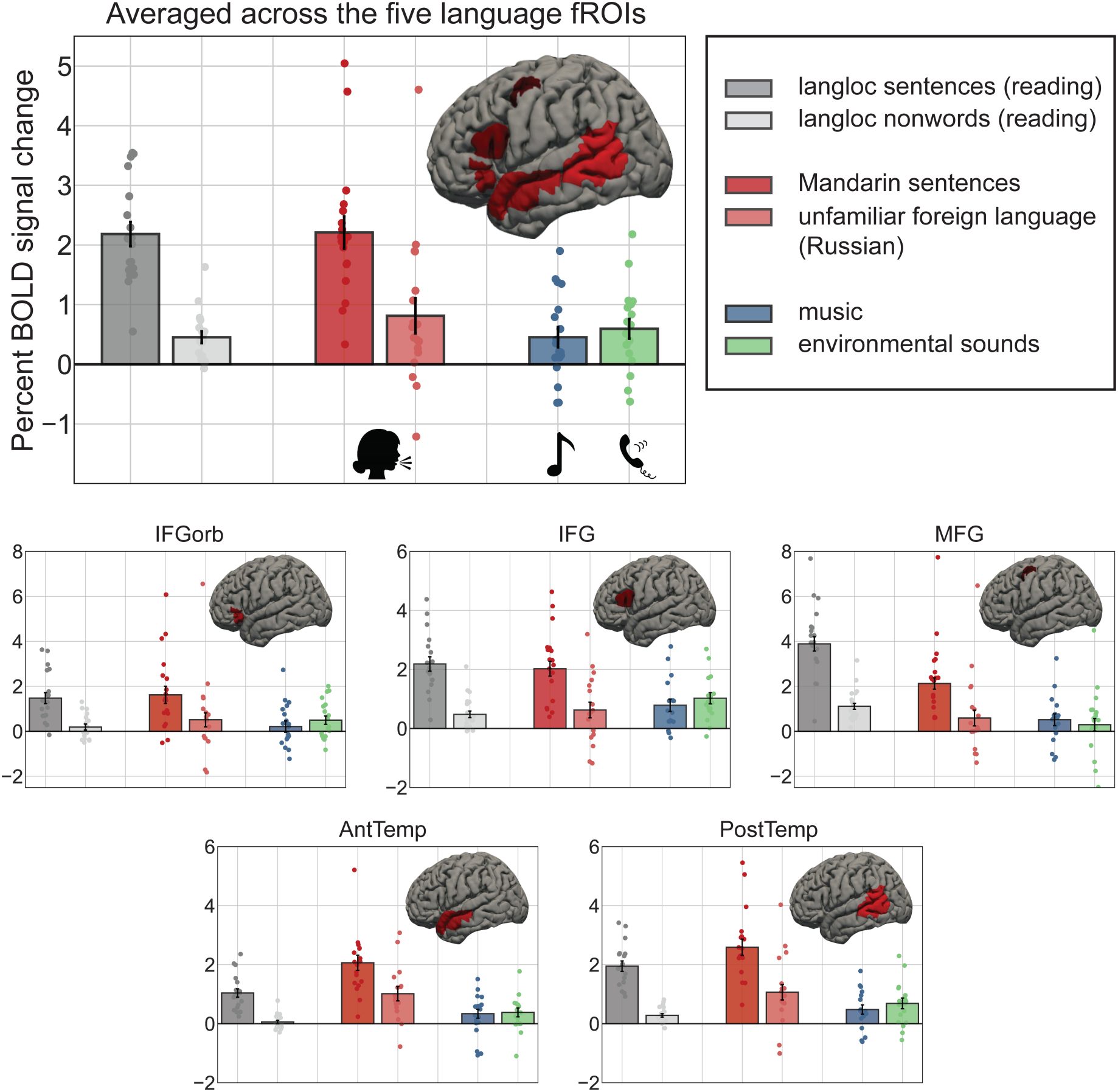
Responses of the language fROIs (pooling across the network – top, and for each fROI individually – bottom) to the language localizer conditions (in grey), to the two auditory conditions containing speech (red shades), to the music condition (blue), and to the non-linguistic/non-music auditory condition (green) in Experiment 4. The error bars represent standard error of the mean by participants. The response to the music condition is much lower than the response to sentences, and is not higher than the response to foreign language and environmental sounds.

**Table 6.** Statistical results (two-sided) for the contrasts between the music condition and fixation, foreign language, and environmental sounds in Experiment 4. Abbreviations: b=the beta estimate for the effect; se=standard error of the mean by participants; df=degrees of freedom; d=Cohen’s d (Westfall et al. 2014; Brysbaert and Stevens 2018); t=the t statistic; p=the significance value (for the individual fROIs, these values have been FDR-corrected for the number of fROIs (n=5)). In light grey, we highlight the results that are **not consistent** with the role of the language regions in music perception: of the 18 tests performed, 3 showed an effect predicted by language-music overlap accounts: a small positive response to the music condition relative to the weakest baseline (fixation) at the network level and in two fROIs individually; this response was still ∼2 lower than the unfamiliar foreign language condition and was numerically lower than the environmental sounds condition.

| Contrast | Language<br>network | LIFGorb | LIFG | LMFG | LAnt<br>Temp | LPost<br>Temp |
| --- | --- | --- | --- | --- | --- | --- |
| music<br>>fixation | $\beta=0.454$<br>se=0.177<br>df=17.646<br>d=0.517<br>t=2.565<br>p=0.020* | $\beta=0.299$<br>se=0.228<br>df=nan<br>d=nan<br>t=1.308<br>p=1.000 | $\beta=0.761$<br>se=0.207<br>df=nan<br>d=nan<br>t=3.683<br>p=0.010* | $\beta=0.480$<br>se=0.260<br>df=nan<br>d=nan<br>t=1.848<br>p=0.410 | $\beta=0.268$<br>se=0.171<br>df=nan<br>d=nan<br>t=1.568<br>p=0.675 | $\beta=0.462$<br>se=0.156<br>df=nan<br>d=nan<br>t=2.962<br>p=0.045* |
| music<br>>foreign | $\beta=-0.359$<br>se=0.141<br>df=162.000<br>d=-0.308<br>t=-2.547<br>p=0.012* | $\beta=-0.360$<br>se=0.416<br>df=18.000<br>d=-0.258<br>t=-0.865<br>p=1.000 | $\beta=0.123$<br>se=0.309<br>df=18.000<br>d=0.124<br>t=0.398<br>p=1.000 | $\beta=-0.219$<br>se=0.473<br>df=18.000<br>d=-0.149<br>t=-0.463<br>p=1.000 | $\beta=-0.703$<br>se=0.240<br>df=18.000 d=-<br>0.870<br>t=-2.926<br>p=0.045* | $\beta=-0.638$<br>se=0.254<br>df=18.000<br>d=-0.686<br>t=-2.511<br>p=0.110 |
| music<br>>environmental<br>sounds | $\beta=-0.141$<br>se=0.108<br>df=157.749<br>d=-0.154 t=-<br>1.299<br>p=0.196 | $\beta=-0.249$<br>se=0.187<br>df=18.000<br>d=-0.280 t=-<br>1.328<br>p=1.000 | $\beta=-0.240$<br>se=0.193<br>df=18.000<br>d=-0.302 t=-<br>1.248<br>p=1.000 | $\beta=0.038$<br>se=0.304<br>df=18.000<br>d=0.030<br>t=0.125<br>p=1.000 | $\beta=-0.042$<br>se=0.147<br>df=18.000 d=-<br>0.065<br>t=-0.285<br>p=1.000 | $\beta=-0.210$<br>se=0.179<br>df=18.000<br>d=-0.310<br>t=-1.171<br>p=1.000 |

## Discussion

We here tackled a much investigated but still debated question: do the brain regions of the language network support the processing of music, especially music structure? Across three fMRI experiments, we obtained a clear answer: the brain regions of the language network, which support the processing of linguistic syntax (e.g., Fedorenko et al. 2010, 2020; Pallier et al. 2011; Bautista and Wilson 2016; Blank et al. 2016), do not support music processing (see **Table 7** for a summary of the results). We found overall low responses to music (including orchestral pieces, solo pieces played on different instruments, synthetic music, and vocal music) in the language brain regions (**Figure 3**; see Sueoka et al. 2022, for complementary evidence from the inter-subject correlation approach applied to a rich naturalistic music stimulus), including in speakers of a tonal language (**Figure 6**), and no consistent sensitivity to manipulations of music structure (**Figure 4**). We further found that the ability to make well-formedness judgments about the tonal structure of music was preserved in patients with severe aphasia who cannot make grammaticality judgments for sentences (**Figure 5**), although we acknowledge the possibility that general ability to detect unexpected events may have contributed to performance on the critical music-structure tasks (e.g., Bigand et al. 2014; Collins et al. 2014) and that additional controls would be needed to conclusively determine whether these patients have preserved music-structure processing abilities. Nevertheless, given the brain imaging results (summarized in **Table 7**), a critical role of the language system in music structure processing is unlikely.

**Table 7.**
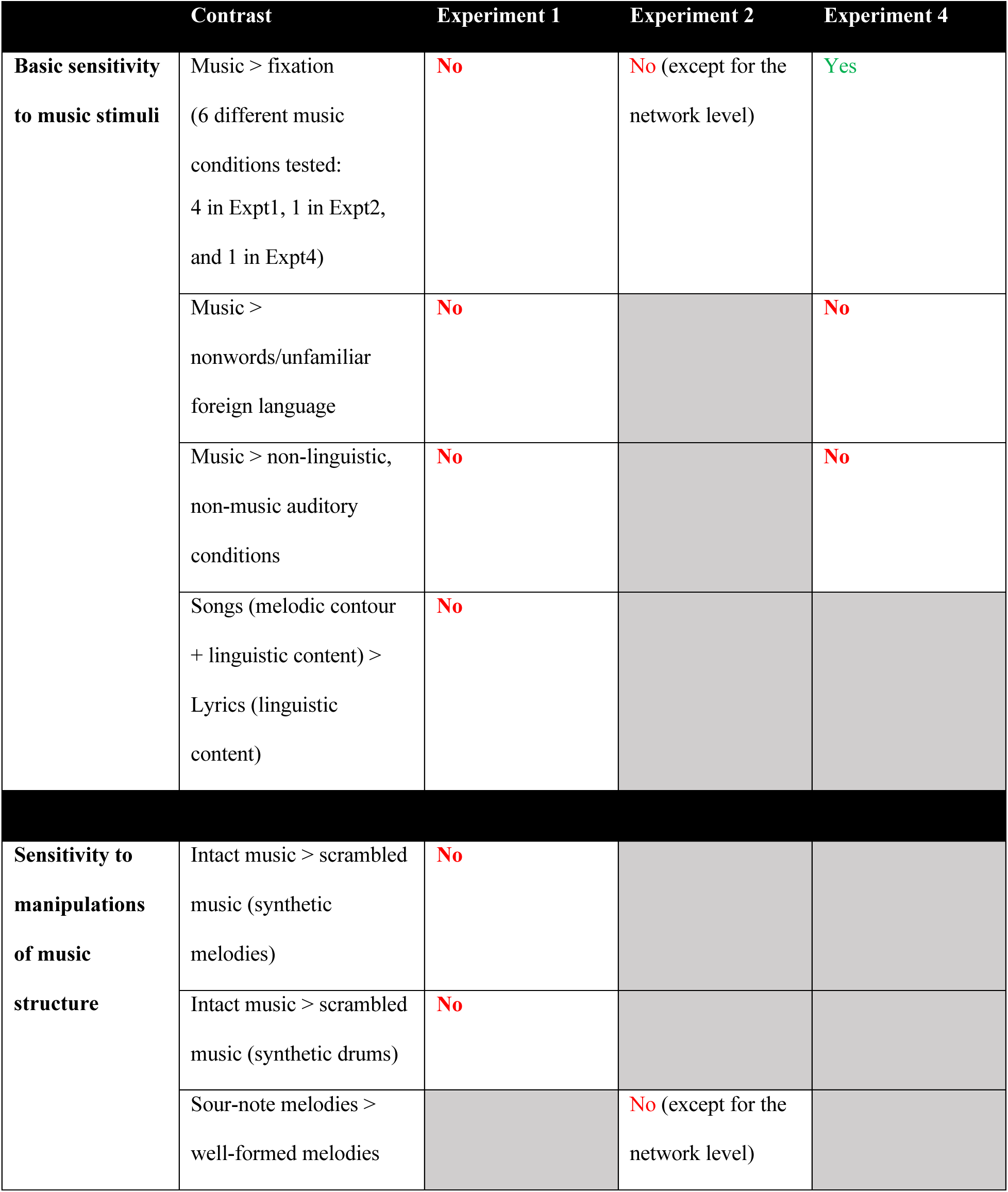
A summary of the results for the tests of the language network’s sensitivity to music in general and to music structure specifically. This pattern of results constitutes strong evidence against the role of the language system—or any of its components—in music perception, including the processing of music structure. With respect to sensitivity to music stimuli: 4 of the 6 conditions failed to elicit a response above the low-level (fixation) baseline anywhere in the language network; 1 condition (in Experiment 2) elicited a small and weakly significant above-fixation response at the network level only (not in any individual fROIs); and 1 condition (in Experiment 4) elicited a small above-fixation response (including in two individual fROIs) but this response was not higher than that elicited by other auditory conditions like environmental sounds. With respect to sensitivity to music structure: 2 of the 3 manipulations failed to elicit a response anywhere in the language network, and the remaining manipulation elicited a small and weakly significant effect at the network level, which was not reliable in any individual ROI.

Our findings align with a) prior neuropsychological patient evidence of language/music dissociations (e.g., Luria et al. 1965; Brust 1980; Marin 1982; Basso and Capitani 1985; Polk and Kertesz 1993; Peretz et al. 1994, 1997; Piccirilli et al. 2000; Peretz and Coltheart 2003; Slevc et al. 2016; Faroqi-Shah et al. 2020; Chiapetta et al. 2022) and with b) prior evidence that music is processed by music-selective areas in the auditory cortex (Norman-Haignere et al. (2015; see also Boebinger et al. 2021; see Peretz et al. 2015, for review and discussion). The latter, music-selective areas are strongly sensitive to the scrambling of music structure in stimuli like those used here in Experiment 1 (see also Fedorenko et al. 2012c, Boebinger 2021; see Mehr et al. 2019 for *a priori* reasons to expect the effects of tonal structure manipulations in music-selective brain regions). (We provide the responses of music-responsive areas to the conditions of Experiments 1 and 2 at: https://osf.io/68y7c/).) In contrast, our findings stand in sharp contrast to numerous reports arguing for shared structure processing mechanisms in the two domains, including specifically in the inferior frontal cortex, within ‘Broca’s area’ (e.g., Patel et al. 1998; Koelsch et al. 2000; Maess et al. 2001; Koelsch et al. 2002; Levitin and Menon 2003; see Kunert and Slevc 2015; LaCroix et al. 2016; Vuust et al. 2022 for reviews).

Below, we discuss several issues that are relevant for interpreting the current results and/or that these results inform, and outline some limitations of scope of our study.

### 1. Theoretical considerations about the language-music relationship

Why might we *a priori* think that the language network, or some of its components, may be important for processing music in general, or for processing music structure specifically? Similarities between language and music have long been noted and discussed. For example, as summarized in Jackendoff (2009; see also Patel 2008), both capacities are human-specific, involve the production of sound (though this is not always the case for language: cf. sign languages, or written language in literate societies), and have multiple culture-specific variants. Furthermore, language and music are intertwined in songs, which appear to be a cultural universal (e.g., Brown 1991; Nettl 2015; see Mehr et al. 2019 for empirical support; see Norman-Haignere et al. 2021 for evidence of neural selectivity for songs in the auditory cortex). However, Jackendoff (2009) notes that i) most cognitive capacities / mechanisms that have been argued to be common to language and music are not *uniquely* shared by language and music, and ii) language and music differ in several critical ways, and these differences are important to consider alongside potential similarities when theorizing about possible shared representations and computations.

To elaborate on the first point: the cognitive capacity that has perhaps received the most attention in discussions of cognitive and neural mechanisms that may be shared by language and music is the combinatorial capacity of the two domains (e.g., Riemann 1877, as cited in Swain 1995; Lindblom and Sundberg 1969; Fay 1971; Sundberg and Lindblom 1976; Lerdahl and Jackendoff 1977, 1983; Roads 1979; Krumhansl and Keil 1982). In particular, in language, words can be combined into complex hierarchical structures to form novel phrases and sentences, and in music, notes and chords can similarly be combined to form novel melodies. Further, in both domains, the combinatorial process is constrained by a set of conventions. However, this capacity can be observed, in some form, in many other domains, from visual processing, to math, to social cognition, to motor planning, to general reasoning. Similarly, other cognitive capacities that are necessary to process language and music—including a large long-term memory store for previously encountered elements and patterns, a working memory capacity needed to integrate information as it comes in, an ability to form expectations about upcoming elements, and an ability to engage in joint action—are important for information processing in other domains. An observation that some mental capacity is necessary for multiple domains is compatible with at least two architectures: one where the relevant capacity is implemented (perhaps in a similar way) in each relevant set of domain-specific circuits, and another where the relevant capacity is implemented in a centralized mechanism that all domains draw on (e.g., Fedorenko and Shain 2021). Those arguing for overlap between language and music processing advocate a version of the latter. Critically, any shared mechanism that language and music would draw on should also support information processing in other domains that require the relevant computation (see Section 3 below for arguments against this kind of architecture). (A possible exception, according to Jackendoff (2009), may be the fine-scale vocal motor control that is needed for speech and vocal music production (cf. sign language or instrumental music), but not any other behaviors, but this kind of ability is implemented outside of the core high-level language system, in the network of brain areas that support articulation (e.g., Basilakos et al. 2015; Guenter 2016).)

More importantly, aside from the similarities that have been noted between language and music, numerous differences characterize the two domains. Most notable are their different functions. Language enables humans to express propositional meanings, and thus to share thoughts with one another. The function of music has long been debated (e.g., Darwin 1871; Pinker 1994; see e.g., McDermott 2008 and Mehr et al. 2020, for a summary of key ideas), but most proposed functions have to do with emotional or affective processing, often with a social component^1^ (Jackendoff 2009; Savage et al. 2020). If function drives the organization of the brain (and biological systems more generally; e.g., Rueffler et al. 2012) by imposing particular computational demands on each domain (e.g., Mehr et al. 2020), these fundamentally different functions of language and music provide a theoretical reason to expect cognitive and neural separation between them. Besides, even the components of language and music that appear similar on the surface (e.g., combinatorial processing) differ in deep and important ways (e.g., Patel 2008; Jackendoff 2009; Slevc 2009; Temperley 2022).

### 2. Functional selectivity of the language network

The current results add to the growing body of evidence that the left-lateralized fronto-temporal brain network that supports language processing is highly selective for linguistic input (e.g., Fedorenko et al. 2011; Monti et al. 2009, 2012; Deen et al. 2015; Pritchett et al. 2018; Jouravlev et al. 2019; Ivanova et al. 2020, 2021; Benn, Ivanova et al. 2021; Liu et al. 2020; Deen and Freiwald 2021; Paunov et al. 2022; Sueoka et al. 2022; see Fedorenko and Blank 2020 for a review) and not critically needed for many forms of complex cognition (e.g., Lecours and Joanette 1980; Varley and Siegal 2000; Varley et al. 2005; Apperly et al. 2006; Woolgar et al. 2018; Ivanova et al. 2021; see Fedorenko and Varley 2016 for a review). Importantly, this selectivity holds across all components of the language network, including the parts that fall within ‘Broca’s area’ in the left inferior frontal gyrus. As discussed in the introduction, many claims about shared structure processing in language and music have focused specifically on Broca’s area (e.g., Patel 2003; Fadiga et al. 2009; Fitch and Martins 2014). The evidence presented here shows that the language-responsive parts of Broca’s area, which are robustly sensitive to linguistic syntactic manipulations (e.g., Just et al. 1996; Stromswold et al. 1996; Ben-Shachar et al. 2003; Caplan et al. 2008; Peelle et al. 2010; Blank et al. 2016; see e.g., Friederici 2011 and Hagoort and Indefrey 2014 for meta-analyses), do not respond when we listen to music and are not sensitive to structure in music. These results rule out the hypothesis that language and music processing rely on the same mechanism housed in Broca’s area.

It is also worth noting that the very *premise* of the latter hypothesis—of a special relationship between Broca’s area and the processing of linguistic syntax (e.g., Caramazza and Zurif 1976; Friederici 2018)—has been questioned and overturned. *First*, syntactic processing does not appear to be carried out focally, but is instead distributed across the entire language network, with all of its regions showing sensitivity to syntactic manipulations (e.g., Fedorenko et al. 2010, 2020; Pallier et al. 2011; Blank et al. 2016; Shain, Blank et al. 2020; Shain et al. 2022), and with damage to different components leading to similar syntactic comprehension deficits (e.g., Caplan et al. 1996; Dick et al. 2001; Wilson and Saygin 2004; Mesulam et al. 2014; Mesulam et al. 2015). And *second*, the language-responsive part of Broca’s area, like other parts of the language network, is sensitive to both syntactic processing and word meanings, and even sub-lexical structure (Fedorenko et al. 2010, 2012b, 2020; Regev et al. 2021; Shain et al. 2021). The lack of segregation between syntactic and lexico-semantic processing is in line with the idea of ‘lexicalized syntax’ where the conventions for how words can combine with one another are highly dependent on the particular lexical items (e.g., Goldberg 2002; Jackendoff 2002, 2007; Sag et al. 2003; Levin and Rappaport-Hovav 2005; Bybee 2010; Jackendoff and Audring 2020), and is contra the idea of combinatorial rules that are blind to the content/meaning of the to-be-combined elements (e.g., Chomsky 1965, 1995; Fodor 1983; Pinker and Prince 1988; Pinker 1991, 1999; Pallier et al. 2011).

### 3. Overlap in structure processing in language and music outside of the core language network?

We have here focused on the core fronto-temporal language network. Could structure processing in language and music draw on shared resources elsewhere in the brain? The prime candidate is the domain-general executive control, or Multiple Demand (MD), network (e.g., Duncan and Owen 2000; Duncan 2001, 2010; Assem et al. 2020), which supports functions like working memory and inhibitory control. Indeed, according to Patel’s Shared Structural Integration Resource Hypothesis (SSIRH; 2003, 2008, 2012), language and music draw on separate representations, stored in distinct cortical areas, but rely on the same working memory store to integrate incoming elements into evolving structures. Relatedly, Slevc et al. (2013; see Asano et al. 2021 for a related proposal) have argued that another executive resource—inhibitory control—may be required for structure processing in both language and music. Although it is certainly possible that some aspects of linguistic and/or musical processing would require domain-general executive resources, based on the available evidence from the domain of language, we would argue that any such engagement does not reflect the engagement of computations like syntactic structure building. In particular, Blank and Fedorenko (2017) found that activity in the brain regions of the domain-general MD network does not closely ‘track’ linguistic stimuli, as evidenced by low inter-subject correlations during the processing of linguistic input (see Paunov et al. 2021 and Sueoka et al. 2022 for replications). Further, Diachek, Blank, Siegelman et al. (2020) showed in a large-scale fMRI investigation that the MD network is not engaged during language processing in the absence of secondary task demands (cf. the core language network, which is relatively insensitive to task demands and responds robustly even during passive listening/reading). And Shain, Blank et al. (2020; also, Shain et al. 2022) have shown that the language network, but not the MD network, is sensitive to linguistic surprisal and working-memory integration costs (see also Wehbe et al. 2021 for evidence that activity in the language, but not the MD, network reflects general incremental processing difficulty).

In tandem, this evidence argues against the role of executive resources in core linguistic computations like those related to lexical access and combinatorial processing, including syntactic parsing and semantic composition (see also Hasson et al. 2015 and Dasgupta and Gershman 2021 for general arguments against the separation between memory and computation in the brain). Thus, although the contribution of executive resources to music processing deserves further investigation (cf. https://osf.io/68y7c/ for evidence of low responses of the MD network to the music conditions in the current study), any overlap within the executive system between linguistic and music processing cannot reflect core linguistic computations, as those seem to be carried out by the language network (see Fedorenko and Shain 2021, for a review). Functionally identifying the MD network in individual participants (e.g., Fedorenko et al. 2013; Shashidhara et al. 2019) is a powerful way to help interpret the observed effects of music manipulations as reflecting general executive demands (see Saxe et al. 2006, Blank et al. 2017 and Fedorenko 2021, for general discussions of greater interpretability of fMRI results obtained from the functional localization approach). Importantly, given the ubiquitous sensitivity of the MD network to cognitive demands, it is / will be important to rule out task demands, rather than stimulus processing, as the source of overlap between music and language processing in interpreting past studies and designing future ones.

### 4. Overlap between music processing and other aspects of speech / language

The current study investigated the role of the language network—which supports ‘high-level’ comprehension and production—in music processing. As a result, the claims we make are restricted to those aspects of language that are supported by this network. These include the processing of word meanings and combinatorial (syntactic and semantic) processing, but exclude speech perception, prosodic processing, higher-level discourse structure building, and at least some aspects of pragmatic reasoning. Some of these components of language (e.g., pragmatic reasoning) seem *a priori* unlikely to share resources with music. Others (e.g., speech perception) have been shown to robustly dissociate from music (Norman-Haignere et al. 2015; Overath et al. 2015; Kell et al. 2018; Boebinger et al. 2021). However, some components of speech and language may, and some do, draw on the same resources as aspects of music. For example, aspects of pitch perception have been argued to overlap between speech and music based on behavioral and neuropsychological evidence (e.g., Wong and Perrachione 2007; Perrachione et al. 2013; Patel et al. 2008b). Indeed, brain regions that selectively respond to different kinds of pitched sounds have been previously reported (Patterson et al. 2002; Penagos et al. 2004; Norman-Haignere et al. 2013, 2015). Some studies have also suggested that music training may improve general rapid auditory processing and pitch encoding that are important for speech perception and language comprehension (e.g., Overy 2003; Tallal and Gaab 2006; Wong et al. 2007), although at least some of these effects likely originate in the brainstem and subcortical auditory regions (e.g., Wong et al. 2007). Other aspects of high-level auditory perception, including aspects of rhythm, may turn out to overlap as well, and deserve further investigation (see Patel 2008, for a review).

We also have focused on Western tonal instrumental music here. In the future, it would be useful to extend these findings to more diverse kinds of music. That said, given that individuals are most sensitive to structure in music with which they have experience (e.g., Cuddy et al. 1981; Cohen 1982; Curtis and Barucha 2009), it seems unlikely that music from less familiar traditions would elicit a strong response in the language areas (see Boebinger 2021, for evidence that music-selective areas of the auditory cortex respond to culturally diverse music styles). Further, given that evolutionarily early forms of music were likely vocal (e.g., Trehub 2003; Mehr 2017), it would be useful to examine the responses of the language regions to vocal music without linguistic content, like humming or whistling. Based on preliminary unpublished data from our lab (available upon request), responses to such stimuli in the language areas appear low.

In conclusion, we have here provided extensive evidence against the role of the language network in music perception, including the processing of music structure. Although the relationship between music and aspects of speech and language will likely continue to generate interest in the research community, and aspects of speech and language other than those implemented in the core fronto-temporal language-selective network (Fedorenko et al. 2011; Fedorenko and Thompson-Schill 2014) may indeed share some processing resources with (aspects of) music, we hope that the current study helps bring clarity to the debate about structure processing in language and music.

## Data availability

The datasets generated during and/or analyzed during the current study are available in the OSF repository: https://osf.io/68y7c/.

## Code availability

Scripts for statistical analysis are available at: https://osf.io/68y7c/.

## Acknowledgements

We would like to acknowledge the Athinoula A. Martinos Imaging Center at the McGovern Institute for Brain Research at MIT, and its support team (Steve Shannon and Atsushi Takahashi). We thank former and current EvLab members for their help with fMRI data collection (especially Meilin Zhan for help with Experiment 4). We thank Josh McDermott for input on many aspects of this work, Jason Rosenberg for composing the melodies used in Experiments 2 and 3, and Zuzanna Balewski for help with creating the final materials used in Experiments 2 and 3. For Experiment 3, we thank Vitor Zimmerer for help with creating the grammaticality judgment task, Ted Gibson for help with collecting the control data, and Anya Ivanova for help with Figure 2. For Experiment 4, we thank Anne Cutler, Peter Graff, Morris Alper, Xiaoming Wang, Taibo Li, Terri Scott, Jeanne Gallée, and Lauren Clemens for help with constructing and/or recording and/or editing the language materials, and Fatemeh Khalilifar, Caitlyn Hoeflin, and Walid Bendris for help with selecting the music materials and with the experimental script. Finally, we thank the audience at the Society for Neuroscience conference (2014), the Neurobiology of Language conference (virtual edition, 2020), Ray Jackendoff, Dana Boebinger, and members of the Fedorenko and Gibson labs for helpful comments and discussions. RR was supported by NIH award F32-DC-015163. SNH was supported by a graduate NSF award, as well as postdoctoral awards from the HHMI / Life Sciences research foundation and a K99/R00 award from the NIH (1K99DC018051-01A1). SMM was supported by a La Caixa fellowship LCF/BQ/AA17/11610043. RV was supported by Alzheimer’s Society and The Stroke Association. EF was supported by the R00 award HD057522, R01 awards DC016607, DC016950, and NS121471, by the Paul and Lilah Newton Brain Science Award, and funds from the Brain and Cognitive Sciences department, the McGovern Institute for Brain Research, and the Simons Center for the Social Brain.

## Author contributions

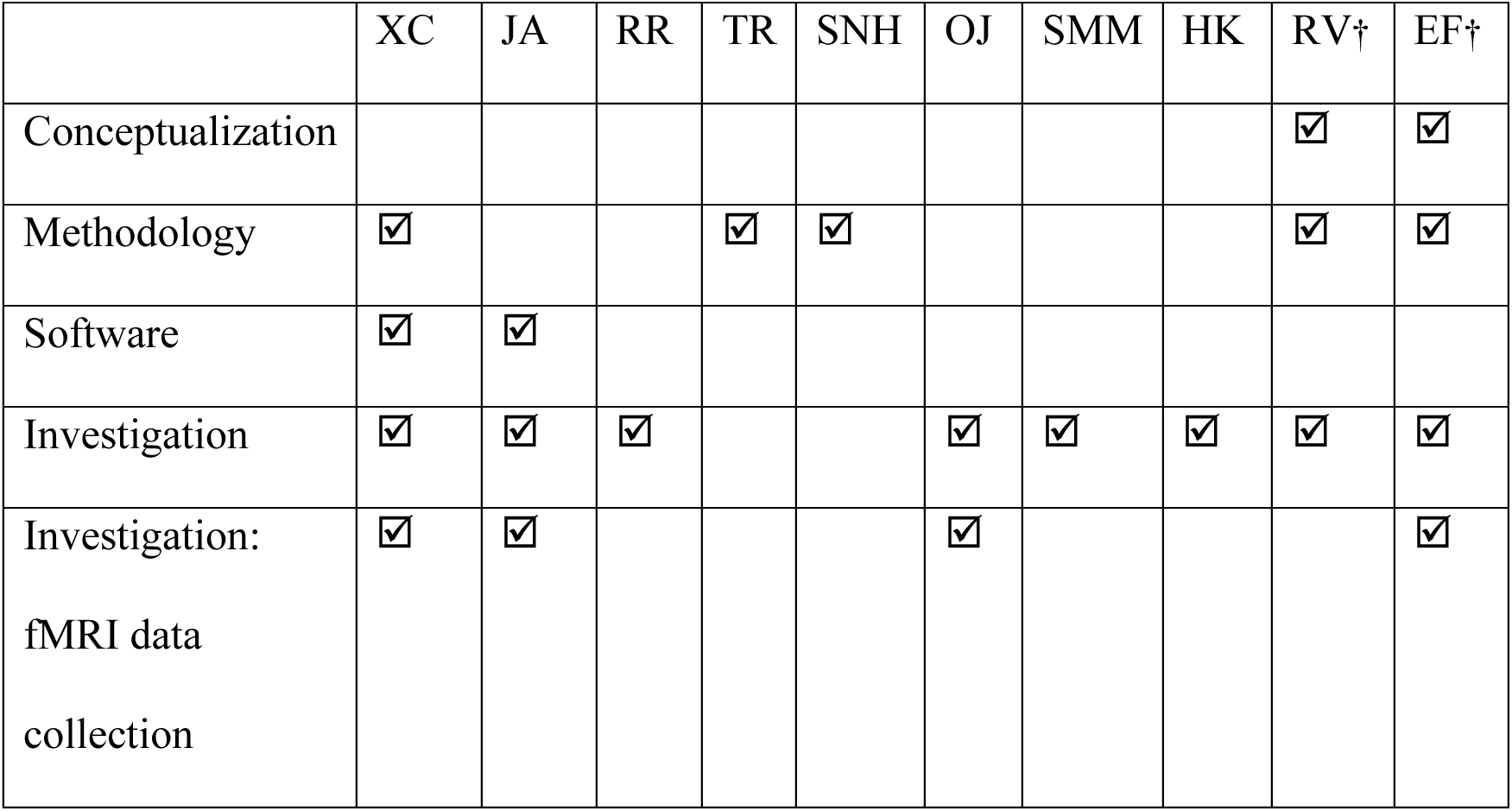
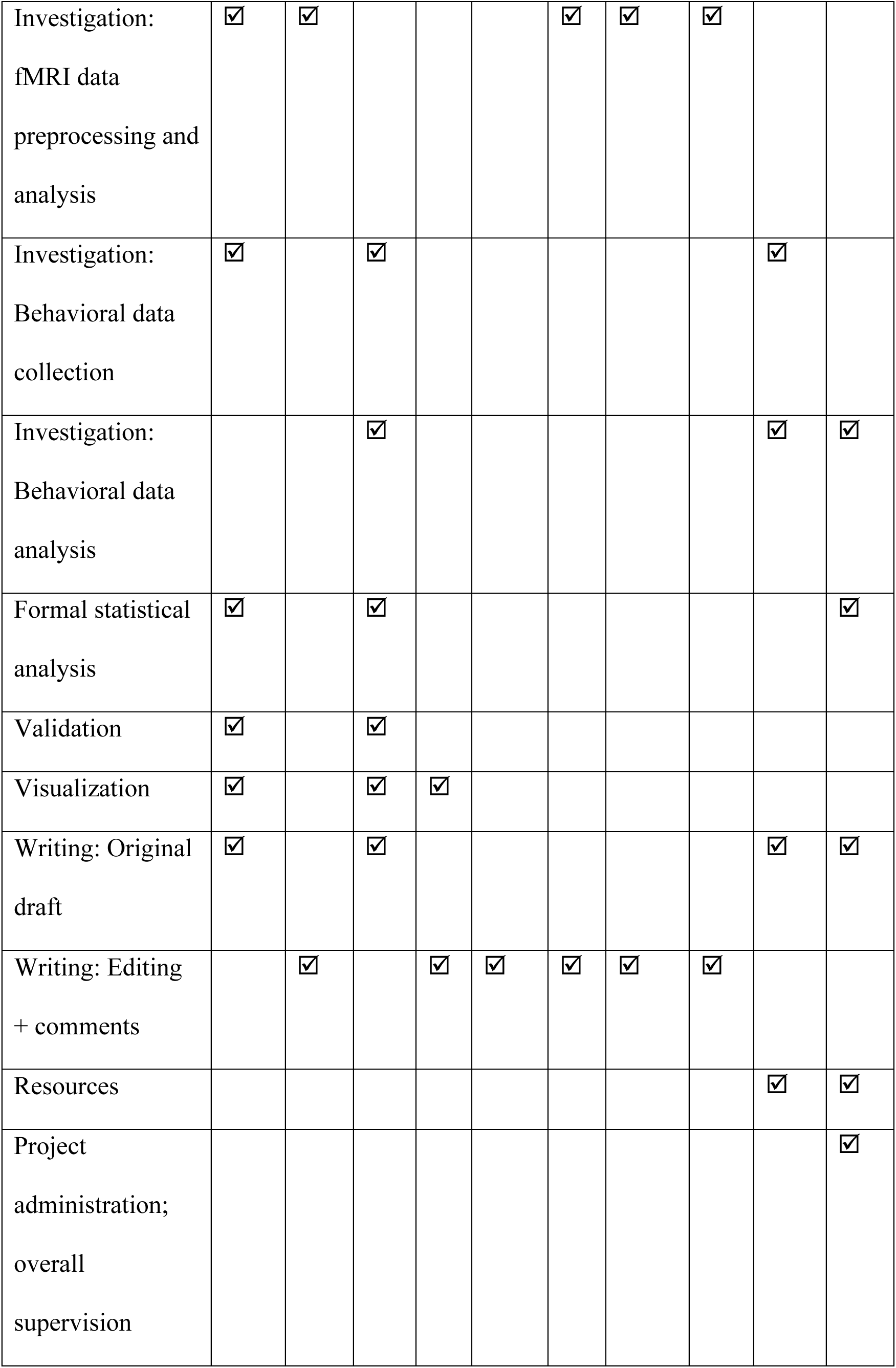

## Footnotes

1 Although some have discussed the notions of ‘meaning’ in music (e.g., Meyer 1961; Raffman 1993; Cross and Tolbert 2009; Koelsch 2001), it is uncontroversial that music cannot be used to express propositional thought (for discussion, see Patel 2008; Jackendoff 2009; Slevc 2009).

## References

Alcock KJ, Wade D, Anslow P, Passingham RE. 2000. Pitch and timing abilities in adult left-hemisphere-dysphasic and right-hemisphere-damaged subjects. Brain Lang. 75(1):47–65.

Amalric M, Dehaene S. 2018. Cortical circuits for mathematical knowledge: evidence for a major subdivision within the brain’s semantic networks. Philos Trans R Soc Lond B Biol Sci. 373(1740):20160515.

Apperly IA, Samson D, Carroll N, Hussain S, Humphreys G. 2006. Intact first-and second-order false belief reasoning in a patient with severely impaired grammar. Soc Neurosci. 1(3-4):334–348.

Asano R, Boeckx C, Seifert U. 2021. Hierarchical control as a shared neurocognitive mechanism for language and music. Cognition. 216:104847.

Assem M, Glasser MF, Van Essen DC, Duncan J. 2020. A domain-general cognitive core defined in multimodally parcellated human cortex. Cereb Cortex. 30(8):4361–4380.

Baillet S. 2014. Forward and Inverse Problems of MEG/EEG. In: Jaeger D, Jung R, editors. Encyclopedia of Computational Neuroscience. New York (NY): Springer. p.1–8.

Baroni M, Maguire S, Drabkin W. 1983. The concept of musical grammar. Music Anal, 2(2):175–208.

Basilakos A, Rorden C, Bonilha L, Moser D, Fridriksson J. 2015. Patterns of poststroke brain damage that predict speech production errors in apraxia of speech and aphasia dissociate. Stroke. 46(6):1561–1566.

Basso A, Capitani E. 1985. Spared musical abilities in a conductor with global aphasia and ideomotor apraxia. J Neurol Neurosurg Psychiatry. 48(5):407–412.

Bates D, Mächler M, Bolker B, Walker S. 2015. Fitting linear mixed-effects models using lme4. J Stat Softw. 67(1):1–48.

Bautista A, Wilson SM. 2016. Neural responses to grammatically and lexically degraded speech. Lang Cogn Neurosci. 31(4):567–574.

Ben-Shachar M, Hendler T, Kahn I, Ben-Bashat D, Grodzinsky Y. 2003. The neural reality of syntactic transformations: Evidence from functional magnetic resonance imaging. Psychol Sci. 14(5):433–440.

Benn Y*, Ivanova A*, Clark O, Mineroff Z, Seikus C, Santos Silva J, Varley R^, Fedorenko E^. 2021. No evidence for a special role of language in feature-based categorization. bioRxiv.

Bernstein L. 1976. The unanswered question: Six talks at Harvard. Cambridge (MA): Harvard University Press.

Bidelman GM, Gandour JT, Krishnan A. 2011. Musicians and tone-language speakers share enhanced brainstem encoding but not perceptual benefits for musical pitch. Brain Cogn. 77(1):1–10.

Bigand E, Tillmann B, Poulin B, D’Adamo DA, Madurell F. 2001. The effect of harmonic context on phoneme monitoring in vocal music. Cognition. 81(1):B11–B20.

Bigand E, Delbé C, Poulin-Charronnat B, Leman M, Tillmann B. 2014. Empirical evidence for musical syntax processing? Computer simulations reveal the contribution of auditory short-term memory. Front Syst Neurosci. 8:94.

Bishop DVM, Norbury CF. 2002. Exploring the borderlands of autistic disorder and specific language impairment: a study using standardised diagnostic instruments. J Child Psychol Psychiatry. 43(7):917–929.

Blank I, Kanwisher N, Fedorenko E. 2014. A functional dissociation between language and multiple-demand systems revealed in patterns of BOLD signal fluctuations. J Neurophysiol. 112(5):1105–1118.

Blank I, Balewski Z, Mahowald K, Fedorenko E. 2016. Syntactic processing is distributed across the language system. Neuroimage. 127:307–323.

Blank IA, Fedorenko E. 2017. Domain-general brain regions do not track linguistic input as closely as language-selective regions. J Neurosci. 37(41):9999–10011.

Blank IA, Kiran S, Fedorenko E. 2017. Can neuroimaging help aphasia researchers? Addressing generalizability, variability, and interpretability. Cogn Neuropsychol, 34(6):377–393.

Boebinger D, Norman-Haignere SV, McDermott JH, Kanwisher N. 2021. Music-selective neural populations arise without musical training. J Neurophysiol, 125(6):2237–2263.

Boilès CL. 1973. Reconstruction of proto-melody. Anuario Interamericano de Investigacion Musical, 9:45–63.

Bortolini U, Leonard LB, Caselli MC. 1998. Specific Language Impairment in Italian and English: evaluating alternative accounts of grammatical deficits. Lang Cogn Process. 13(1):1–20.

Braga RM, DiNicola LM, Becker HC, Buckner RL. 2020. Situating the left-lateralized language network in the broader organization of multiple specialized large-scale distributed networks. J Neurophysiol. 124(5):1415–1448.

Brown DR. 1991. Human Universals. Philadelphia (PA):Temple University Press.

Brust JC. 1980. Music and language: musical alexia and agraphia. Brain. 103(2):367–392.

Brysbaert M, Stevens M. 2018. Power analysis and effect size in mixed effects models: A tutorial. J Cogn. 1(1):9.

Bybee J. 2010. Language, usage and cognition. Cambridge (UK): Cambridge University Press.

Caplan D, Hildebrandt N, Makris N. 1996. Location of lesions in stroke patients with deficits in syntactic processing in sentence comprehension. Brain. 119(3):933–949.

Caplan D, Stanczak L, Waters G. 2008. Syntactic and thematic constraint effects on blood oxygenation level dependent signal correlates of comprehension of relative clauses. J Cogn Neurosci. 20(4):643–656.

Caramazza A, Zurif EB. 1976. Dissociation of algorithmic and heuristic processes in language comprehension: Evidence from aphasia. Brain Lang. 3(4):572–582.

Chen G, Taylor PA, Cox RW. 2017. Is the statistic value all we should care about in neuroimaging?. Neuroimage. 147:952–959.

Chiappetta B, Patel AD, Thompson CK. 2022. Musical and linguistic syntactic processing in agrammatic aphasia: An ERP study. J Neurolinguistics. 62:101043.

Chomsky N. 1965. Aspects of the Theory of Syntax. Cambridge (MA): MIT press.

Chomsky N. 1995. The minimalist program. Cambridge (MA): MIT Press.

Collins T, Tillmann B, Barrett FS, Delbé C, Janata P. 2014. A combined model of sensory and cognitive representations underlying tonal expectations in music: from audio signals to behavior. Psychol Rev, 121(1):33.

Cooke A, Grossman M, DeVita C, Gonzalez-Atavales J, Moore P, Chen W, Gee J, Detre J. 2006. Large-scale neural network for sentence processing. Brain Lang, 96(1):14–36.

Cooper R. 1973. Propositions pour un modele transformationnel de description musicale. Musique en Jeu. 10:70–88.

Corbetta M, Shulman GL. 2002. Control of goal-directed and stimulus-driven attention in the brain. Nat Rev Neurosci. 3(3):201–215.

Corlett PR, Mollick JA, Kober H. 2021. Substrates of Human Prediction Error for Incentives, Perception, Cognition, and Action. psyarxiv.

Crawford JR, Garthwaite PH. 2007. Comparison of a single case to a control or normative sample in neuropsychology: Development of a Bayesian approach. Cogn Neuropsychol. 24(4), 343–372.

Creel SC, Weng M, Fu G, Heyman GD, Lee K. 2018. Speaking a tone language enhances musical pitch perception in 3–5-year-olds. Dev Sci. 21(1): e12503.

Crump MJ, McDonnell JV, Gureckis TM. 2013. Evaluating Amazon’s Mechanical Turk as a tool for experimental behavioral research. PLoS One. 8(3):e57410.

Cohen AJ. 1982. Exploring the sensitivity to structure in music. Can Univ Music Rev. 3:15–30.

Cuddy LL, Cohen AI, Mewhort DJK. 1981. Perception of structure in short melodic sequences. J Exp Psychol Hum Percept Perform. 7:869–883.

Cumming G. 2012. Understanding the new statistics: Effect sizes, confidence intervals, and meta-analysis. New York (NY): Taylor & Francis.

Curtis ME, Bharucha JJ. 2009. Memory and musical expectation for tones in cultural context. Music Percept. 26:365–375.

Dale AM. 1999. Optimal experimental design for event-related fMRI. Hum Brain Mapp. 8(2-3):109–114.

Darwin C. 1871. The Descent of Man, and Selection in Relation to Sex. London (UK): John Murray.

Dasgupta I, Gershman SJ. 2021. Memory as a Computational Resource. Trends Cogn Sci, 25(3):240–251.

Deen B, Koldewyn K, Kanwisher N, Saxe R. 2015. Functional organization of social perception and cognition in the superior temporal sulcus. Cereb Cortex. 25(11):4596–4609.

Deen B, Freiwald WA. 2021. Parallel systems for social and spatial reasoning within the cortical apex. bioRxiv.

Deutsch D, Henthorn T, Marvin E, Xu H. 2006. Absolute pitch among American and Chinese conservatory students: Prevalence differences, and evidence for a speech-related critical period. J Acoust Soc Am. 119(2):719–722.

Deutsch D, Dooley K, Henthorn T, Head B. 2009. Absolute pitch among students in an American music conservatory: Association with tone language fluency. J Acoust Soc Am. 125(4):2398–2403.

Diachek E*, Blank I*, Siegelman M*, Affourtit J, Fedorenko E. 2020. The domain-general multiple demand (MD) network does not support core aspects of language comprehension: a large-scale fMRI investigation. J Neurosci. 40(23):4536–4550.

Dick F, Bates E, Wulfeck B, Utman JA, Dronkers N, Gernsbacher MA. 2001. Language deficits, localization, and grammar: evidence for a distributive model of language breakdown in aphasic patients and neurologically intact individuals. Psychol Rev, 108(4):759–788.

Ding J, Martin RC, Hamilton AC, Schnur TT. (2020). Dissociation between frontal and temporal-parietal contributions to connected speech in acute stroke. Brain. 143(3), 862–876.

Duncan J, Owen AM. 2000. Common regions of the human frontal lobe recruited by diverse cognitive demands. Trends Neurosci. 23(10):475–483.

Duncan J. 2001. An adaptive coding model of neural function in prefrontal cortex. Nat Rev Neurosci. 2(11):820–829.

Duncan J. 2010. The multiple-demand (MD) system of the primate brain: mental programs for intelligent behaviour. Trends Cogn Sci. 14(4):172–179.

Duncan J. 2013. The structure of cognition: attentional episodes in mind and brain. Neuron. 80(1):35–50.

Embick D, Marantz A, Miyashita Y, O’Neil W, Sakai KL. 2000. A syntactic specialization for Broca’s area. Proc Natl Acad Sci USA, 97(11):6150–6154.

Fadiga L, Craighero L, D’Ausilio A. 2009. Broca’s area in language, action, and music. Ann N Y Acad Sci. 1169(1):448–458.

Fancourt A. 2013. Exploring musical cognition in children with Specific Language Impairment. Doctoral thesis, Goldsmiths, University of London.

Faroqi-Shah Y, Slevc LR, Saxena S, Fisher SJ, Pifer M. 2020. Relationship between musical and language abilities in post-stroke aphasia. Aphasiology. 34(7):793–819.

Fay T. 1971. Perceived hierarchic structure in language and music. J Music Theory. 15(1/2):112–137.

Fedorenko E, Patel A, Casasanto D, Winawer J, Gibson E. 2009. Structural integration in language and music: Evidence for a shared system. Mem Cognit, 37(1):1–9.

Fedorenko E, Hsieh P-J, Nieto-Castañon A, Whitfield-Gabrieli S, Kanwisher N. 2010. A new method for fMRI investigations of language: Defining ROIs functionally in individual subjects. J Neurophysiol. 104(2):1177–94.

Fedorenko E, Behr M, Kanwisher N. 2011. Functional specificity for high-level linguistic processing in the human brain. Proc Natl Acad Sci USA. 108(39):16428–16433.

Fedorenko E, Duncan J, Kanwisher N. 2012a. Language-selective and domain-general regions lie side by side within Broca’s area. Curr Biol. 22(21):2059–2062.

Fedorenko E, Nieto-Castañon A, Kanwisher N. 2012b. Lexical and syntactic representations in the brain: An fMRI investigation with multi-voxel pattern analyses. Neuropsychologia. 50(4):499–513.

Fedorenko E, McDermott J, Norman-Haignere S, Kanwisher N. 2012c. Sensitivity to musical structure in the human brain. J Neurophysiol. 108(12):3289–3300.

Fedorenko E, Duncan J, Kanwisher N. 2013. Broad domain-generality in focal regions of frontal and parietal cortex. Proc Natl Acad Sci USA. 110(41):16616–16621.

Fedorenko E. 2014. The role of domain-general cognitive control in language comprehension. Front Psychol. 5:335.

Fedorenko E, Thompson-Schill SL. 2014. Reworking the language network. Trends Cogn Sci. 18(3):120–126.

Fedorenko E, Varley, R. 2016. Language and thought are not the same thing: Evidence from neuroimaging and neurological patients. Ann N Y Acad Sci. 1369(1):132–153.

Fedorenko E, Blank I. 2020. Broca’s Area Is Not a Natural Kind. Trends Cogn Sci. 24(4):270–284.

Fedorenko E, Blank I, Siegelman M, Mineroff Z. 2020. Lack of selectivity for syntax relative to word meanings throughout the language network. Cognition. 203:104348.

Fedorenko E. 2020. The brain network that supports high-level language processing. In Gazzaniga M, Ivry RB, Mangun GR, editors. Cognitive Neuroscience: The Biology of the Mind (5th edition). New York (NY): WW Norton and Company.

Fedorenko E. 2021. The early origins and the growing popularity of the individual-subject analytic approach in human neuroscience. Curr Opin Behav Sci. 40:105–112.

Fedorenko E, Shain C. 2021. Similarity of computations across domains does not imply shared implementation: The case of language comprehension. Curr Dir Psychol Sci. 30(6):526–534.

Fischl B, Rajendran N, Busa E, Augustinack J, Hinds O, Yeo BT, Mohlberg H, Amunts K, Zilles K. 2008. Cortical folding patterns and predicting cytoarchitecture. Cereb Cortex. 18(8):1973–1980.

Fitch WT, Martins MD. 2014. Hierarchical processing in music, language, and action: Lashley revisited. Ann N Y Acad Sci. 1316(1):87–104.

Fodor JD. 1983. Phrase structure parsing and the island constraints. Linguist Philos. 6(2):163–223.

Fouragnan E, Retzler C, Philiastides MG. 2018. Separate neural representations of prediction error valence and surprise: Evidence from an fMRI meta-analysis. Hum Brain Mapp. 39(7):2887–2906.

Franklin S, Turner JE, Ellis AW. 1992. ADA Comprehension Battery. York (UK): University of York.

Friederici AD, Fiebach CJ, Schlesewsky M, Bornkessel ID, Von Cramon DY. 2006. Processing linguistic complexity and grammaticality in the left frontal cortex. Cereb Cortex. 16(12):1709–1717.

Friederici AD, Kotz SA, Scott SK, Obleser J. 2010. Disentangling syntax and intelligibility in auditory language comprehension. Hum Brain Mapp. 31(3):448–457.

Friederici AD. 2011. The brain basis of language processing: from structure to function. Physiol Rev. 91(4):1357–1392.

Friederici AD. 2018. The neural basis for human syntax: Broca’s area and beyond. Curr Opin Behav Sci. 21:88–92.

Frost MA, Goebel R. 2012. Measuring structural–functional correspondence: spatial variability of specialised brain regions after macro-anatomical alignment. Neuroimage 59(2):1369–1381.

Giesbrecht F, Burns J. 1985. Two-Stage Analysis Based on a Mixed Model: Large-Sample Asymptotic Theory and Small-Sample Simulation Results. Biometrics. 41(2):477–486.

Goldberg AE. 2002. Construction Grammar. In: Nadel L, editor. Encyclopedia of Cognitive Science. Stuttgart (Germany): Macmillan.

Green DM, Swets JA. 1966. Signal detection theory and psychophysics. New York (NY): Wiley.

Guenther FH. 2016. Neural control of speech. Cambridge (MA): MIT Press.

Hagoort P, Indefrey P. 2014. The neurobiology of language beyond single words. Ann Rev Neurosci, 37:347–362.

Hasson U, Chen J, Honey CJ. 2015. Hierarchical process memory: memory as an integral component of information processing. Trends Cogn Sci. 19(6):304–313.

Herholz SC, Zatorre RJ. 2012. Musical training as a framework for brain plasticity: behavior, function, and structure. Neuron. 76(3):486–502.

Herrmann B, Obleser J, Kalberlah C, Haynes JD, Friederici AD. 2012. Dissociable neural imprints of perception and grammar in auditory functional imaging. Hum Brain Mapp. 33(3):584–595.

Hoch L, Poulin-Charronnat B, Tillmann B. 2011. The influence of task-irrelevant music on language processing: syntactic and semantic structures. Front Psychol. 2:112.

Hrong-Tai Fai A, Cornelius PL. 1996. Approximate F-tests of multiple degree of freedom hypotheses in generalized least squares analyses of unbalanced split-plot experiments. J Stat Comput Simul. 54(4):363–378.

Ivanova A, Srikant S, Sueoka Y, Kean H, Dhamala R, O’Reilly U-M, Bers MU, Fedorenko E. 2020. Comprehension of computer code relies primarily on domain-general executive resources. eLife. 9:e58906.

Ivanova A, Mineroff Z, Zimmerer V, Kanwisher N, Varley R, Fedorenko E. 2021. The language network is recruited but not required for non-verbal semantic processing. bioRxiv.

Jackendoff R. 2002. English particle constructions, the lexicon, and the autonomy of syntax. In Dehé N, Jackendoff R, McIntyre A, Urban S, editors. Verb-particle explorations. Berlin (Germany): De Gruyter. p. 67–94.

Jackendoff R. 2007. A parallel architecture perspective on language processing. Brain Res. 1146:2–22.

Jackendoff R. 2009. Parallels and nonparallels between language and music. Music Percept. 26(3):195–204.

Jackendoff R, Audring J. 2020. The texture of the lexicon: relational morphology and the parallel architecture. Oxford (UK): Oxford University Press.

Janata P. 1995. ERP measures assay the degree of expectancy violation of harmonic contexts in music. J Cogn Neurosci. 7(2):153–164.

Jentschke S, Koelsch S, Sallat S, Friederici AD. 2008. Children with specific language impairment also show impairment of music-syntactic processing. J Cogn Neurosci. 20(11):1940–1951.

Jouravlev O, Zheng D, Balewski Z, Pongos A, Levan Z, Goldin-Meadow S, Fedorenko E. 2019. Speech-accompanying gestures are not processed by the language-processing mechanisms. Neuropsychologia. 132:107132.

Jouravlev O, Kell A, Mineroff Z, Haskins AJ, Ayyash D, Kanwisher N, Fedorenko E. 2020. Reduced language lateralization in autism and the broader autism phenotype as assessed with robust individual-subjects analyses. Autism Res. 13(10):1746–1761.

Just MA, Carpenter PA, Keller TA, Eddy WF, Thulborn KR. 1996. Brain activation modulated by sentence comprehension. Science. 274(5284):114–116.

Kaplan E, Goodglass H, Weintraub S. 2001. Boston Naming Test. 2nd Ed. Philadelphia (PA): Lippincott Williams & Wilkins.

Kay J, Lesser R, Coltheart M. 1992. Psycholinguistic Assessments of Language Processing in Aphasia (PALPA). Hove (UK): Lawrence Erlbaum.

Kell AJ, Yamins DL, Shook EN, Norman-Haignere SV, McDermott JH. 2018. A task-optimized neural network replicates human auditory behavior, predicts brain responses, and reveals a cortical processing hierarchy. Neuron. 98(3):630–644.

Keller TA, Carpenter PA, Just MA. 2001. The neural bases of sentence comprehension: a fMRI examination of syntactic and lexical processing. Cereb Cortex. 11(3):223–237.

Koelsch S, Gunter T, Friederici AD, Schröger E. 2000. Brain indices of music processing: “nonmusicians” are musical. J Cogn Neurosci. 12(3):520–541.

Koelsch S, Gunter TC, Schröger E, Tervaniemi M, Sammler D, Friederici AD. 2001. Differentiating ERAN and MMN: an ERP study. NeuroReport. 12(7):1385–1389.

Koelsch S, Gunter TC, von Cramon DY, Zysset S, Lohmann G, Friederici AD. 2002. Bach speaks: A cortical “language-network” serves the processing of music. Neuroimage. 17(2):956–966.

Koelsch S. 2006. Significance of Broca’s area and ventral premotor cortex for music-syntactic processing. Cortex. 42(4):518–520.

Koelsch S, Jentschke S, Sammler D, Mietchen D. 2007. Untangling syntactic and sensory processing: An ERP study of music perception. Psychophysiology. 44(3):476–490.

Koelsch S, Rohrmeier M, Torrecuso R, Jentschke S. 2013. Processing of hierarchical syntactic structure in music. Proc Natl Acad Sci USA. 110(38):15443–15448.

Kriegeskorte N, Simmons WK, Bellgowan PS, Baker CI. 2009. Circular analysis in systems neuroscience: the dangers of double dipping. Nat Neurosci. 12(5):535.

Krumhansl CL, Keil FC. 1982. Acquisition of the hierarchy of tonal functions in music. Mem Cognit. 10(3):243–251.

Kunert R, Slevc LR. 2015. A Commentary on:“Neural overlap in processing music and speech”. Front Hum Neurosci. 9:330.

Kunert R, Willems RM, Casasanto D, Patel AD, Hagoort P. 2015. Music and language syntax interact in Broca’s area: An fMRI study. PLoS One. 10(11):e0141069.

Kunert R, Willems RM, Hagoort P. 2016. Language influences music harmony perception: effects of shared syntactic integration resources beyond attention. R Soc Open Sci. 3(2):150685.

Kuperberg GR, Holcomb PJ, Sitnikova T, Greve D, Dale AM, Caplan D. 2003. Distinct patterns of neural modulation during the processing of conceptual and syntactic anomalies. J Cogn Neurosci. 15(2):272–293.

Kuznetsova A, Brockhoff PB, Christensen RH. 2017. lmerTest package: tests in linear mixed effects models. J Stat Softw. 82(13):1–26.

LaCroix A, Diaz AF, Rogalsky C. 2015. The relationship between the neural computations for speech and music perception is context-dependent: an activation likelihood estimate study. Front Psychol. 6:1138.

Lartillot O, Grandjean D. 2019. Tempo and metrical analysis by tracking multiple metrical levels using autocorrelation. Appl Sci. 9(23):5121.

Lartillot O, Toiviainen P. 2007. September). A Matlab toolbox for musical feature extraction from audio. In: Proceedings of the 10th International Conference on Digital Audio Effects; 2007 Sep 10-15; Bordeaux, France. p. 244.

Lecours A, Joanette Y. 1980. Linguistic and other psychological aspects of paroxysmal aphasia. Brain Lang. 10(1):1–23.

Lerdahl F, Jackendoff R. 1977. Toward a formal theory of tonal music. J Music Theory. 21(1):111–171.

Lerdahl F, Jackendoff R. 1983. An overview of hierarchical structure in music. Music Percept. 1(2):229–252.

Levin B, Rappaport-Hovav M. 2005. Argument realization. Cambridge (UK): Cambridge University Press.

Levitin DJ, Menon V. 2003. Musical structure is processed in “language” areas of the brain: a possible role for Brodmann Area 47 in temporal coherence. Neuroimage. 20(4):2142–2152.

Linebarger MC, Schwartz MF, Saffran EM. 1983. Sensitivity to grammatical structure in so-called agrammatic aphasics. Cognition. 13(3):361–392.

Lindblom B, Sundberg J. 1969. Towards a generative theory of melody. Speech Transmission Laboratory. Quarterly Progress and Status Reports. 10:53–86.

Lipkin B, Tuckute G, Affourtit J, Small H, Mineroff Z, Jouravlev O, Rakocevic L, Pritchett B, et al. 2022. Probabilistic atlas for the language network based on precision fMRI data from> 800 individuals. Sci Data. 9(1):1–10.

Liu YF, Kim J, Wilson C, Bedny M. 2020. Computer code comprehension shares neural resources with formal logical inference in the fronto-parietal network. Elife. 9:e59340.

Liu J, Hilton CB, Bergelson E, Mehr SA. 2021. Language experience shapes music processing across 40 tonal, pitch-accented, and non-tonal languages. bioRxiv.

Luria AR, Tsvetkova LS, Futer DS. 1965. Aphasia in a composer. J Neurol Sci. 2(3):288–292.

Maess B, Koelsch S, Gunter TC, Friederici AD. 2001. Musical syntax is processed in Broca’s area: an MEG study. Nat Neurosci. 4(5):540–545.

Mahowald K, Fedorenko E. 2016. Reliable individual-level neural markers of high-level language processing: A necessary precursor for relating neural variability to behavioral and genetic variability. Neuroimage. 139:74–93.

Makowski D. 2018. The psycho Package: An Efficient and Publishing-Oriented Workflow for Psychological Science. J Open Source Softw. 3(22):470.

Malik-Moraleda S, Ayyash D, Gallée J, Affourtit J, Hoffmann M, Mineroff Z, Jouravlev O, Fedorenko E. 2022. An investigation across 45 languages and 12 language families reveals a universal language network. Nat Neurosci. 25(8):1014–1019.

Marin OSM. 1982. Neurological Aspects of Music Perception and Performance. New York (NY): Academic Press.

Matchin W, Hickok G. 2020. The cortical organization of syntax. Cereb Cortex. 30(3):1481–1498.

Mehr SA, Krasnow MM. 2017. Parent-offspring conflict and the evolution of infant-directed song. Evol Hum Behav. 38(5):674–684.

Mehr SA, Singh M, Knox D, Ketter DM, Pickens-Jones D, Atwood S, Lucas C, Jacoby N, Egner AA, Hopkins EJ, et al. (2019). Universality and diversity in human song. Science. 366(6468):eaax0868.

Mehr S, Krasnow M, Bryant G, Hagen E. 2020. Origins of music in credible signaling. Behav Brain Sci. 44:e60.

Mesulam MM, Rogalski EJ, Wieneke C, Hurley RS, Geula C, Bigio EH, Thompson CK, Weintraub S. 2014. Primary progressive aphasia and the evolving neurology of the language network. Nat Rev Neurol. 10(10):554.

Mesulam MM, Thompson CK, Weintraub S, Rogalski EJ. 2015. The Wernicke conundrum and the anatomy of language comprehension in primary progressive aphasia. Brain. 138(8):2423–2437.

McDermott J. 2008. The evolution of music. Nature. 453(7193):287–288.

Mineroff Z*, Blank I*, Mahowald K, Fedorenko E. 2018. A robust dissociation among the language, multiple demand, and default mode networks: evidence from inter-region correlations in effect size. Neuropsychologia. 119:501–511.

Mollica F, Shain C, Affourtit J, Kean H, Siegelman M, Fedorenko E. 2020. Another look at the constituent structure of sentences in the human brain [Poster presentation]. SNL 2020; October 21-24; virtual.

Monti MM, Parsons LM, Osherson DN. 2009. The boundaries of language and thought in deductive inference. Proc Natl Acad Sci USA. 106(30):12554–12559.

Monti MM, Parsons LM, Osherson DN. 2012. Thought beyond language: Neural dissociation of algebra and natural language. Psychol Sci. 23(8):914–922.

Morosan P, Rademacher J, Schleicher A, Amunts K, Schormann T, Zilles K. 2001. Human primary auditory cortex: cytoarchitectonic subdivisions and mapping into a spatial reference system. Neuroimage. 13(4):684–701.

Musso M, Weiller C, Horn A, Glauche V, Umarova R, Hennig J, Schneider A, Rijntjes M. 2015. A single dual-stream framework for syntactic computations in music and language. Neuroimage. 117:267–283.

Nettl B. 2015. The study of ethnomusicology: Thirty-three discussions. Champaign (IL): University of Illinois Press.

Newman AJ, Pancheva R, Ozawa K, Neville HJ, Ullman MT. 2001. An event-related fMRI study of syntactic and semantic violations. J Psycholinguist Res. 30(3):339–364.

Ngo MK, Vu KPL, Strybel TZ. 2016. Effects of music and tonal language experience on relative pitch performance. Am J Psychol. 129(2):125–134.

Nieto-Castañon A, Fedorenko E. 2012. Subject-specific functional localizers increase sensitivity and functional resolution of multi-subject analyses. Neuroimage. 63(3):1646–1669.

Norman-Haignere S, Kanwisher N, McDermott JH. 2013. Cortical pitch regions in humans respond primarily to resolved harmonics and are located in specific tonotopic regions of anterior auditory cortex. J Neurosci. 33(50):19451–19469.

Norman-Haignere S, Kanwisher NG, McDermott JH. 2015. Distinct cortical pathways for music and speech revealed by hypothesis-free voxel decomposition. Neuron. 88(6):1281–1296.

Norman-Haignere SV, Feather J, Boebinger D, Brunner P, Ritaccio A, McDermott JH, Schalk G, Kanwisher N. 2022. A neural population selective for song in human auditory cortex. Curr Biol. 32(7):1470–1484.

Oldfield RC. 1971. The assessment and analysis of handedness: the Edinburgh inventory. Neuropsychologia. 9(1):97–113.

Omigie D, Samson S. 2014. A protective effect of musical expertise on cognitive outcome following brain damage?. Neuropsychol Rev. 24(4):445–460.

Overath T, McDermott JH, Zarate JM, Poeppel D. 2015. The cortical analysis of speech-specific temporal structure revealed by responses to sound quilts. Nat Neurosci. 18(6):903–911.

Pallier C, Devauchelle AD, Dehaene S. 2011. Cortical representation of the constituent structure of sentences. Proc Natl Acad Sci USA. 108(6):2522–2527.

Patel AD, Gibson E, Ratner J, Besson M, Holcomb PJ. 1998. Processing syntactic relations in language and music: An event-related potential study. J Cogn Neurosci. 10(6):717–733.

Patel AD. 2003. Language, music, syntax and the brain. Nat Neurosci. 6(7):674–681.

Patel AD. 2008. Music, Language, and the Brain. Oxford (UK): Oxford University Press.

Patel AD, Iversen JR, Wassenaar M, Hagoort P. 2008a. Musical syntactic processing in agrammatic Broca’s aphasia. Aphasiology. 22(7-8):776–789.

Patel AD, Wong M, Foxton J, Lochy A, Peretz I. 2008b. Speech intonation perception deficits in musical tone deafness (congenital amusia). Music Percept. 25(4):357–368.

Patel AD. 2012. Language, music, and the brain: a resource-sharing framework. In: Rebuschat P, Rohrmeier M, Hawkins J, Cross I, editors. Language and Music as Cognitive Systems. Oxford (UK): Oxford University Press. p. 204–223.

Patel AD, Morgan E. 2017. Exploring cognitive relations between prediction in language and music. Cogn Sci. 41:303–320.

Patterson RD, Uppenkamp S, Johnsrude IS, Griffiths TD. 2002. The processing of temporal pitch and melody information in auditory cortex. Neuron. 36(4):767–776.

Paunov A, Blank IA, Fedorenko E. 2019. Functionally distinct language and Theory of Mind networks are synchronized at rest and during language comprehension. J Neurophysiol. 121:1244–1265.

Paunov AM, Blank IA, Jouravlev O, Mineroff Z, Gallée J, Fedorenko E. 2022. Differential tracking of linguistic vs. mental state content in naturalistic stimuli by language and Theory of Mind (ToM) brain networks. Neurobiol Lang. 3(3):413–440.

Peelle JE, Troiani V, Wingfield A, Grossman M. 2010. Neural processing during older adults’ comprehension of spoken sentences: age differences in resource allocation and connectivity. Cereb Cortex. 20(4):773–782.

Penagos H, Melcher JR, Oxenham AJ. 2004. A neural representation of pitch salience in nonprimary human auditory cortex revealed with functional magnetic resonance imaging. J Neurosci. 24(30):6810–6815.

Peretz I. 1990. Processing of local and global musical information by unilateral brain-damaged patients. Brain. 113(4):1185–1205.

Peretz I, Kolinsky R, Tramo M, Labrecque R, Hublet C, Demeurisse G, Belleville S. 1994. Functional dissociations following bilateral lesions of auditory cortex. Brain. 117(6):1283–1301.

Peretz I, Belleville S, Fontaine S. 1997. Dissociations between music and language functions after cerebral resection: a new case of amusia without aphasia. Can J Exp Psychol 51(4):354–368.

Peretz I, Champod AS, Hyde K. 2003. Varieties of musical disorders: the Montreal Battery of Evaluation of Amusia. Ann N Y Acad Sci. 999(1):58–75.

Peretz I, Coltheart M. 2003. Modularity of music processing. Nat Neurosci. 6(7):688–691.

Peretz I, Vuvan D, Lagrois MÉ, Armony JL. 2015. Neural overlap in processing music and speech. Philos Trans R Soc Lond B Biol Sci. 370(1664):20140090.

Perrachione TK, Fedorenko EG, Vinke L, Gibson E, Dilley LC. 2013. Evidence for shared cognitive processing of pitch in music and language. PLoS One. 8(8):e73372.

Perruchet P, Poulin-Charronnat B. 2013. Challenging prior evidence for a shared syntactic processor for language and music. Psychon Bull Rev. 20(2):310–317.

Piccirilli M, Sciarma T, Luzzi S. 2000. Modularity of music: evidence from a case of pure amusia. J Neurol Neurosurg Psychiatry. 69(4):541–545.

Pinker S, Prince A. 1988. On language and connectionism: Analysis of a parallel distributed processing model of language acquisition. Cognition. 28(1-2):73–193.

Pinker S. 1991. Rules of language. Science. 253(5019):530–535.

Pinker S. 1994. The Language Instinct: How the Mind Creates Language, New York (NY): Harper Collins Publishers, Inc.

Pinker S. 1999. Out of the minds of babes. Science. 283(5398):40–41.

Poldrack RA. 2006. Can cognitive processes be inferred from neuroimaging data?. Trends Cogn Sci. 10(2):59–63.

Poldrack RA. 2011. Inferring mental states from neuroimaging data: from reverse inference to large-scale decoding. Neuron. 72(5):692–697.

Polk M, Kertesz A. 1993. Music and language in degenerative disease of the brain. Brain Cogn. 22(1):98–117.

Poulin-Charronnat B, Bigand E, Madurell F, Peereman R. 2005. Musical structure modulates semantic priming in vocal music. Cognition. 94:B67–B78.

Pritchett B, Hoeflin C, Koldewyn K, Dechter E, Fedorenko E. 2018. High-level language processing regions are not engaged in action observation or imitation. J Neurophysiol. 120(5):2555–2570.

Riemann H. 1877. Musikalische Syntaxis: Grundriss einer harmonischen Satzbildungslehre. Leipzig: Breitkopf und Härtel.

Roads C, Wieneke P. 1979. Grammars as representations for music. Comput Music J. 3(1):48–55.

Roberts I. 2012. Comments and a conjecture inspired by Fabb and Halle. In: Rebuschat P, Rohrmeier M, Hawkins JA, Cross I, editors. Language and Music as Cognitive Systems. Oxford: Oxford University Press. p. 51–66.

Röder B, Stock O, Neville H, Bien S, Rösler F. 2002. Brain activation modulated by the comprehension of normal and pseudo-word sentences of different processing demands: a functional magnetic resonance imaging study. Neuroimage. 15(4):1003–1014.

Rogalsky C, Hickok G. 2011. The role of Broca’s area in sentence comprehension. J Cogn Neurosci. 23(7):1664–1680.

Rueffler C, Hermisson J, Wagner GP. 2012. Evolution of functional specialization and division of labor. Proc Natl Acad Sci USA. 109(6):E326–E335.

Sag I, Wasow T, Bender E. 2003. Formal syntax, an introduction. CSLI publication.

Sammler D, Koelsch S, Ball T, Brandt A, Elger CE, Friederici AD, Grigutsch M, Huppertz H-J, Knosche TR, Wellmer J, Widman G, Schulze-Bonhaged A. 2009. Overlap of musical and linguistic syntax processing: intracranial ERP evidence. Ann N Y Acad Sci. 1169(1):494–498.

Sammler D, Koelsch S, Friederici AD. 2011. Are left fronto-temporal brain areas a prerequisite for normal music-syntactic processing?. Cortex. 47(6):659–673.

Sammler D, Koelsch S, Ball T, Brandt A, Grigutsch M, Huppertz HJ, Wellmer J, Widman G, Elger CE, Friederici AD, Schulze-Bonhaged A. 2013. Co-localizing linguistic and musical syntax with intracranial EEG. Neuroimage. 64:134–146.

Savage PE, Loui P, Tarr B, Schachner A, Glowacki L, Mithen S, Fitch WT. 2021. Music as a coevolved system for social bonding. Behav Brain Sci. 44(e59):1–22.

Schmidt S. 2009. Shall We Really Do It Again? The Powerful Concept of Replication Is Neglected in the Social Sciences. Rev Gen Psychol. 13(2):90–100.

Scott TL, Gallée J, Fedorenko E. 2017. A new fun and robust version of an fMRI localizer for the frontotemporal language system. Cogn Neurosci. 8(3):167–176.

Shain C*, Blank I*, Van Shijndel M, Schuler W, Fedorenko E. 2020. fMRI reveals language-specific predictive coding during naturalistic sentence comprehension. Neuropsychologia. 138:107307.

Shain C, Kean H, Lipkin B, Affourtit J, Siegelman M, Mollica F, Fedorenko E. 2021. ‘Constituent length’effects in fMRI do not provide evidence for abstract syntactic processing. bioRxiv.

Shain C, Blank IA, Fedorenko E, Gibson E, Schuler W. 2022. Robust effects of working memory demand during naturalistic language comprehension in language-selective cortex. J Neurosci. 42(39):7412–7430.

Shashidhara S, Mitchell DJ, Erez Y, Duncan J. 2019. Progressive recruitment of the frontoparietal multiple-demand system with increased task complexity, time pressure, and reward. J Cogn Neurosci. 31(11):1617–1630.

Sihvonen AJ, Särkämö T, Leo V, Tervaniemi M, Altenmüller E, Soinila S. 2017. Music-based interventions in neurological rehabilitation. Lancet Neurol. 16(8):648–660.

Slevc LR, Rosenberg JC, Patel AD. 2009. Making psycholinguistics musical: Self-paced reading time evidence for shared processing of linguistic and musical syntax. Psychon Bull Rev. 16(2):374–381.

Slevc LR, Reitman J, Okada B. 2013. Syntax in music and language: the role of cognitive control. In: Proceedings of the Annual Meeting of the Cognitive Science Society; 2013 Jul 31-Aug 3; Berlin, Germany; p. 3414–3419.

Slevc LR, Okada BM. 2015. Processing structure in language and music: a case for shared reliance on cognitive control. Psychon Bull Rev. 22(3):637–652.

Slevc LR, Faroqi-Shah Y, Saxena S, Okada BM. 2016. Preserved processing of musical structure in a person with agrammatic aphasia. Neurocase. 22(6):505–511.

Stromswold K, Caplan D, Alpert N, Rauch S. 1996. Localization of syntactic comprehension by positron emission tomography. Brain Lang. 52(3):452–473.

Sueoka Y, Paunov A, Ivanova A, Blank IA, Fedorenko E. 2022. The language network reliably ‘tracks’ naturalistic meaningful non-verbal stimuli. bioRxiv

Sullivan GM, Feinn R. 2012. Using effect size—or why the P value is not enough. J Grad Med Educ. 4(3):279–282.

Sundberg J, Lindblom B. 1976. Generative theories in language and music descriptions. Cognition. 4(1):99–122.

Swain JP. 1995. The concept of musical syntax. Music Q. 79(2):281–308.

Tahmasebi AM, Davis MH, Wild CJ, Rodd JM, Hakyemez H, Abolmaesumi P, Johnsrude IS. 2012. Is the link between anatomical structure and function equally strong at all cognitive levels of processing?. Cereb Cortex. 22(7):1593–1603.

Tarantola A. 2005. Inverse problem theory and methods for model parameter estimation. Philadelphia (PA): Society for Industrial and Applied Mathematics.

Temperley D. 2022. Music and Language. Annu Rev Linguist. 8:153–170.

te Rietmolen NA, Mercier M, Trebuchon A, Morillon B, Schon D. 2022. Speech and music recruit frequency-specific distributed and overlapping cortical networks. bioRxiv.

Tillmann B, Janata P, Bharucha JJ. 2003. Activation of the inferior frontal cortex in musical priming. Cogn Brain Res. 16(2):145–161.

Tillmann B, Koelsch S, Escoffier N, Bigand E, Lalitte P, Friederici AD, von Cramon DY. 2006. Cognitive priming in sung and instrumental music: activation of inferior frontal cortex. Neuroimage. 31(4):1771–1782.

Tillmann B. 2012. Music and Language Perception: Expectations, Structural Integration, and Cognitive Sequencing. Top Cogn Sci. 4(4):568–584.

Trehub SE. 2003. The developmental origins of musicality. Nat Neurosci. 6(7):669–673.

Tyler LK, Marslen-Wilson WD, Randall B, Wright P, Devereux BJ, Zhuang J, Papoutsi M, Stamatakis EA. 2011. Left inferior frontal cortex and syntax: function, structure and behaviour in patients with left hemisphere damage. Brain. 134(2):415–431.

Van de Cavey J, Hartsuiker RJ. 2016. Is there a domain-general cognitive structuring system? Evidence from structural priming across music, math, action descriptions, and language. Cognition. 146:172–184.

Varley R, Siegal M. 2000. Evidence for cognition without grammar from causal reasoning and ‘theory of mind’ in an agrammatic aphasic patient. Curr Biol. 10(12):723–726.

Varley RA, Klessinger NJ, Romanowski CA, Siegal M. 2005. Agrammatic but numerate. Proc Natl Acad Sci USA. 102(9):3519–3524.

Vázquez-Rodríguez B, Suárez LE, Markello RD, Shafiei G, Paquola C, Hagmann P, van den Heuvel MP, Bernhardt BC, Spreng RN, Misic B. 2019. Gradients of structure–function tethering across neocortex. Proc Natl Acad Sci USA. 116(42):21219–21227.

Vuust P, Heggli OA, Friston KJ, Kringelbach ML. 2022. Music in the brain. Nat Rev Neurosci. 23(5):287–305.

Wehbe L, Blank I, Shain C, Futrell R, Levy R, Malsburg T, Smith N, Gibson E, Fedorenko E. 2021. Incremental language comprehension difficulty predicts activity in the language network but not the multiple demand network. Cereb Cortex. 31(9):4006–4023.

Westfall J, Kenny DA, Judd CM. 2014. Statistical power and optimal design in experiments in which samples of participants respond to samples of stimuli. J Exp Psychol. 143(5):2020–2045.

Willems RM, Van der Haegen L, Fisher SE, Francks C. 2014. On the other hand: including left-handers in cognitive neuroscience and neurogenetics. Nat Rev Neurosci. 15(3):193–201.

Wilson SM, Saygın AP. 2004. Grammaticality judgment in aphasia: Deficits are not specific to syntactic structures, aphasic syndromes, or lesion sites. J Cogn Neurosci. 16(2):238–252.

Wilson SM, Galantucci S, Tartaglia MC, Gorno-Tempini ML. 2012. The neural basis of syntactic deficits in primary progressive aphasia. Brain Lang. 122(3):190–198.

Wong PC, Perrachione TK. 2007. Learning pitch patterns in lexical identification by native English-speaking adults. Appl Psycholinguist. 28(4):565–585.

Woolgar A, Duncan J, Manes F, Fedorenko E. 2018. Fluid intelligence is supported by the multiple-demand system not the language system. Nat Hum Behav. 2(3):200–204.

Zatorre RJ. 1984. Musical perception and cerebral function: A critical review. Music Percept. 2(2):196–221.

