## Supplementary Information for "The human language system, including its inferior frontal component in ‘Broca’s area’, does not support music perception"

#### SI-1. Sanity Check Analyses

Auditory *sentences* > *nonwords* and *sentences* > *foreign* contrasts

| Contrast | Language network | LIFGorb | LIFG | LMFG | LAnt Temp | LPost Temp |
|---|---|---|---|---|---|---|
| sentences >nonwords (Expt 1) | b=0.612<br>se=0.096<br>t=6.397<br>p<0.001*** | b=0.288<br>se=0.236<br>t=1.218<br>p=1.000 | b=0.557<br>se=0.184<br>t=3.035<br>p=0.036* | b=0.722<br>se=0.198<br>t=3.639<br>p=0.010* | b=0.740<br>se=0.106<br>t=6.953<br>p<0.001*** | b=0.754<br>se=0.123<br>t=6.154<br>p<0.001*** |
| sentences >foreign (Expt 4) | b=1.397<br>se=0.133<br>t=10.529<br>p<0.001*** | b=1.134<br>se=0.337<br>t=3.367<br>p=0.017* | b=1.518<br>se=0.247<br>t=6.151<br>p<0.001*** | b=1.723<br>se=0.213<br>t=8.097<br>p<0.001*** | b=1.044<br>se=0.164<br>t=6.384<br>p<0.001*** | b=1.565<br>se=0.207<br>t=7.554<br>p<0.001*** |

**Table SI-1a**. Responses to the auditory *sentences* > *nonwords* and *sentences* > *foreign* contrasts in Experiments 1 and 4. The significance values for the individual ROIs have been FDR-corrected for the number of fROIs (n=5).

Response to the six music conditions in bilateral primary auditory cortex

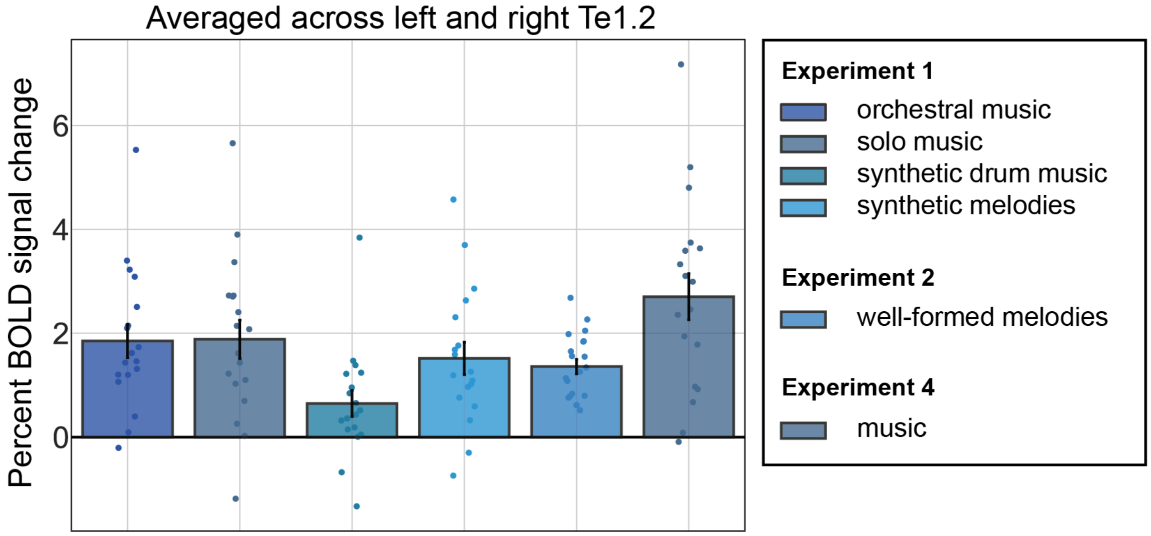


**Figure SI-1**. Responses of the bilateral Te1.2 to the six music conditions in Experiments 1, 2, and 4. All music conditions show reliable above-baseline responses.

| Contrast | Bilateral Te1.2 |
|---|---|
| orchestral music (Expt 1) > fixation | β=1.851 se=0.312 df=18.000 d=3.357 t=5.936 p<0.001*** |
| single-instrument music (Expt 1) > fixation | β=1.884 se=0.360 df=18.000 d=3.417 t=5.235 p<0.001*** |
| synthetic drum music (Expt 1) > fixation | β=0.647 se=0.245 df=18.000 d=1.173 t=2.644 p=0.017* |

| | |
|---|---|
| synthetic melodies (Expt 1) > fixation | β=1.516 se=0.311 df=7.864 d=2.750 t=4.880 p=0.001* |
| well-formed melodies (Expt 2) > fixation | β=1.354 se=0.135 df=20.000 d=2.456 t=10.028 p<0.001*** |
| music (Expt 3) > fixation | β=2.704 se=0.575 df=5.793 d=4.904 t=4.705 p=0.004* |

**Table SI-1b**. Responses to the music conditions relative to the fixation baseline in bilateral Te1.2. The significance values for the individual ROIs have been FDR-corrected for the number of fROIs (n=5).

### SI-2. Critical Analyses in language fROIs defined by an auditory contrast (Experiments 1 and 4)

We performed the same set of critical analyses in language fROIs defined using auditory *sentences* > *nonwords* in English (Experiment 1) and *Mandarin sentences* > *foreign* (Experiment 4). Similar to the approach described in the main text for the definition of language fROIs based on the visual *sentences* > *nonwords* contrast, an across-runs cross-validation procedure was used to ensure independence between the data used to define the fROIs and to estimate their response magnitudes. The results are consistent with the results from the visual *sentences* > *nonwords* language fROIs: 1) responses to music fall at or below baseline, are not higher than responses elicited by nonwords, and do not differ from other non-linguistic, non-music conditions, and songs do not elicit a stronger response than lyrics; 2) responses to synthetic melodies and synthetic drum music do not significantly differ from their scrambled counterparts; and 3) for Mandarin native speakers, although the response to music is above baseline at the network level, the responses do not significantly differ from nonwords and environmental sounds.

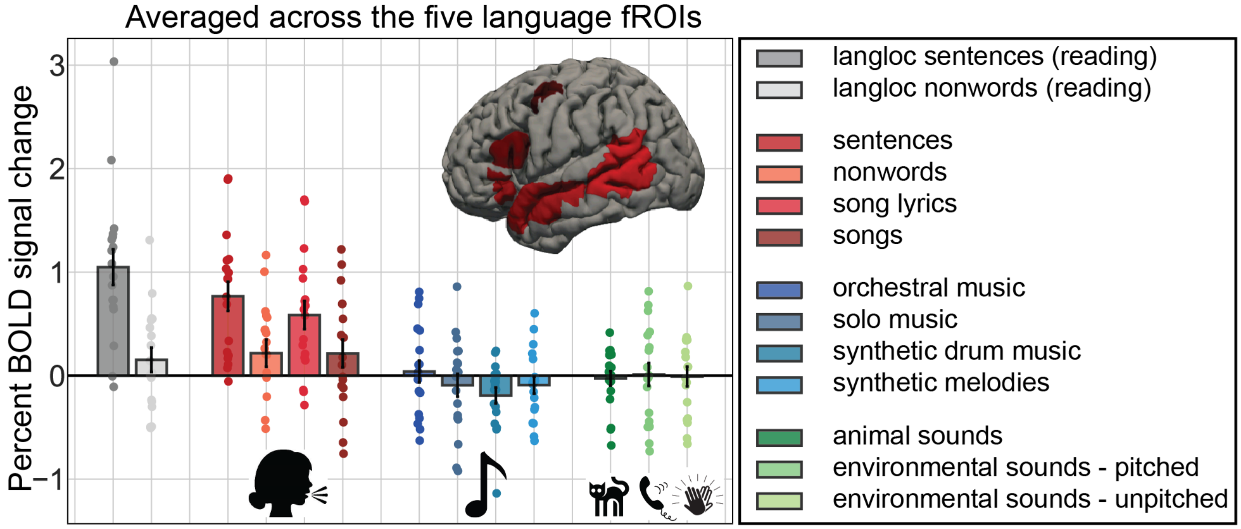


**Figure SI-2a**. Responses of the language fROIs (defined by auditory *sentences* > *nonwords*) to the language localizer conditions (in grey), to the four auditory conditions containing linguistic information in (red shades), to the four music conditions (blue shades), and to the three non-linguistic/non-music auditory conditions (green shades). For the language localizer results, we include here participants in Experiments 1 and 2. The responses to the music conditions cluster around the fixation baseline, are much lower than the responses to sentences, and not higher than the responses to non-music sounds.

| Contrast | Language network | LIFGorb | LIFG | LMFG | LAnt Temp | LPost Temp |
|---|---|---|---|---|---|---|
| **music > fixation** | | | | | | |
| orchestral music >fixation | β=0.040 se=0.103 df=18.000 d=0.076 t=0.394 p=0.699 | β=0.125 se=0.170 df=nan d=nan t=0.737 p=1.000 | β=0.090 se=0.121 df=nan d=nan t=0.746 p=1.000 | β=-0.017 se=0.141 df=nan d=nan t=-0.124 p=1.000 | β=-0.001 se=0.096 df=nan d=nan t=-0.008 p=1.000 | β=0.004 se=0.094 df=nan d=nan t=0.046 p=1.000 |
| single-instrument music >fixation | β=-0.092 se=0.109 df=17.265 d=-0.166 t=-0.846 p=0.409 | β=-0.132 se=0.150 df=nan d=nan t=-0.883 p=1.000 | β=0.033 se=0.161 df=nan d=nan t=0.208 p=1.000 | β=-0.031 se=0.152 df=nan d=nan t=-0.204 p=1.000 | β=-0.218 se=0.091 df=nan d=nan t=-2.389 p=0.145 | β=-0.112 se=0.091 df=nan d=nan t=-1.229 p=1.000 |
| drum music >fixation | β=-0.192 se=0.075 df=18.000 d=-0.437 t=-2.561 p=0.020* | β=-0.181 se=0.134 df=nan d=nan t=-1.345 p=0.980 | β=-0.215 se=0.142 df=nan d=nan t=-1.511 p=0.745 | β=-0.209 se=0.100 df=nan d=nan t=-2.103 p=0.255 | β=-0.224 se=0.075 df=nan d=nan t=-2.995 p=0.040* | β=-0.132 se=0.052 df=nan d=nan t=-2.512 p=0.110 |
| synthetic melodies >fixation | β=-0.092 se=0.079 df=18.000 d=-0.199 t=-1.160 p=0.261 | β=-0.052 se=0.123 df=nan d=nan t=-0.426 p=1.000 | β=-0.056 se=0.124 df=nan d=nan t=-0.453 p=1.000 | β=-0.028 se=0.143 df=nan d=nan t=-0.196 p=1.000 | β=-0.172 se=0.069 df=nan d=nan t=-2.493 p=0.115 | β=-0.150 se=0.078 df=nan d=nan t=-1.933 p=0.350 |
| **music > nonwords** | | | | | | |
| orchestral music >nonwords | β=-0.176 se=0.071 df=157.734 d=-0.289 t=-2.499 p=0.013* | β=-0.224 se=0.192 df=18.000 d=-0.326 t=-1.165 p=1.000 | β=-0.038 se=0.169 df=18.000 d=-0.059 t=-0.227 p=1.000 | β=-0.006 se=0.170 df=18.000 d=-0.010 t=-0.036 p=1.000 | β=-0.436 se=0.126 df=18.000 d=-0.911 t=-3.452 p=0.015* | β=-0.177 se=0.162 df=18.000 d=-0.339 t=-1.092 p=1.000 |
| single-instrument music >nonwords | β=-0.309 se=0.077 df=162.000 d=-0.499 t=-4.025 p<0.001*** | β=-0.482 se=0.210 df=18.000 d=-0.745 t=-2.289 p=0.170 | β=-0.095 se=0.209 df=18.000 d=-0.132 t=-0.456 p=1.000 | β=-0.020 se=0.171 df=18.000 d=-0.031 t=-0.115 p=1.000 | β=-0.654 se=0.157 df=36.000 d=-1.388 t=-4.165 p<0.001*** | β=-0.294 se=0.161 df=18.000 d=-0.569 t=-1.829 p=0.420 |
| synthetic drum music >nonwords | β=-0.409 se=0.070 df=157.653 d=-0.714 t=-5.877 p<0.001*** | β=-0.530 se=0.166 df=18.000 d=-0.859 t=-3.183 p=0.025* | β=-0.343 se=0.202 df=18.000 d=-0.502 t=-1.700 p=0.530 | β=-0.198 se=0.167 df=18.000 d=-0.360 t=-1.188 p=1.000 | β=-0.659 se=0.137 df=18.000 d=-1.481 t=-4.828 p<0.001*** | β=-0.313 se=0.145 df=18.000 d=-0.667 t=-2.165 p=0.220 |
| synthetic melodies >nonwords | β=-0.308 se=0.071 df=161.998 d=-0.531 | β=-0.402 se=0.168 df=18.000 d=-0.674 | β=-0.185 se=0.180 df=18.000 d=-0.284 | β=-0.017 se=0.183 df=18.000 d=-0.027 | β=-0.608 se=0.110 df=18.000 d=-1.391 | β=-0.332 se=0.156 df=18.000 d=-0.667 |

| | | | | | | |
|---|---|---|---|---|---|---|
| | t=-4.318<br>p<0.001*** | t=-2.387<br>p=0.140 | t=-1.029<br>p=1.000 | t=-0.091<br>p=1.000 | t=-5.520<br>p<0.001*** | t=-2.124<br>p=0.240 |
| **music > non-linguistic, non-music condition** | | | | | | |
| music (combined) >animal sounds | β=-0.057<br>se=0.053<br>df=427.674<br>d=-0.116<br>t=-1.085<br>p=0.279 | β=-0.215<br>se=0.152<br>df=72.000<br>d=-0.370<br>t=-1.416<br>p=0.805 | β=-0.125<br>se=0.131<br>df=72.000<br>d=-0.225<br>t=-0.954<br>p=1.000 | β=-0.021<br>se=0.134<br>df=72.000<br>d=-0.039 t<br>=-0.161<br>p=1.000 | β=0.019<br>se=0.078<br>df=72.000<br>d=0.054<br>t=0.237<br>p=1.000 | β=0.058<br>se=0.074<br>df=72.000<br>d=0.170<br>t=0.779<br>p=1.000 |
| music (combined) >environmental (pitched) | β=-0.095<br>se=0.054<br>df=427.756<br>d=-0.184<br>t=-1.776<br>p=0.076 | β=-0.111<br>se=0.142<br>df=72.000<br>d=-0.184<br>t=-0.780<br>p=1.000 | β=-0.221<br>se=0.135<br>df=72.000<br>d=-0.370<br>t=-1.636<br>p=0.530 | β=-0.093<br>se=0.144<br>df=72.000<br>d=-0.159 t=-<br>0.646<br>p=1.000 | β=0.038<br>se=0.079<br>df=72.000<br>d=0.103<br>t=0.485<br>p=1.000 | β=-0.091<br>se=0.075<br>df=72.000<br>d=-0.266<br>t=-1.210<br>p=1.000 |
| music (combined) >environmental (unpitched) | β=-0.075<br>se=0.056<br>df=432.000<br>d=-0.148<br>t=-1.345<br>p=0.179 | β=-0.121<br>se=0.160<br>df=72.000<br>d=-0.195<br>t=-0.754<br>p=1.000 | β=-0.067<br>se=0.142<br>df=72.000<br>d=-0.114<br>t=-0.474<br>p=1.000 | β=-0.089<br>se=0.143<br>df=90.000<br>d=-0.164<br>t=-0.622<br>p=1.000 | β=-0.036<br>se=0.080<br>df=72.000<br>d=-0.105<br>t=-0.445<br>p=1.000 | β=-0.065<br>se=0.083<br>df=72.000<br>d=-0.182<br>t=-0.774<br>p=1.000 |
| **(melodic + linguistic content) > linguistic content** | | | | | | |
| songs >lyrics | β=-0.370<br>se=0.090<br>df=157.753<br>d=-0.503<br>t=-4.114<br>p<0.001*** | β=-0.603<br>se=0.294<br>df=18.000<br>d=-0.666<br>t=-2.050<br>p=0.275 | β=-0.413<br>se=0.220<br>df=18.000<br>d=-0.616<br>t=-1.876<br>p=0.385 | β=-0.210<br>se=0.183<br>df=18.000<br>d=-0.300<br>t=-1.150<br>p=1.000 | β=-0.235<br>se=0.127<br>df=18.000<br>d=-0.491<br>t=-1.847<br>p=0.405 | β=-0.392<br>se=0.153<br>df=18.000<br>d=-0.588<br>t=-2.563<br>p=0.100 |

**Table SI-2a**. Statistical results for the contrasts between the music conditions and fixation, nonwords, animal sounds, and environmental sounds, and to the contrast between songs and lyrics in Experiment 1. The significance values for the individual ROIs have been FDR-corrected for the number of fROIs (n=5).

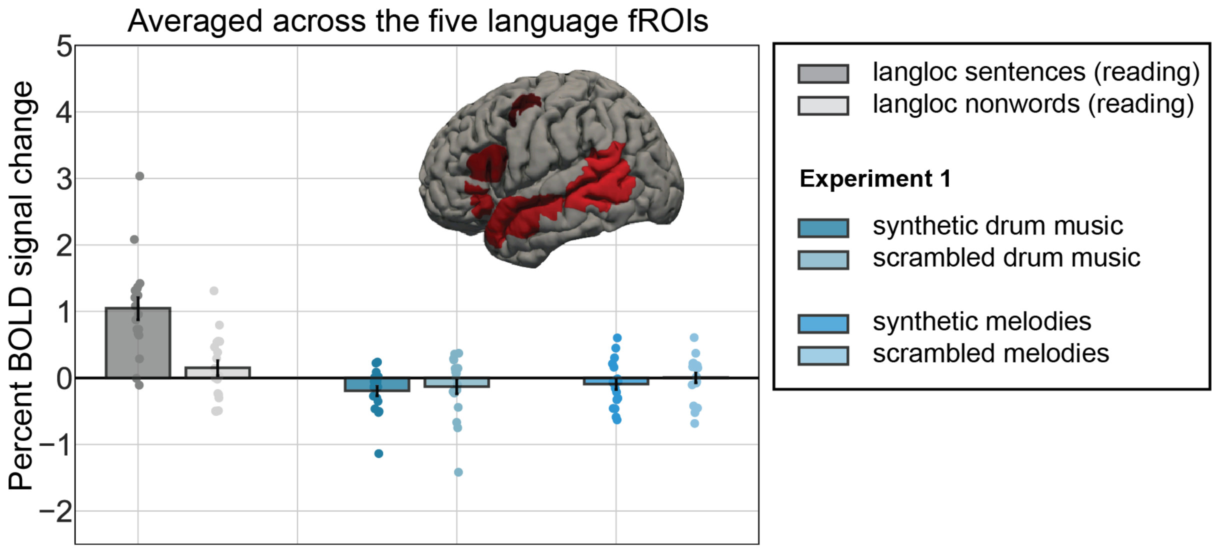


**Figure SI-2b**. Responses of the language fROIs (defined by auditory *sentences* > *nonwords*) to the language localizer conditions (in grey), and to the two sets of conditions targeting structure in music (in blue) from Experiment 1. The responses to the music conditions cluster around the fixation baseline, are much lower than the response to sentences. Neither of the two critical elicits reliable effect.

| **Contrast** | **Language network** | **LIFGorb** | **LIFG** | **LMFG** | **LAntTemp** | **LPostTemp** |
|---|---|---|---|---|---|---|
| synthetic drum music >scrambled drum music | β=-0.061<br>se=0.061<br>df=162.000<br>d=-0.120<br>t=-1.008<br>p=0.315 | β=0.080<br>se=0.178<br>df=18.000<br>d=0.104<br>t=0.451<br>p=1.000 | β=-0.112<br>se=0.140<br>df=18.000<br>d=-0.194<br>t=-0.798<br>p=1.000 | β=-0.227<br>se=0.123<br>df=18.000<br>d=-0.531<br>t=-1.851<br>p=0.405 | β=0.010<br>se=0.085<br>df=18.000<br>d=0.030<br>t=0.121<br>p=1.000 | β=-0.057<br>se=0.064<br>df=18.000<br>d=-0.235<br>t=-0.892<br>p=1.000 |
| synthetic melodies >scrambled synthetic melodies | β=-0.099<br>se=0.058<br>df=162.000<br>d=-0.219<br>t=-1.698<br>p=0.091 | β=-0.176<br>se=0.151<br>df=18.000<br>d=-0.351<br>t=-1.169<br>p=1.000 | β=-0.009<br>se=0.133<br>df=18.000<br>d=-0.018<br>t=-0.070<br>p=1.000 | β=-0.084<br>se=0.171<br>df=18.000<br>d=-0.141<br>t=-0.491<br>p=1.000 | β=-0.126<br>se=0.064<br>df=18.000<br>d=-0.494<br>t=-1.955<br>p=0.330 | β=-0.097<br>se=0.096<br>df=36.000<br>d=-0.337<br>t=-1.005<br>p=1.000 |

**Table SI-2b**. Statistical results for the contrasts between the synthetic drum music and scrambled drum music, and synthetic melodies and scrambled melodies in Experiments 1. The significance values for the individual ROIs have been FDR-corrected for the number of fROIs (n=5).

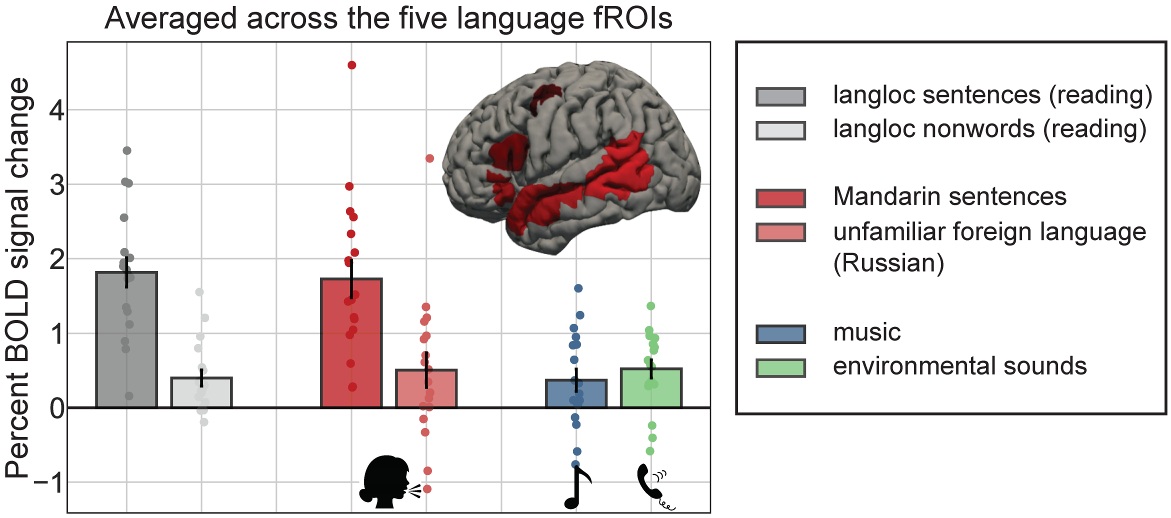


**Figure SI-2c.** Responses of the language fROIs (defined by *Mandarin sentences > foreign*) to the language localizer conditions (in grey), to the language localizer conditions (in grey), to the two auditory conditions containing speech (red shades), to the music condition (blue), and to the non-linguistic/non-music auditory condition (green) in Experiment 4. The response to the music condition is much lower than the responses to sentences, and is not higher than the response to foreign language and environmental sounds.

| **Contrast** | **Language network** | **LIFGorb** | **LIFG** | **LMFG** | **LAntTemp** | **LPostTemp** |
|---|---|---|---|---|---|---|

| | | | | | | |
|---|---|---|---|---|---|---|
| music >fixation | β=0.370 se=0.145 df=18.000 d=0.476 t=2.550 p=0.020* | β=0.437 se=0.216 df=nan d=nan t=2.022 p=0.295 | β=0.485 se=0.166 df=nan d=nan t=2.930 p=0.045* | β=0.353 se=0.230 df=nan d=nan t=1.534 p=0.715 | β=0.191 se=0.155 df=nan d=nan t=1.231 p=1.000 | β=0.385 se=0.154 df=nan d=nan t=2.510 p=0.115 |
| music >foreign | β=-0.134 se=0.113 df=162.000 d=-0.141 t=-1.185 p=0.238 | β=-0.127 se=0.336 df=18.000 d=-0.100 t=-0.377 p=1.000 | β=-0.097 se=0.293 df=36.000 d=-0.110 t=-0.331 p=1.000 | β=0.112 se=0.295 df=18.000 d=0.107 t=0.380 p=1.000 | β=-0.348 se=0.197 df=18.001 d=-0.529 t=-1.763 p=0.475 | β=-0.212 se=0.178 df=18.000 d=-0.287 t=-1.188 p=1.000 |
| music >environmental sounds | β=-0.152 se=0.092 df=157.708 d=-0.197 t=-1.653 p=0.100 | β=-0.219 se=0.170 df=18.000 d=-0.256 t=-1.286 p=1.000 | β=-0.265 se=0.164 df=18.000 d=-0.379 t=-1.613 p=0.620 | β=0.003 se=0.181 df=18.000 d=0.003 t=0.017 p=1.000 | β=-0.083 se=0.114 df=18.000 d=-0.142 t=-0.726 p=1.000 | β=-0.198 se=0.147 df=18.000 d=-0.303 t=-1.346 p=0.975 |

**Table SI-2c**. Statistical results for the contrasts between the music condition and fixation, foreign language, and environmental sounds in Experiments 4. The significance values for the individual ROIs have been FDR-corrected for the number of fROIs (n=5).

### SI-3. Critical analyses with LH fROIs regardless of language network lateralization and excluding RH lateralized subject (Experiment 4)

#### SI-3a. LH fROIs regardless of language network lateralization

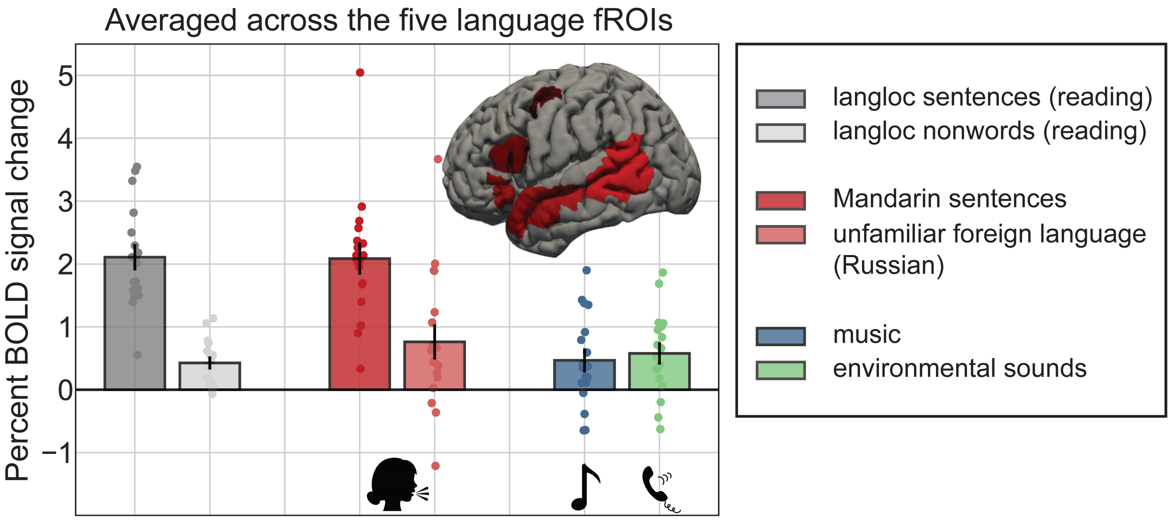


**Figure SI-3a.** Responses of the language fROIs (with all LH fROIs used, including the right-lateralized subject) to the language localizer conditions (in grey), to the two auditory conditions containing speech (red shades), to the music condition (blue), and to the non-linguistic/non-music auditory condition (green) in Experiment 4. The response to the music condition is much lower than the responses to sentences, and is not higher than the response to foreign language and environmental sounds.

| Contrast | Language network | LIFGorb | LIFG | LMFG | LAntTemp | LPostTemp |
|---|---|---|---|---|---|---|
| music >fixation | β=0.466<br>se=0.179<br>df=16.982<br>d=0.518<br>t=2.605<br>p=0.019* | β=0.211<br>se=0.254<br>df=nan<br>d=nan<br>t=0.831<br>p=1.000 | β=0.787<br>se=0.210<br>df=nan<br>d=nan<br>t=3.753<br>p=0.010* | β=0.513<br>se=0.262<br>df=nan<br>d=nan<br>t=1.956<br>p=0.335 | β=0.339<br>se=0.152<br>df=nan<br>d=nan<br>t=2.231<br>p=0.195 | β=0.481<br>se=0.157<br>df=nan<br>d=nan t=3.062<br>p=0.035* |
| music >foreign | β=-0.295<br>se=0.126<br>df=157.697<br>d=-0.277 t=-<br>2.341<br>p=0.020* | β=-0.300<br>se=0.367<br>df=18.000<br>d=-0.256<br>t=-0.818<br>p=1.000 | β=0.161<br>se=0.295<br>df=18.000<br>d=0.163<br>t=0.546<br>p=1.000 | β=-0.075<br>se=0.390<br>df=18.000<br>d=-0.059<br>t=-0.193<br>p=1.000 | β=-0.676<br>se=0.222<br>df=18.000<br>d=-0.810<br>t=-3.052<br>p=0.035* | β=-0.585<br>se=0.224<br>df=18.000<br>d=-0.662<br>t=-2.616<br>p=0.090 |
| music >environmental sounds | β=-0.111<br>se=0.102<br>df=157.789<br>d=-0.125 t=-<br>1.089<br>p=0.278 | β=-0.284<br>se=0.209<br>df=18.000<br>d=-0.310<br>t=-1.359<br>p=0.955 | β=-0.239<br>se=0.192<br>df=18.000<br>d=-0.290<br>t=-1.244<br>p=1.000 | β=0.222<br>se=0.206<br>df=18.000<br>d=0.199<br>t=1.076<br>p=1.000 | β=-0.047<br>se=0.149<br>df=18.000<br>d=-0.075<br>t=-0.312<br>p=1.000 | β=-0.207<br>se=0.178<br>df=18.000<br>d=-0.297<br>t=-1.161<br>p=1.000 |

**Table SI-3a**. Statistical results for the contrasts between the music condition and fixation, foreign language, and environmental sounds in Experiments 4. The significance values for the individual ROIs have been FDR-corrected for the number of fROIs (n=5).

**SI-3b. Excluding subject with RH language lateralization**

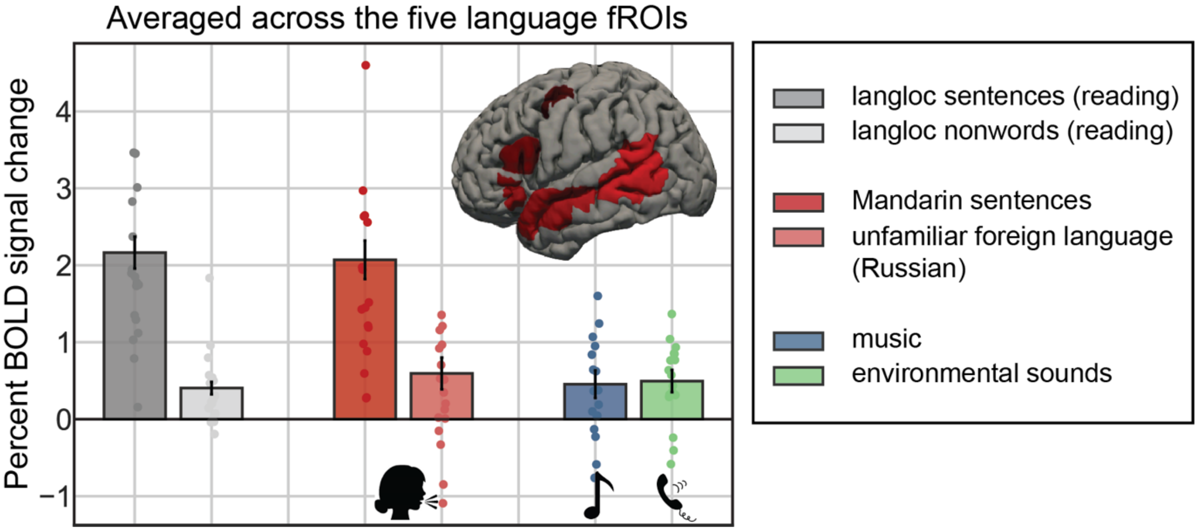


**Figure SI-3b.** Responses of the language fROIs (with all LH fROIs used, including the right-lateralized subject) to the language localizer conditions (in grey), to the two auditory conditions containing speech (red shades), to the music condition (blue), and to the non-linguistic/non-music auditory condition (green) in Experiment 4. The response to the music condition is much lower than the responses to sentences, and is not higher than the response to foreign language and environmental sounds.

| Contrast | Language network | LIFGorb | LIFG | LMFG | LAntTemp | LPostTemp |
|---|---|---|---|---|---|---|

| | | | | | | |
|---|---|---|---|---|---|---|
| music >fixation | β=0.472<br>se=0.180<br>df=16.648<br>d=0.535<br>t=2.624<br>p=0.018* | β=0.324<br>se=0.241<br>df=nan<br>d=nan<br>t=1.347<br>p=0.985 | β=0.750<br>se=0.219<br>df=nan<br>d=nan<br>t=3.426<br>p=0.015* | β=0.477<br>se=0.276<br>df=nan<br>d=nan<br>t=1.732<br>p=0.510 | β=0.347<br>se=0.161<br>df=nan<br>d=nan<br>t=2.154<br>p=0.235 | β=0.461<br>se=0.165<br>df=nan<br>d=nan<br>t=2.788<br>p=0.065 |
| music >foreign | β=-0.119<br>se=0.114<br>df=148.718<br>d=-0.126<br>t=-1.041<br>p=0.299 | β=0.013<br>se=0.217<br>df=17.000<br>d=0.013<br>t=0.058<br>p=1.000 | β=0.262<br>se=0.295<br>df=17.000<br>d=0.285<br>t=0.890<br>p=1.000 | β=0.118<br>se=0.362<br>df=17.000<br>d=0.106<br>t=0.327<br>p=1.000 | β=-0.519<br>se=0.170<br>df=17.000<br>d=-0.709<br>t=-3.056<br>p=0.035* | β=-0.467<br>se=0.203<br>df=17.000<br>d=-0.579<br>t=-2.298<br>p=0.175 |
| music >environmental sounds | β=-0.030<br>se=0.099<br>df=148.792<br>d=-0.035<br>t=-0.297<br>p=0.767 | β=-0.137<br>se=0.161<br>df=17.000<br>d=-0.157<br>t=-0.848<br>p=1.000 | β=-0.171<br>se=0.191<br>df=17.000<br>d=-0.219<br>t=-0.894<br>p=1.000 | β=0.274<br>se=0.212<br>df=17.000<br>d=0.247<br>t=1.290<br>p=1.000 | β=0.021<br>se=0.142<br>df=17.000<br>d=0.034<br>t=0.145<br>p=1.000 | β=-0.134<br>se=0.173<br>df=17.000<br>d=-0.203<br>t=-0.776<br>p=1.000 |

**Table SI-3b**. Statistical results for the contrasts between the music condition and fixation, foreign language, and environmental sounds in Experiments 4. The significance values for the individual ROIs have been FDR-corrected for the number of fROIs (n=5).

### SI-4. Information on the music pieces in Experiment 1

| **Original Piece** | **Composer** |
|---|---|
| Anvil Chorus (From 'Il Trovatore') | Jerry Gray<br>Originally by Giuseppe Verdi |
| Apple Honey | Woody Herman |
| Central Services/ The Office | Michael Kamen |
| Death of Falstaff | William Walton |
| Divertimento in D Major, K. 136 "Salzburg Symphony No. 1": I. Allegro | Wolfgang Amadeus Mozart |
| General Lee’s Solitude | Randy Edelman |
| I Remember Clifford | Benny Golson |
| Just You & I | |
| South Rampart Street Parade | Bob Haggart, Ray Bauduc |
| Symphony No. 5 in E-flat major, Op. 82 | Jean Sibelius |
| Symphony No. 7 in D minor, Op. 70, B. 141 | Antonín Dvořák |

**Table SI-4a**. Original piece and composer of the orchestral music pieces in Experiment 1

| **Instrument** | **Original Piece** | **Composer** |
|---|---|---|
| cello | Suite No. 3 in C major, BWV 1009: VI. Gigue | Johann Sebastian Bach |
| flute | Partita in A minor for solo flute, BWV 1013: IV. Bourree Anglaise | Johann Sebastian Bach |
| guitar | Blue In Green | Bill Evans, Miles Davis |
| guitar | E Is For Emmett | Richard Hyman |
| guitar | In A Mellow Tone | Duke Ellington |

| | | |
|---|---|---|
| guitar | Isn't It A Pity? | George Gershwin, Ira Gershwin |
| piano | Piano Sonata No. 13 in E flat major, Op.27, No.1, "Quasi una fantasia": III. Adagio con espressione | Ludwig van Beethoven |
| piano | Necturne No. 13 In C Minor, Op.48 No.1 | Frédéric Chopin |
| piano | Cubano Chant | Ray Bryant |
| piano | These Foolish Things (Remind Me Of You) | Jack Strachey |
| saxophone | Body and Soul | Johnny Green |
| violin | Partita No. 2 In D Minor, BWV 1004: Giga | Johann Sebastian Bach |

**Table SI-4b**. Original piece and composer of the solo music pieces in Experiment 1

### SI-5. Acoustic properties of stimuli in Experiment 1

| Condition | Duration | Average Pitch | Metre Strength | Tempo | Key Strength | Inharmonicity | Roughness | Irregularity |
|---|---|---|---|---|---|---|---|---|
| Sentences | 9.00 (0.00) | 920.76 (702.54) | 0.38 (0.23) | 129.20 (8.31) | 0.79 (0.06) | 0.45 (0.02) | 26381.51 (11590.47) | 0.67 (0.27) |
| Nonwords | 9.00 (0.00) | 1229.46 (714.77) | 0.31 (0.15) | 129.24 (8.29) | 0.63 (0.11) | 0.52 (0.07) | 18603.92 (10453.71) | 0.46 (0.23) |
| Song lyrics | 9.07 (0.06) | 1526.66 (333.31) | 0.22 (0.09) | 121.92 (6.48) | 0.39 (0.10) | 0.53 (0.02) | 55805.04 (15468.19) | 0.75 (0.17) |
| Songs | 9.00 (0.00) | 1167.79 (224.77) | 0.19 (0.05) | 118.06 (10.84) | 0.39 (0.13) | 0.57 (0.02) | 14971.92 (3980.05) | 0.21 (0.06) |
| Orchestra | 9.00 (0.00) | 1556.95 (597.06) | 0.20 (0.08) | 124.79 (5.28) | 0.52 (0.14) | 0.56 (0.04) | 4349.62 (3448.09) | 0.33 (0.11) |
| Solo | 9.00 (0.00) | 1127.60 (189.53) | 0.21 (0.10) | 124.51 (11.57) | 0.45 (0.10) | 0.56 (0.02) | 30192.75 (5331.37) | 0.19 (0.06) |
| Song | 9.00 (0.00) | 968.37 (555.92) | 0.63 (0.17) | 122.42 (22.14) | 0.26 (0.06) | 0.45 (0.06) | 14001.28 (14098.45) | 0.32 (0.17) |
| Synthetic drum music | 9.00 (0.00) | 1348.72 (647.97) | 0.24 (0.08) | 120.70 (9.84) | 0.28 (0.03) | 0.42 (0.06) | 13286.92 (12816.13) | 0.28 (0.20) |
| Scrambled drum music | 9.00 (0.00) | 930.45 (780.87) | 0.60 (0.20) | 129.81 (9.59) | 0.87 (0.06) | 0.48 (0.05) | 7423.12 (2150.37) | 0.20 (0.07) |
| Synthetic melodies | 9.00 (0.00) | 819.51 (782.99) | 0.25 (0.10) | 130.58 (7.60) | 0.58 (0.14) | 0.49 (0.04) | 7291.55 (2267.12) | 0.25 (0.09) |
| Scrambled melodies | 9.00 (0.00) | 1512.08 (436.00) | 0.27 (0.14) | 118.85 (10.53) | 0.42 (0.12) | 0.50 (0.13) | 29919.49 (41335.58) | 0.67 (0.35) |
| Animal sounds | 9.00 (0.00) | 1311.10 (704.19) | 0.35 (0.22) | 123.95 (11.54) | 0.50 (0.14) | 0.47 (0.16) | 37557.42 (36258.69) | 1.27 (0.54) |
| Environmental sounds (pitched) | 9.00 (0.00) | 1938.94 (180.06) | 0.21 (0.08) | 121.26 (13.19) | 0.31 (0.12) | 0.51 (0.04) | 122197.95 (81563.04) | 0.10 (0.07) |
| Environmental sounds (unpitched) | 9.00 (0.00) | 920.76 (702.54) | 0.38 (0.23) | 129.20 (8.31) | 0.79 (0.06) | 0.45 (0.02) | 26381.51 (11590.47) | 0.67 (0.27) |

**Table SI-5**. Mean and standard deviations of acoustic properties of each condition in Experiment 1.
